# A ubiquitous *Streptomyces* biosynthetic megacluster encodes an arsenal of synergistic biotin-targeting antibiotics

**DOI:** 10.1101/2025.10.09.681295

**Authors:** R. Gordzevich, M. Xu, W. Wang, M.A. Cook, D. Hackenberger, J.P. Deisinger, M.M. Tu, L.A. Carfrae, M. George, K. Rachwalski, K. Koteva, D. Sychantha, A. Wei, G.D. Wright, E.D. Brown

## Abstract

The rise of multidrug-resistant pathogens underscores the urgent need for antibiotics that act through new targets and mechanisms. Biotin metabolism, essential in most bacteria, remains underexploited therapeutically. Here, we uncover a highly conserved, co-located biosynthetic megacluster in *Streptomyces*, a striking “cluster of clusters”, that encodes four distinct natural product families: acidomycin, stravidins, dapamycins, and α-methyl-KAPA, and is flanked by genes that encode streptavidin, a high-affinity biotin-binding protein. Remarkably, all molecules target different steps in bacterial biotin metabolism, revealing a multi-pronged natural strategy for biotin starvation. This arrangement of four functionally convergent biosynthetic gene clusters at a single genomic locus is without precedent. Even more surprisingly, we find that this anti-biotin megacluster is widespread across *Streptomyces* bacteria, suggesting a deeply conserved evolutionary solution to microbial competition. Mechanistically, the compounds inhibit biotin biosynthesis through enzyme blockade, prodrug activation, covalent cofactor mimicry, and biotin sequestration via co-expressed streptavidin. Stravidin S2 and α-methyl-KAPA are effective in a murine model of multidrug-resistant *E. coli* infection. These findings expose a coordinated biosynthetic logic in microbial secondary metabolites and point to higher-order biosynthetic architectures as promising reservoirs of antibiotic innovation.

## Introduction

The rise of antimicrobial resistance is rapidly eroding the efficacy of clinically used antibiotics, rendering many bacterial infections untreatable^1^. Most antibiotics currently in use are derived from natural products (NPs) produced by environmental microbes, particularly *Streptomyces* spp.^2^. Despite their success, few new NP scaffolds have entered clinical practice in recent decades, mainly due to the rediscovery of known chemotypes. Advances in genome sequencing and NP biosynthetic pathway annotation have reinvigorated NP discovery by enabling genome mining, an approach that leverages biosynthetic gene clusters (BGCs) to predict and prioritize new chemical diversity^3^.

To date, genome mining has primarily focused on identifying single bioactive molecules^4–8^. However, many microbes produce ensembles of secondary metabolites that act in concert, and recent studies suggest that certain BGCs co-localize to form superclusters, generally comprised of two BGCs ^9–12^. These linked loci can encode chemically distinct, co-regulated compounds with synergistic activity. Despite their potential, only a handful of such synergistic NPs have been experimentally characterized.

Here, we describe a conserved and uniquely complex BGC architecture in *Streptomyces* spp., a megacluster that encodes four structurally and mechanistically distinct classes of biotin-targeting compounds—stravidins, acidomycin, dapamycins, and α-methyl-KAPA (α-Me-KAPA)—flanked by genes encoding the biotin-sequestering protein streptavidin, adding a fifth anti-biotin element. Remarkably, this megacluster is more prevalent than BGCs that produce well-known *Streptomyces* antibiotics, such as streptomycin or tetracycline. We demonstrate that each compound encoded in the megacluster inhibits a different stage of the biotin biosynthesis pathway and that their combinations result in potent antibacterial activity against Gram-negative and mycobacterial species. Stravidins (prodrugs of amiclenomycin) and acidomycin, established inhibitors of BioA and BioB, respectively, are shown here for the first time to be co-produced by a single organism^13–16^. Mechanistic studies reveal that dapamycin B acts as a prodrug that releases a non-proteinogenic amino acid upon uptake and is hypothesized to target BioD, while α-Me-KAPA hijacks four biosynthetic enzymes to form α-Me-biotin, which is conjugated to the essential biotin-accepting domain of acetyl-CoA carboxylase, leading to cell death. Furthermore, all four compounds display potent pair-wise synergistic interactions. Stravidin S2 and α-Me-KAPA also exhibit therapeutic promise, significantly reducing the bacterial load of a multi-drug resistant *E. coli* in a murine model of infection.

Our findings reveal a naturally encoded chemical arsenal of unprecedented complexity that disables a conserved bacterial pathway through coordinated, multi-target inhibition. This work further supports the emerging view that synergy is an evolved feature of natural product biosynthesis and highlights the power of genome mining to uncover multi-component systems rather than isolated compounds^17, 18^.

## Results

### Identification of a conserved megacluster in *Streptomyces*

Our longstanding interest in biotin metabolism as an antibiotic target prompted an examination of the BGC responsible for stravidin production^19–22^. We found that the stravidin BGC (*svn* BGC) is co-localized with genes encoding polyketide synthases (PKSs), non-ribosomal peptide synthases (NRPSs), and other enzymes typically associated with natural product biosynthesis in the genomes of *Streptomyces* sp. NRRL S-98 and *Streptomyces* sp. XY533, where the *svn* BGC was first characterized^22^, and *Streptomyces* sp. WAC05950 from our in-house actinomycete collection (**Table S1**). Notably, these biosynthetic genes are flanked by two copies of a streptavidin gene, which encodes a high-affinity biotin-binding protein. Given the presence of genes for producing the biotin biosynthesis inhibitor stravidin and two copies of genes for the biotin-sequestering agent streptavidin in the same chromosomal region, we hypothesized that this gene arrangement represents a first-of-its-kind ‘cluster of clusters’—or megacluster—that targets biotin metabolism.

To test this premise, we queried a local database of antiSMASH^23^-predicted BGCs in *Streptomyces* genomes for homologous sequences encompassing the *svn* BGC and its neighbouring genes. Based on cumulative similarity scores and manual inspection of gene arrangement and content, we identified 37 closely related BGCs. Of these, 29 exhibited near-identical boundary organization, suggesting deep conservation across *Streptomyces* strains (**Fig. 1a**). A subgroup of eight BGCs lacked four upstream genes, forming a mini PKS/NRPS hybrid subcluster. As part of this effort, we revisited eleven *Streptomyces* strains previously identified as biotin antimetabolite producers in a high-throughput screen of 960 natural product extracts, but which had not been genomically characterized^20^. Whole-genome sequencing of these strains revealed six unique genomes—remarkably, all encode the megacluster (**Fig. S1**). Analysis of the sequences flanking the *svn* BGC revealed additional BGCs, including one encoding acidomycin, as well as two other previously uncharacterized subclusters that we describe below as producing dapamycins and α-Me-KAPA derivatives. Strikingly, the prevalence and conservation of this megacluster rival the frequency of well-studied *Streptomyces* BGCs responsible for streptomycin and tetracycline production (**Fig. 1b**).

**Figure 1.**
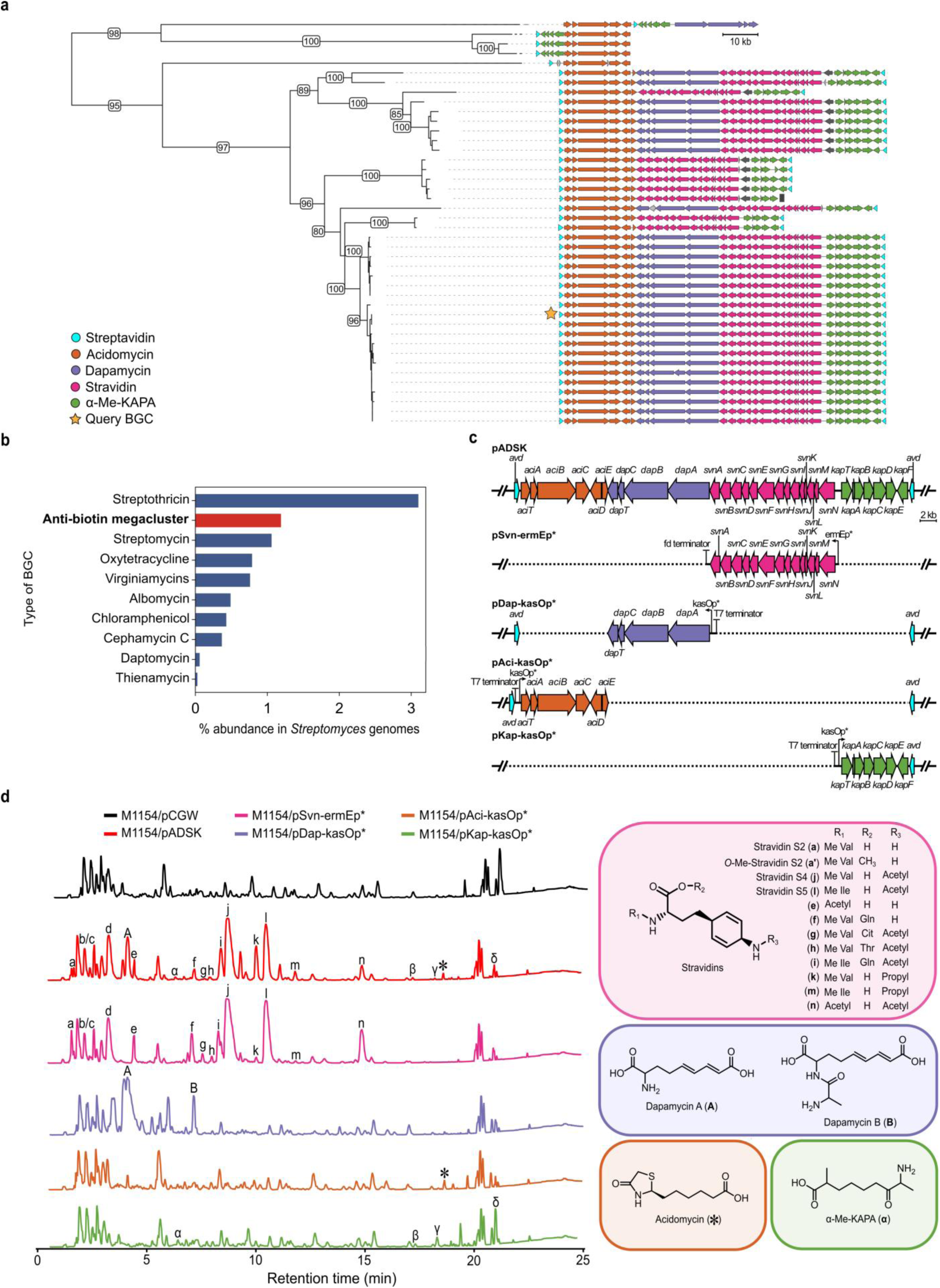
Discovery of a conserved anti-biotin megacluster and structural characterization of its metabolites. **a**, Phylogenetic reconstruction using the anti-biotin megacluster from *Streptomyces* sp. WAC05950 (yellow star) as the query gene cluster. The subclusters are coloured as: orange, acidomycin; lavender, dapamycin; pink, stravidin; green, α-Me-KAPA. Streptavidin (cyan) genes mark the boundary of the megacluster. **b,** Prevalence and conservation of the anti-biotin megacluster in *Streptomyces* genomes, compared to well-characterized antibiotic BGCs. BGC counts were based on the presence of >50% of genes conserved across 3,287 unique NCBI *Streptomyces* genomes. **c,** Schematic representation of the refactored subclusters. Genes are coloured as in (**a**). The dashed lines represent deleted regions from pADSK. **d**, Comparative metabolic profiling of refactored subclusters heterologously expressed in *S. coelicolor* M1154. Compounds a-n are produced from *svn* BGC, A-B from dap BGC, * from *aci* BGC, and α-δ from the *kap* BGC. Me Val – methyl valine; Me Ile – methyl isoleucine; Cit – citrulline.

### Anti-biotin megacluster encodes four distinct biosynthetic subclusters

To assess the small-molecule biosynthetic potential of the megacluster, we captured a 65,808 bp DNA containing the megacluster from the genome of *Streptomyces* sp. WAC05950 into pADSK. In parallel, we cloned and refactored the known *svn* BGC driven by the strong constitutive promoter ermEp* (pSvn-ermEp*)^24^ to assess its individual metabolic output (**Fig. 1c, Fig. S2**). Both constructs were introduced into the surrogate host *S. coelicolor* M1154 for heterologous expression and comparative metabolic profiling^25^. Mass spectrometry (MS) analysis revealed that the refactored *svn* BGC alone produced a suite of stravidin analogs (a–n) (**Fig. 1d, Fig. S3**). In contrast, the full megacluster (pADSK) yielded a broader metabolite profile, including several additional peaks (A, α, β, γ, δ & *) in the base peak chromatogram (BPC) (**Fig. 1d**). One of these peaks (*) matched the theoretical molecular mass of acidomycin (calculated [M+H]^+^ = 218.0845; observed 218.0853) and exhibited an identical MS/MS fragmentation pattern to that of a commercial standard (**Fig. S4**), suggesting that acidomycin is encoded within a distinct subcluster. Indeed, *S. virginiae* ISP-5094, one of the strains identified in the phylogenetic analysis, is known to produce acidomycin^26^.

To assign the additional compounds to their respective subclusters, we generated targeted gene deletions within the pADSK construct and performed metabolic profiling of the resulting strains **(Fig. 1c, S5, S6a**)^27^. This approach deconvoluted the megacluster into four functional subclusters, each responsible for the biosynthesis of a distinct compound series: the *svn* BGC (stravidins, peaks a–n), the *dap* BGC (dapamycins, peaks A and B), the *aci* BGC (acidomycin, peak *), and the *kap* BGC (α-Me-KAPA derivatives, peaks α–δ) (**Fig. 1d, S6b**).

We next purified and structurally characterized representative members from each compound class. MS/MS fragmentation data revealed 12 predicted known and unknown stravidin analogs from the *svn* BGC, of which four—stravidin S2 (a), *O*-methyl-stravidin S2 (a′), stravidin S4 (j), and stravidin S5 (l)—were purified and confirmed by one- and two-dimensional nuclear magnetic resonance (NMR) spectroscopy (**Fig. S18–S37, Tables S2, S3**). Acidomycin (*) was similarly isolated from the *aci* BGC (**Fig. S38-42, Table S4**). This corrects a previous misannotation that assigned acidomycin production to the core biotin operon (*bioFBAD*) in *S. virginiae* ISP-5094^28^. Using targeted deletions in both *Streptomyces* sp. WAC05950 and the pADSK construct, we confirmed that the *aci* BGC is necessary and sufficient for acidomycin biosynthesis (**Fig. S7, S8**). Peaks A and B, derived from the *dap* BGC, were elucidated as 2-amino-5,7-dienenonanedioic acid (dapamycin A) and 2-*N*-alanyl-5,7-dienenonanedioic acid (dapamycin B), respectively (**Fig. S43–S52, Table S5**). From the *kap* BGC, we identified peak α as 2-methyl-7-keto-8-aminopelargonic acid (α-Me-KAPA) and peak γ as its *N*-acetylated derivative. Peaks β and δ were determined to be spontaneously dimerized pyrazine derivatives of α-Me-KAPA and were excluded from further functional analyses (**Fig. S53–S67, Tables S6–S7**). In parallel, we confirmed that *Streptomyces* sp. WAC05950, the native producer of the megacluster, co-produces acidomycin, dapamycin B, and α-Me-KAPA under laboratory conditions (**Fig. S11**). This co-production validates that these metabolites are not only genetically encoded but also biosynthesized in their native context.

Together, these findings demonstrate that the megacluster encodes four distinct subclusters, each capable of producing a unique set of small molecules independently and in the context of the full megacluster. From the *svn* BGC, we purified previously known stravidins (S2, S4, S5), and a novel methylated stravidin S2 analog. The *aci* BGC was identified for the first time, coinciding with the first observation of its co-production with stravidins. The *dap* BGC yielded a series of novel, uncharacterized molecules, which we named dapamycins. The *kap* BGC produced a series of compounds, with α-Me-KAPA as the primary product.

### Megacluster compounds are antimicrobial under biotin limitation

Bacterial survival under biotin-limited conditions requires *de novo* biosynthesis of biotin (**Fig. 2a**)^29^. To evaluate the antibacterial activity of the compounds produced by each subcluster, we performed susceptibility testing against a panel of Gram-negative and mycobacterial species using nutrient-minimal or biotin-free media (**Fig. S12a**). Each subcluster yielded at least one compound with biotin-suppressible activity (**Fig. 2b**).

**Figure 2.**
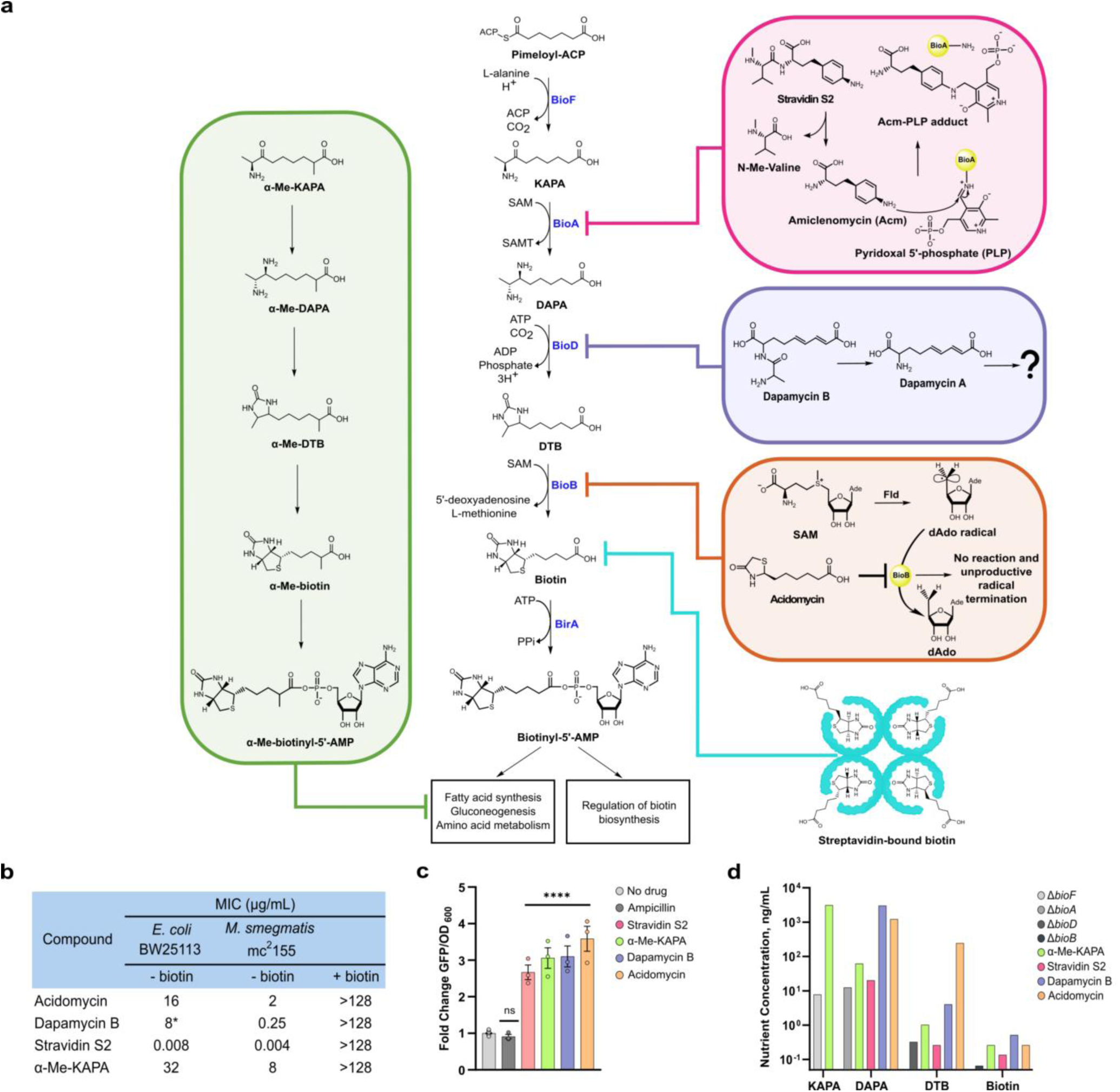
The megacluster produces four compounds targeting bacterial biotin metabolism. **a,** A conserved stage of biotin biosynthesis and its downstream utilization, highlighting the mechanisms of action of known and proposed enzymatic targets for the isolated compounds. Pimeloyl-ACP is converted to biotinyl-5-AMP, a bioactive form of biotin, via five enzymatic reactions involving BioF, BioA, BioD, BioB and BirA. **b,** Antibacterial activities of acidomycin, dapamycin B, stravidin S2 and α-Me-KAPA against *E. coli* BW25113 and *M. smegmatis* mc^2^155 are suppressed with biotin supplementation. An asterisk next to a MIC value denotes transient inhibition, with bacterial growth appearing after a ∼8-hour growth delay. **c,** Upregulation of the *bioB* operon promoter upon incubation with the four isolated compounds. Compounds were tested at a 0.5×MIC concentration in *E. coli* harbouring a *bioB* ALON plasmid. Ampicillin served as a negative control. Data are representative of three biological replicates and presented as mean ± SEM. Statistical significance was determined using a one-way analysis of variance (ANOVA) with Dunnett’s multiple comparisons test. \*\*\*\**p* < 0.0001. NS, not significant. **d,** Minimum amounts of biotin and its biosynthesis intermediates required to rescue auxotrophic mutants or suppress the activities of the four compounds by more than 4-fold. Data are representative of at least three biological replicates from the checkerboard microdilution assays in **Fig. S13**.

Stravidin S2 exhibited the highest antibacterial potency among its analogs; in contrast, its *O*-methylated derivative (*O*-Me-S2) and acetylated forms (S4 and S5) showed markedly reduced activity and were excluded from further investigation. Acidomycin, previously reported to act only against mycobacteria^15^, demonstrated a broader spectrum of activity in our assays, including inhibition of selected Gram-negative strains. Dapamycin A inhibited mycobacterial growth and had no detectable effect on Gram-negative bacteria, while dapamycin B showed transient inhibition in Gram-negative species and more sustained activity in mycobacteria (**Fig. S12b**). Among all tested compounds, α-Me-KAPA displayed the widest spectrum of activity across both bacterial groups. Based on the potent *in vitro* susceptibility of *E. coli* BW25113 and *M. smegmatis* mc²155 to these NPs, these two strains were selected as model organisms for downstream mechanistic studies.

### Antibacterial compounds inhibit bacterial biotin metabolism

Stravidins are di- and tripeptide prodrugs that release amiclenomycin (Acm)^14^, a non-proteinogenic amino acid, that irreversibly inhibits BioA by forming a covalent adduct with its pyridoxal 5′-phosphate (PLP) cofactor^13^. Acidomycin is a competitive inhibitor of BioB that prevents the natural substrate, D-dethiobiotin (DTB), from being converted into biotin^15, 16^. Dapamycins A/ B, newly identified in this study, share structural features with known intermediates of the biotin pathway—KAPA and DAPA—while α-Me-KAPA is a methylated analog of KAPA previously isolated from *Streptomyces diastaticus*, though not examined for antibacterial activity^30^. These structural similarities led us to hypothesize that dapamycins and α-Me-KAPA may also disrupt biotin metabolism as pathway antimetabolites.

To validate dapamycin A/B and α-Me-KAPA as biotin biosynthesis inhibitors, we examined the activation of the biotin operon promoter (*bioB*), a reporter of biotin starvation^31^, in response to these NPs. Exposure to stravidin S2, acidomycin, α-Me-KAPA, and dapamycin B significantly upregulated *bioB* promoter activity compared to the negative and untreated controls, consistent with the thesis that all of these compounds deplete the biotin pool (**Fig. 2c**).

To explore which steps of the biotin biosynthetic pathway dapamycin A/B and α-Me-KAPA could target, we assessed whether supplementation with pathway intermediates could rescue bacterial growth, a method previously used to identify a synthetic inhibitor of BioA^19^. First, we established the suppression profiles for the known inhibitors of biotin biosynthesis—stravidin S2 and acidomycin—in *E. coli*. Both compounds were potently suppressed by biotin at concentrations as low as 0.125 ng/mL. As expected, stravidin S2 activity was suppressed ≥16-fold by the downstream metabolites of BioA— DAPA and DTB. Notably, supplementation with KAPA, the natural substrate of BioA, had no effect, likely due to the formation of a stable, irreversible Acm–PLP adduct that renders BioA catalytically inactive (**Fig. S13a)**^13, 14^. Acidomycin activity was suppressed ≥16-fold by DTB but only at high concentrations (>250 ng/mL) and partially suppressed by KAPA and DAPA at even higher levels (>1.56 µg/mL and >1.28 µg/mL, respectively, **Fig. S13b**), consistent with its role as a reversible BioB inhibitor.

We next applied this approach to dapamycin B and α-Me-KAPA. Dapamycin B exhibited no suppression with KAPA, weak suppression with DAPA, and inconclusive suppression with DTB, which may be attributed to compound instability in *E. coli* (**Fig. S13c**). In contrast, assays in *M. smegmatis* were more consistent; DTB rescued growth at low concentrations (>4 ng/mL), while KAPA and DAPA had no effect (**Fig. S13d**), implicating BioD as a probable target. By comparison, the activity of α-Me-KAPA in *E. coli* was suppressed ≥16-fold by KAPA, DAPA, and DTB, with KAPA requiring the highest concentration (>3.13 µg/mL, **Fig. S13e**). This suppression profile resembled that of stravidin S2, suggesting that the target of α-Me-KAPA may be related to BioA (**Fig. S13a, e**).

To further support these target assignments, we compared the minimum concentration of pathway intermediates required to suppress compound activity with those needed to restore growth in biotin pathway auxotrophs, Δ*bioFADB* (**Fig. 2d**). The nutrient levels required to recover auxotrophies closely matched the suppressing concentrations for the established targets, such as DAPA suppression (Δ*bioA*) of stravidin S2 and biotin suppression (Δ*bioB*) of acidomycin. Similarly, the suppressing concentrations of DAPA and DTB for α-Me-KAPA and dapamycin B, respectively, aligned with those required to restore growth in Δ*bioA* and Δ*bioD* strains, further supporting BioA and BioD as their likely targets.

### Dapamycins are prodrugs that inhibit dethiobiotin synthase

Dapamycin B exhibits antimicrobial activity against *E. coli* BW25113 and *M. smegmatis* mc^2^155, however, dapamycin A shows no activity to most tested bacteria except for weak antimicrobial activity against *M. tuberculosis* H37Ra and *M. bovis* BCG Pasteur ATCC 35734 (**Fig. S12a**). We envision that dapamycin B may be imported through a peptide transporter and then hydrolyzed to release the active warhead of dapamycin A in a similar prodrug fashion to that of stravidin. This hypothesis was supported by the deletion of a dipeptide transporter gene, *dppC*, in *E. coli* that abolished the antibacterial activity of dapamycin B (**Fig. S12c**), suggesting that compound uptake is transporter-dependent. Quantitative LC-MS analysis confirmed that dapamycin B is metabolized intracellularly into dapamycin A, supporting the hypothesis that dapamycin B functions as a prodrug (**Fig. S14a**). In a *dppC* deletion strain, extracellular levels of dapamycin B were greatly increased compared to the WT strain, while levels of dapamycin A were reduced, further implicating DppC in both uptake and conversion (**Fig. S14a**).

To investigate whether dapamycin A/B act on enzymes in the biotin pathway, we first examined the effect of gene overexpression on compound activity in *M. smegmatis*. Overexpression of *bioD*, but not *bioFAB*, suppressed the activity of dapamycin B, implicating BioD as its target (**Fig. 3a**). As a positive control, stravidin S2 activity was specifically suppressed by *bioA* overexpression, confirming the assay’s specificity, while activities of ethambutol, amikacin, and ciprofloxacin were unaffected by *bioFADB* overexpression (**Fig. S14b**). To verify the functionality of the overexpressed enzymes, plasmids encoding *bioFADB* were introduced into the corresponding *E. coli* knockouts, which rescued their auxotrophies with the exception of *bioB*, as expected (**Fig. S14c**)^16^. We next tested whether dapamycin A or B directly inhibited purified BioD. Neither compound caused significant inhibition of BioD enzymatic activity (**Fig. 3b**). We then assessed their effects on the upstream enzyme BioA. Likewise, neither dapamycin A nor B measurably inhibited BioA compared to the positive control, stravidin S2 (**Fig. 3c**). These findings suggest that dapamycin B perturbs biotin metabolism via BioD inhibition, likely mediated by a metabolite yet to be structurally characterized.

**Figure 3.**
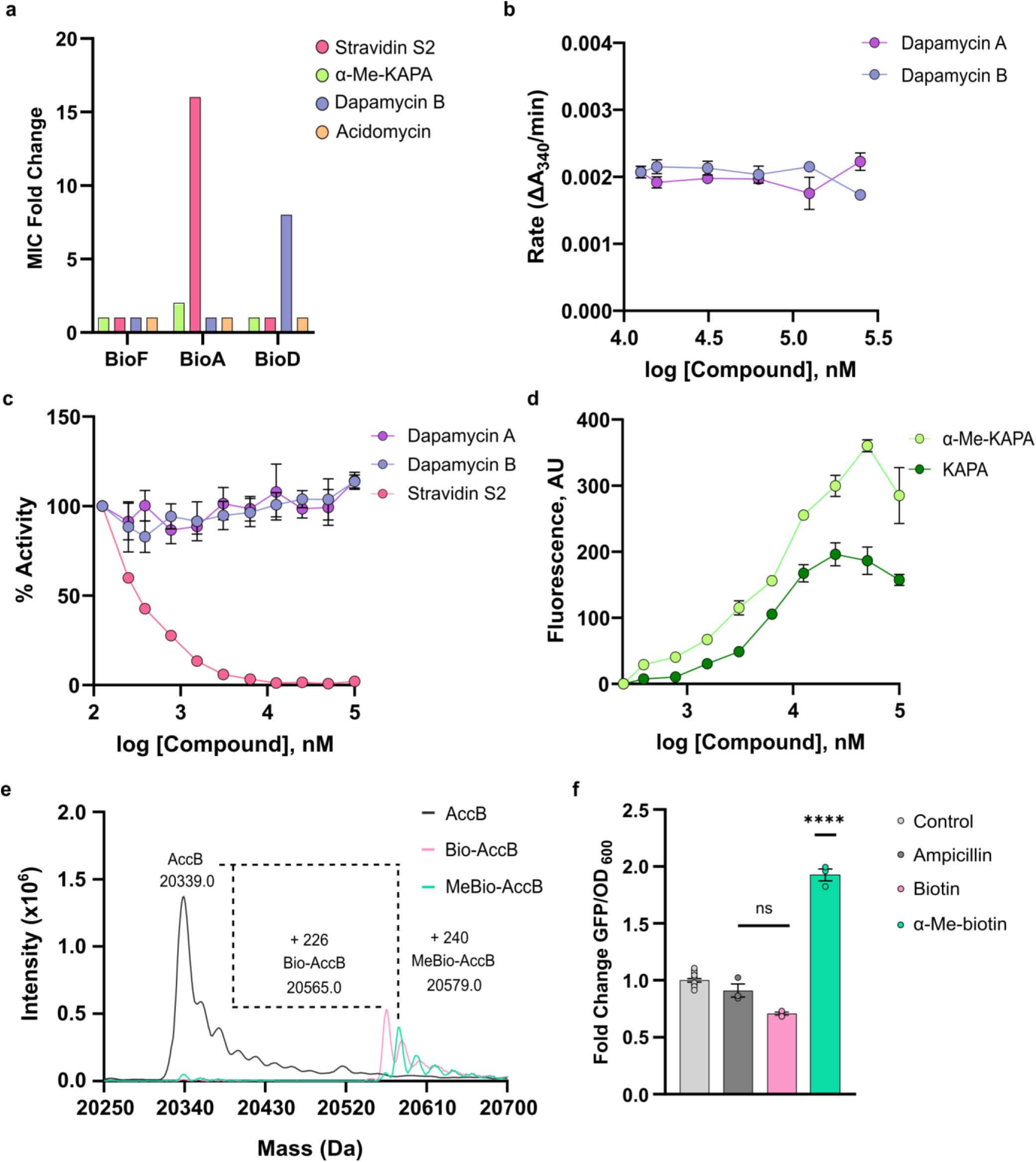
Cellular and enzymatic evaluation of dapamycins and α-Me-KAPA in biotin metabolism inhibition. **a,** Activity of stravidin S2, α-Me-KAPA, dapamycin B, and acidomycin against *M. smegmatis* overexpressing *bioFADB* in the presence of the inducer, theophylline. Data are representative of two biological replicates. **b**, Rates of recombinant *E. coli* BioD enzymatic reactions, measured via the coupled PK/LDH assay, across 15–250 µM of dapamycin A (purple) and dapamycin B (blue). Data are represented as means ± SEM (n = 3). **c**, Activity of the recombinant *E. coli* BioA across 0.1–100 µM of dapamycin A (purple) and B (blue). Stravidin S2 (red) is a positive control for inhibition. Data are represented as means ± SEM (n = 3). **d**, Fluorescence signal from the BioA reaction with KAPA (green) or α-Me-KAPA (light green) as substrates, measured across the indicated concentration ranges. Data are represented as means ± SEM (n = 3). **e**, Intact protein LC ESI–MS analyses of AccB (10 μM) incubated for 3 h with BirA in buffer either lacking (black trace) or containing 100 μM of biotin (pink) or α-Me-biotin (turquoise). **f**, Regulation of the *bioB* operon promoter in response to biotin (pink) and α-Me-biotin (turquoise). Compounds were tested in *E. coli* harbouring a *bioB* ALON plasmid. Ampicillin (dark grey) served as a negative control. Data are representative of three biological replicates and presented as mean ± SEM. Statistical significance was determined using a one-way analysis of variance (ANOVA) with Dunnett’s multiple comparisons test. ****p < 0.0001. NS, not significant.

To investigate whether dapamycin B undergoes further intracellular modification, we profiled *E. coli* cultures treated with the compound using high-performance liquid chromatography coupled with high-resolution quadrupole time-of-flight mass spectrometry (HPLC-HR-QTOF-MS) in isotope-labelled minimal media. In addition to dapamycin A, we detected several new metabolites absent in untreated controls (**Fig. S15a–d**). One species, designated M, exhibited a +2 Da shift (M*) in ^13^C-glucose media but no shift with ^15^N-ammonium sulphate, indicating the incorporation of two carbons (**Fig. S15e**). Although the precise structure of M remains unresolved, these data suggest that dapamycin A is further processed to generate a modified derivative, potentially responsible for the observed anti-biotin activity.

### α-Me-KAPA hijacks biotin biosynthesis

Given the structural similarity of α-Me-KAPA to KAPA and its metabolite suppression profile resembling that of stravidin S2 (**Fig. 2d and S13a, e**), we investigated whether it functions as a substrate or inhibitor of BioA. Incubation of recombinant *E. coli* BioA with increasing concentrations of α-Me-KAPA produced a dose-dependent fluorescence signal corresponding to enzyme-catalyzed formation of a derivatized α-Me-DAPA–like product, similar to the natural substrate KAPA (**Fig. 3d**). When the reaction was performed without the derivatizing agent and analyzed by LC–MS, a peak was detected with a mass consistent with α-Me-DAPA (calculated [M+H]^+^ = 203.1754; observed 203.1741), and its MS/MS fragmentation pattern matched the expected structure (**Fig. S16a**). Together, these results indicate that α-Me-KAPA acts as a BioA substrate rather than an inhibitor. We next assessed whether downstream enzymes could process α-Me-DAPA in the biotin pathway. Coupling BioA with recombinant BioD yielded a product consistent with a mass of α-Me-DTB (calculated [M+H]^+^ = 229.1547; observed 229.1560) and matching the expected MS/MS fragmentation pattern (**Fig. S16a**).

To evaluate whether these metabolites form in whole cells, we treated *E. coli* with α-Me-KAPA (¼×MIC, 4 h) in minimal media and analyzed cell extracts using LC/MS. We detected peaks matching both α-Me-DTB and α-Me-biotin based on retention time and MS/MS fragmentation, the latter confirmed by comparison to a commercial standard (**Fig. S16b**), demonstrating that α-Me-KAPA is metabolized intracellularly into α-Me-biotin.

We then examined whether α-Me-biotin could serve as a substrate for BirA, the *E. coli* biotin ligase responsible for conjugating biotin to the biotin carboxyl carrier protein (AccB) of acetyl-CoA carboxylase (Acc)^32^. Incubation of α-Me-biotin with recombinant BirA yielded α-Me-biotinyl-5’-AMP (calculated [M+H]^+^ = 588.1636; observed 588.1649), with a fragmentation pattern closely matching that of biotinyl-5’-AMP, confirming that α-Me-biotin is indeed a substrate for BirA (**Fig. S16c**). In a coupled reaction with recombinant AccB, we observed conjugation of α-Me-biotin to AccB via native MS and peak deconvolution, confirming that α-Me-biotin can serve as a functional substrate for biotinylation (**Fig. 3e**).

Together, these findings establish that α-Me-KAPA hijacks the biotin biosynthesis pathway to generate α-Me-biotin, which is subsequently conjugated to AccB by BirA. Unlike native biotin, α-Me-biotin activates the *bioB* operon, similarly to the other NPs, suggesting it does not support normal biotin-dependent metabolism, consistent with an inhibitory role (**Fig. 3f**).

### Anti-biotin compounds from the megacluster act synergistically

Previous studies have shown that amiclenomycin, the warhead of stravidins, synergizes with acidomycin against *M. smegmatis* in minimal media^33^. Similarly, we observed potent synergy between stravidin S2 and acidomycin across multiple Gram-negative species, including *E. coli*, *K. pneumoniae*, and *A. baumannii* (FICI = 0.079–0.376; **Fig. 4a, S17a**). Given this observation, we systematically tested all pairwise combinations of the four compounds.

**Figure 4.**
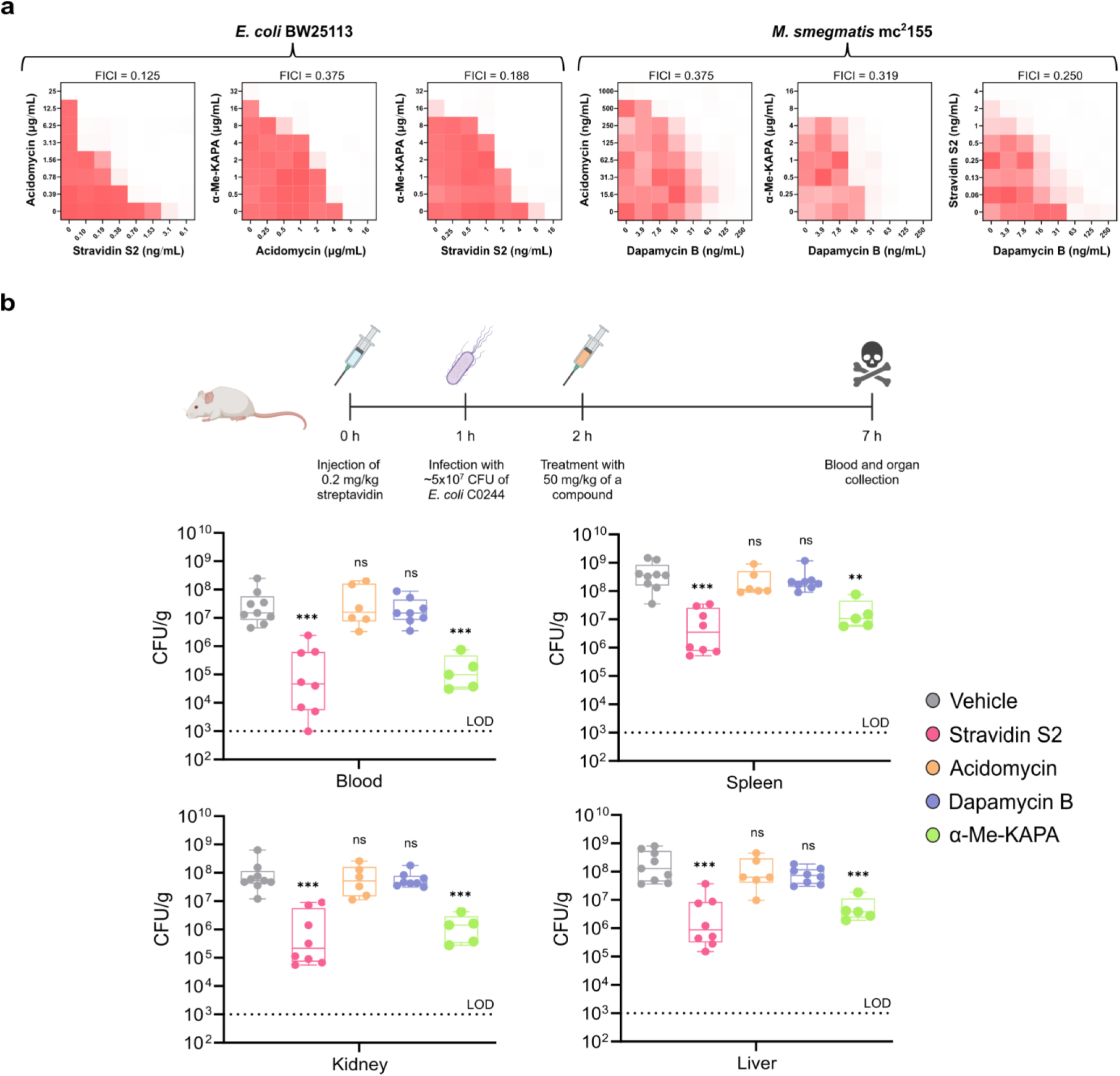
Synergistic activity and *in vivo* efficacy of megacluster-encoded compounds. **a,** Checkerboard microdilution assays of combinations between stravidin S2, acidomycin, dapamycin B and α-Me-KAPA in *E. coli* BW25113 and *M. smegmatis* mc^2^155. Red regions represent higher cell density. Checkerboard data are representative of three biological replicates. FICI = FIC_compound1_ + FIC_compound 2_, where FIC = [X]/MIC_x_ and X is the drug concentration in the presence of the co-drug. **b**, Bacterial load measured in blood and organs following compound treatment of an *E. coli* C0244 systemic infection in CD-1 mice pre-treated with streptavidin (2 mg/kg, 1 h prior to infection). Groups of mice were treated by a single intraperitoneal (IP) injection of vehicle (grey), stravidin S2 (50 mg/kg, pink), acidomycin (50 mg/kg, orange), dapamycin B (50 mg/kg, purple) or α-Me-KAPA (50 mg/kg, green) 1 h after infection. The infection progressed for 6 hours. Each point represents an individual mouse, and the line indicates the mean; \*\**p* <0.01, \*\*\**p* <0.001, non-parametric Mann-Whitney test with Benjamini, Krieger and Yekutieli’s multiple comparisons test. LOD stands for the limit of detection. Created in BioRender.

In *E. coli*, dapamycin B synergized with acidomycin (FICI = 0.31), showed antagonism with stravidin S2 (FICI = 9.00), and had an indifferent interaction with α-Me-KAPA (FICI = 3.00; **Fig. S17b**). α-Me-KAPA synergized with both stravidin S2 and acidomycin in *E. coli* (FICI = 0.19 and 0.38, respectively). By contrast, all compound combinations were synergistic in *M. smegmatis*, with FICI values ranging from 0.38 to 0.5 (**Fig. 4a, S17c**). Together, these findings suggest that the megacluster encodes a coordinated chemical arsenal whose constituents act synergistically to disrupt biotin metabolism across diverse bacterial species.

### *In vivo* efficacy of megacluster-encoded compounds

To assess whether the megacluster-derived compounds are efficacious at reducing bacterial burden *in vivo*, we tested each in a murine model of systemic infection using a previously established protocol that humanizes circulating biotin levels, enabling *in vivo* evaluation of biotin-targeting inhibitors^21^. Mice were challenged with multi-drug resistant *E. coli* C0244, the most susceptible pathogenic strain identified *in vitro* (**Fig. S12a**). Monotherapy with stravidin S2 or α-Me-KAPA (50 mg/kg), administered 1 h post-infection, led to a 97–99% and 95–99% reduction in bacterial burden, respectively, across blood, spleen, kidneys, and liver, relative to vehicle-treated controls (**Fig. 4b**). Neither acidomycin nor dapamycin B (50 mg/kg) showed measurable efficacy under these conditions. These findings demonstrate that at least two members of this natural product ensemble are active *in vivo* against *E. coli* and underscore the therapeutic potential of targeting biotin metabolism.

## Discussion

Natural product BGCs are increasingly recognized not as isolated units but as components of complex, multi-metabolite systems that can act in coordination^11^. Here, we describe a conserved *Streptomyces* megacluster encoding four structurally and mechanistically distinct natural products— stravidins, acidomycin, α-Me-KAPA, and dapamycins—alongside streptavidin, a high-affinity biotin-binding protein. These encoded products converge on a single conserved pathway, bacterial biotin metabolism.

While synergy is often viewed as an emergent pharmacological property, discovered through empirical drug pairing, our findings support a growing body of evidence suggesting that synergy can also be a genetically programmed feature of natural product biosynthesis. Similar to the well-established co-production of synergistic antibiotics—streptogramins^17^ and clavulanic acid/cephamycin C^34^—the anti-biotin megacluster encodes multiple compounds that each inhibit or hijack different enzymes within the same pathway and display enhanced potent antimicrobial potency when combined. Intriguingly, these properties are recapitulated across two phylogenetically distant bacteria, including both Gram-negative species and mycobacteria.

Mechanistic studies revealed that each compound produced by the megacluster interferes with biotin metabolism through a distinct mode of action. Stravidin irreversibly inhibits BioA via its warhead, amiclenomycin^13^. Acidomycin competitively blocks BioB^15^. Dapamycin B functions as a prodrug that enters cells via the Dpp transporter and is metabolized to dapamycin A, which undergoes further conversion and may act on BioD. α-Me-KAPA hijacks the biotin biosynthetic pathway to form α-Me-biotin, which is conjugated to the AccB subunit of acetyl-CoA carboxylase, potentially impairing malonyl-CoA production and downstream fatty acid biosynthesis.

Synergy between these compounds may reflect emergent vulnerabilities within biotin metabolism. For instance, BioA inhibition by stravidin S2 could deplete DAPA pools, increasing sensitivity to BioB inhibition by acidomycin. Conversely, acidomycin-mediated depletion of S-adenosylmethionine^15^, a shared cofactor for biotin enzymes, may secondarily impair BioA and potentiate stravidin activity. These potential reciprocal dependencies suggest that the encoded metabolites are not merely co-produced but co-evolved for collaborative function. This concept resonates with prior models of selective pressure driving the evolution of superclusters for antibiotic synergy^11, 35^.

Genomic analyses underscore the evolutionary significance of this system. The anti-biotin megacluster is widely distributed across *Streptomyces* genomes, with a prevalence comparable to canonical BGCs for tetracycline and streptomycin. Its prior obscurity may reflect limitations of historical screening efforts, which relied heavily on nutrient-rich media that mask the conditional essentiality of biotin metabolism. Indeed, the megacluster-containing *Streptomyces* WAC05950 strain used in this study was first uncovered through a screen exploiting a biotin transporter deletion strain—a strategy that highlights how genome mining under physiologically relevant conditions can reveal otherwise hidden biosynthetic systems^20^.

*In vivo* studies demonstrated that stravidin S2 and α-Me-KAPA exhibit significant efficacy as monotherapies in a murine model of multidrug-resistant *E. coli* infection. Although *in vivo* combination therapies were not pursued due to limited compound availability and the complexity of dosing optimization, the strong synergy observed *in vitro* suggests that co-administration could further enhance efficacy. Future studies will aim to systematically evaluate such combinations to realize their full therapeutic potential. These efforts echo nature’s blueprint, exemplified by co-produced synergistic antibiotic pairs like Synercid^17^.

Together, our findings highlight a multi-tiered strategy for targeting a conserved bacterial pathway using genetically encoded, chemically distinct metabolites. The architecture of the anti-biotin megacluster provides a paradigm for naturally evolved combination therapies, supporting a shift in antibiotic discovery from isolating individual hits to reconstructing native synergistic systems. As genome mining methods advance, the identification of similar megaclusters may reveal new paths for overcoming antimicrobial resistance by mimicking nature’s strategies.

## Materials and Methods

### Oligonucleotides, strains, plasmids, and reagents

Oligonucleotides and gBlocks were synthesized by Integrative DNA Technologies (Coralville, IA, USA) and listed in **Table S8**. All plasmids and strains used in this work are listed in **Table S9** and **S10**. Sanger sequencing and Illumina genome sequencing were performed at the McMaster Genomics Facility (McMaster University). Long-read genome sequencing was performed on an in-house MinION platform (Oxford Nanopore Technologies). Dreamtaq Master Mix (2×) and Phusion Hi-Fidelity DNA polymerase (Thermo Fisher Scientific) were used for PCR screening and high-fidelity PCR amplification. GeneJet plasmid miniprep kit (Thermo Fisher Scientific) was used for general plasmid purification. pADSK and its truncated derivative plasmids were purified using the alkaline lysis method. Restriction enzymes were purchased from Thermo Fisher Scientific. 5-Fluoroorotic acid monohydrate (5-FOA) was purchased from Cedarlane. Zymolyase 20T was purchased from Bioshop Canada. KAPA hydrochloride was purchased from Cayman Chemical Company. DAPA dihydrochloride was purchased from Toronto Research Chemicals. Streptavidin was purchased from Neuromics. All other chemicals used in this study were purchased from Sigma-Aldrich.

### Growth conditions

*E. coli* strains were grown in LB broth (Bioshop Canada) and incubated at 37℃, 250 rpm. When culturing *E. coli* EPI300 strains with captured or refactored plasmids carrying NP BGCs for plasmid isolation, 1 mM of L-arabinose was added to induce plasmid copy number^36^. *S. cerevisiae* VL6-48N^37^ was grown in YPD (10 g yeast extract, 20 g peptone, 20 g dextrose, ddH_2_O 1 L) liquid or agar medium supplemented with 100 µg/mL adenine for spheroplast cells preparation and SD-Trp-5-FOA medium (182 g sorbitol, 20 g glucose, 20 g agar, 100 mL 10× yeast nitrogen base, 100 µg/mL adenine, 1 mg/mL 5-FOA, ddH_2_O 1 L) for selection. 10× yeast nitrogen base (100 mL): 1.7 g yeast nitrogen base w/o amino acids and ammonium sulphate (BD Difco), 5 g (NH_4_)_2_SO_4_, 0.832 g amino acids mix w/o tryptophan, 100 µg/mL adenine; 100× adenine (10 mg/mL), and 100× 5-FOA (100 mg/mL) were added to SD-Trp medium after autoclave. Yeast transformants were grown in SD-Trp liquid medium (182 g sorbitol, 20 g glucose, 100 mL 10× yeast nitrogen base, 100 µg/mL adenine, ddH_2_O 1 L) for plasmid DNA extraction. *Streptomyces* strains were grown in TSBY medium (30 g tryptone soy broth, 5 g yeast extract, ddH_2_O 1 L) and incubated at 30℃, 250 rpm for genomic DNA isolation and seed culture preparation. When preparing genomic DNA, 0.5% glycine was added to the TSBY culture. Soy flour medium (SFM, 20 g soya flour, 20 g D-mannitol, 20 g agar, ddH_2_O 1 L, pH 7.2-7.4) was used for *Streptomyces* sporulation and conjugation (supplemented with 20 mM MgCl_2_). *Streptomyces* antibiotic activity medium (SAM, 15 g glucose, 15 g soytone, 5 g NaCl, 1 g yeast extract, 1 g CaCO_3_, 2.5 mL glycerol, ddH_2_O 1 L, pH 6.8), YBP medium (10 g glucose, 2 g yeast extract, 2 g beef extract, 4 g peptone, 1 g MgSO_4_, 15 g NaCl, ddH_2_O 1 L, pH 7.2-7.4), and *Streptomyces* minimal medium (SMM, 5 g (NH_4_)_2_SO_4_, 0.5 g K_2_HPO_4_, 0.2 g MgSO_4_• 7H_2_O, 0,01 g FeSO_4_•7H_2_O, 10 g glucose, 2 g CaCO_3_, ddH_2_O 1 L, pH 7.0) were used for fermentation. Antibiotics were added as required for selection (100 µg/mL ampicillin, 50 µg/mL kanamycin, 50 µg/mL apramycin, 25 µg/mL chloramphenicol, 25 µg/mL nalidixic acid, and 50 µg/mL trimethoprim).

### Closing the gaps of the anti-biotin megacluster

The whole genome sequence of *Streptomyces* sp. WAC05950 has been deposited into Genbank under the accession number RQJB00000000.1. The full anti-biotin megacluster was located on two different contigs. svn-gF/svn-gR primers were used to amplify the 143 bp gap region and confirmed by Sanger sequencing.

### Megacluster phylogeny construction

Custom Python scripts were used to extract all AciB protein sequences from megacluster homologs. These sequences were then aligned alongside the BlmIV Bleomycin NRPS (AAG02364.1) using Muscle v5.1 with default settings^38^. The alignment was trimmed using TrimAL v1.4.15 to exclude columns with any gaps, leaving a final alignment length of 1,412 AAs^39^. IQTree v2.3.4 constructed a maximum likelihood phylogeny with 1000 ultrafast bootstraps and the bleomycin NRPS as an outgroup, which was trimmed from the final phylogeny^40, 41^. We visualized the tree in iTOL and aligned clusters using clinker v0.0.31 before concatenating the resulting SVG files in Illustrator^42, 43^.

### NCBI *Streptomyces* megacluster search

All available *Streptomyces-*genus genomes from NCBI were downloaded using the NCBI datasets tool and used antiSMASH v7.1.0 to predict associated BGCs^23^. From these, a custom Python script extracted associated proteins from the resulting cluster GenBank files, with annotations of associated BGC and genome to a separate FASTA file. DIAMOND v2.1.9 makedb then made a DIAMOND database from this file, for querying other BGCs against.

To query and quantify BGCs in *Streptomyces,* we used a combination of DIAMOND v2.1.9 and custom Python scripts, with especially heavy reliance on the pandas v2.2.2^44^ and biopython v1.83 libraries^45^. First, query BGC amino acid sequences were extracted to a FASTA file, which was used as input for a DIAMOND blastp search using the –very-sensitive, --max-target-seqs 10000, --approx-id 30 and –query-cover 50 flags. The top 250 BGC hits from this output were parsed from cumulative bitscore against the query, and proteins from these BGCs were sorted into another fasta. This was converted to a DIAMOND database again for another more sensitive DIAMOND blastp search using the query BGC and the --max-target-seqs 100000 and --ultra-sensitive flags. The results were manually curated based on the presence or absence of genes within the BGC, as well as sequence and synteny conservation to determine homologous BGCs of the megacluster.

To quantify the presence of other *Streptomyces*-associated BGCs, the number of BGCs that contained >50% of query BGC proteins with a bitscore of >50% to a perfect match was counted. The plot was created with the Python Seaborn and matplotlib libraries.

### *Streptomyces* genome sequencing

*Streptomyces* strains were precultured in 12 mL TSBY medium supplemented with a single glass bead at 30°C for 1 day. Cultures were then inoculated at 5% (v/v) into 50 mL TSBY containing 0.5% (w/v) glycine and 10.3% (w/v) sucrose, and grown at 30°C with shaking at 200 rpm for 2 days. Cells were harvested by centrifugation, and genomic DNA was extracted from 0.8 g of wet cell pellet using the Wizard Genomic DNA Purification Kit (Promega). Strains were dereplicated by BOX-PCR^46^, resulting in seven unique isolates, including the previously sequenced WAC05950. Whole-genome sequencing was performed by Plasmidsaurus following the provider’s standard protocol.

### CORASON analysis of megacluster-containing WAC strains

To assess evolutionary relationships among BGCs, we used CORASON (CORe Analysis of Syntenic Orthologues to prioritize Natural product BGCs). The AciB protein from the acidomycin BGC served as the query, and *Streptomyces* sp. WAC05950 was used as the reference genome, along with additional strains containing identified megalcuster. CORASON aligned clusters based on shared core genes and extracted AciB sequences, which were aligned using MUSCLE v5.1 (default settings). A neighbour-joining tree was generated and visualized with iTOL alongside the aligned clusters.

### TAR cloning of the megacluster and *svn* BGC

Annotation of the anti-biotin megacluster is shown in **Table S1**. TAR cloning was performed according to the standard protocol^47^. Capturing hook sequences were designed as previously described^48^. ADSK-gbk, possessing the capturing hooks targeting the megacluster, was inserted into the pCGW plasmid between the *Nde*I/*Xho*I sites using Gibson assembly. pADSK-cap plasmid was linearized by *Pme*I digestion and purified through a PCR clean-up kit for transformation into yeast spheroplast cells. High-quality genomic DNA of WAC05950 was prepared using a salting out procedure^49^ followed by RNase A treatment to remove RNA. gDNA was treated with *EcoR*V/*Spe*I digestion to release the megacluster and then purified through sodium acetate precipitation. Linearized pADSK-cap plasmid (∼500 ng) and digested gDNA (∼2 µg) were mixed and co-transformed into *S. cerevisiae* VL6-48N spheroplast cells and then plated onto SD-Trp-5-FOA medium for selection. After incubating at 30℃ for 3-5 d, yeast transformants were picked into SD-Trp liquid medium and grown for 24 h at 30℃. Yeast cells were harvested for plasmid extraction using the alkaline lysis method and screened for positive transformants through PCR using svn-dF/svn-dR as screening primers. Positive hits were selected and re-transformed into *E. coli* EPI300 cells through electroporation and further confirmed through restriction digestion. *svn* BGC was cloned and refactored using the same procedure. ermEp*-svnN-svnA-fd terminator-gbk was used to capture *svn* BGC from pADSK carrying the megacluster. ermEp*-svnN-svnA-fd terminator-gbk was inserted into pCGW plasmid between the *Nde*I/*Xho*I sites using Gibson assembly, generating pSvn-cap plasmid. pADSK was digested with *EcoR*I/*Nsi*I digestion to release the *svn* BGC and then purified through sodium acetate precipitation. *Pme*I linerized and purified pSvn-cap plasmid (∼500 ng) and digested pADSK plasmid (∼2 µg) were mixed and co-transformed into *S. cerevisiae* VL6-48N spheroplast cells for selection of pSvn-ermEp* plasmid.

### Refactoring of pADSK plasmid

λ-red mediated PCR targeting^27^ was used to determine the boundary of each subcluster and refactor each subcluster accordingly. kasOp*^50^-T7 terminator-aac(3)IV-ΔSK-gbk and aac(3)IV-T7 terminator-kasOp*-kap-gbk (**Table S8**) were synthesized and introduced into pUC18 between *EcoR*I/*Hind*III sites to provide editing templates for refactoring each subcluster. Error-free editing templates were confirmed by Sanger sequencing using M13F/R primers. Schematics of pADSK plasmid editing are shown in **Fig. S5**. Genotypes of the refactored plasmids are listed in **Table S9**.

*pADSKΔsvn:* Δsvn-F/R primers were used to amplify the *aac(3)IV* resistant cassette from pCGW, followed by gel purification (**Table S8**). The purified *aac(3)IV* cassette (1 µg) was transformed into *E. coli* BW25113/pKD46/pADSK competent cells (100 µL) through electroporation, plated onto LB agar medium supplemented with 50 μg/mL apramycin and then incubated at 37℃ overnight. The transformants from the agar plate were screened using Δsvn-dF/dR primers by colony PCR. Plasmid DNA was extracted from the positive transformant and then re-transformed into *E. coli* EPI300 cells through electroporation to purify clean knockout mutant, resulting in pADSKΔsvn::aac(3)IV plasmid. pADSKΔsvn::aac(3)IV plasmid was then purified through alkaline lysis and digested with *Avr*II to remove the *aac(3)IV* marker. *Avr*II-treated pADSKΔsvn::aac(3)IV plasmid was purified through sodium acetate precipitation and re-circularized by self-ligation using T4 DNA ligase, resulting in pADSKΔsvn. pADSKΔsvn2, pADSKΔsvn3, pADSKΔDS, pAci and pDSK were generated using an identical procedure except for the primers used for amplifying *aac(3)IV* resistant cassette and colony PCR screening (**Table S8**).

*pAD-kasOp*:* ΔSK-F/R primers were used to amplify the kasOp*-T7 terminator-aac(3)IV-ΔSK-gbk synthetic cassette from pUC18-k*TaSK, followed by gel purification. The purified synthetic cassette (1 µg) was transformed into *E. coli* BW25113/pKD46/pADSK competent cells (100 µL) through electroporation, plated onto LB agar medium supplemented with 50 μg/mL apramycin and then incubated at 37℃ overnight. The transformants from the agar plate were screened using ΔSK-dF/dR primers by colony PCR. Plasmid DNA was extracted from the positive transformant and then re-transformed into *E. coli* EPI300 cells through electroporation to purify clean knockout mutant, resulting in pADSKΔSK::aac(3)IV-kasOp* plasmid. pADSKΔSK::aac(3)IV-kasOp* plasmid was then purified through alkaline lysis and digested with *Nde*I to remove the *aac(3)IV* marker. *Nde*I-treated pADSKΔSK::aac(3)IV-kasOp* plasmid was purified through sodium acetate precipitation and re-circularized by self-ligation using T4 DNA ligase, resulting in pAD-kasOp*.

*pDap-kasOp*:* dap-F/R primers were used to amplify the *aac(3)IV* resistant cassette from pCGW, followed by gel purification. The purified *aac(3)IV* resistant cassette (1 µg) was transformed into *E. coli* BW25113/pKD46/ pAD-kasOp* competent cells (100 µL) through electroporation, plated onto LB agar medium supplemented with 50 μg/mL apramycin and then incubated at 37℃ overnight. The transformants from the agar plate were screened using dap-dF/dR primers by colony PCR. Plasmid DNA was extracted from the positive transformant and then re-transformed into *E. coli* EPI300 cells through electroporation to purify clean knockout mutant, resulting in pAD-kasOp*ΔA::aac(3)IV plasmid. pAD-kasOp*ΔA::aac(3)IV plasmid was then purified through alkaline lysis and digested with *Avr*II to remove the *aac(3)IV* marker. *Avr*II-treated pAD-kasOp*ΔA::aac(3)IV plasmid was purified through sodium acetate precipitation and re-circularized by self-ligation using T4 DNA ligase, resulting in pDap-kasOp*.

*pAci-kasOp*:* aci-F/R primers were used to amplify the kasOp*-T7 terminator-aac(3)IV-ΔSK-gbk synthetic cassette from pUC18-k*TaSK, followed by gel purification. The purified synthetic cassette (1 µg) was transformed into *E. coli* BW25113/pKD46/ pAci competent cells (100 µL) through electroporation, plated onto LB agar medium supplemented with 50 μg/mL apramycin and then incubated at 37℃ overnight. The transformants from the agar plate were screened using dap-dF/dR primers by colony PCR. Plasmid DNA was extracted from the positive transformant and then re-transformed into *E. coli* EPI300 cells through electroporation to purify clean knockout mutant, resulting in pAci::aac(3)IV-kasOp* plasmid. pAci::aac(3)IV-kasOp* plasmid was then purified through alkaline lysis and digested with *Nde*I to remove the *aac(3)IV* marker. *Nde*I-treated pAci::aac(3)IV-kasOp* plasmid was purified through sodium acetate precipitation and re-circularized by self-ligation using T4 DNA ligase, resulting in pAci-kasOp*.

*pKap-kasOp*:* kap-F/R primers were used to amplify the aac(3)IV-T7 terminator-kasOp*-kap-gbk synthetic cassette from pUC18-aTk*K, followed by gel purification. The purified synthetic cassette (1 µg) was transformed into *E. coli* BW25113/pKD46/ pADSK competent cells (100 µL) through electroporation, plated onto LB agar medium supplemented with 50 μg/mL apramycin and then incubated at 37℃ overnight. The transformants from the agar plate were screened using kap-dF/dR primers by colony PCR. Plasmid DNA was extracted from the positive transformant and then re-transformed into *E. coli* EPI300 cells through electroporation to purify clean knockout mutant, resulting in pADSKΔADS::aac(3)IV-kasOp* plasmid. pADSKΔADS::aac(3)IV-kasOp* plasmid was then purified through alkaline lysis and digested with *Xho*I to remove the *aac(3)IV* marker. *Xho*I-treated pADSKΔADS::aac(3)IV-kasOp* plasmid was purified through sodium acetate precipitation and re-circularized by self-ligation using T4 DNA ligase, resulting in pKap-kasOp*.

### Heterologous production of NPs

All plasmids were introduced into *S. coelicolor* M1154^25^ for heterologous expression using an identical procedure^51^. Using pADSK as an example, the procedure is described in the following way. pADSK was electroporated into *E. coli* ET12567 cells, followed by mobilizing into *S. coelicolor* M1154 through *E. coli*-*Streptomyces* interspecies tri-parental mating using *E. coli* ET12567/pADSK as the donor strain, *E. coli* ET12567/pR9406 as the helper strain, and *S. coelicolor* M1154 as the recipient strain. *E. coli* ET12567/pADSK and *E. coli* ET12567/pR9406 cells were grown to OD_600_ = 0.6-1.0; 0.1 mL of each culture was harvested in 1.5 mL microcentrifuge tubes. The *E. coli* cultures were then washed twice with an equal volume of fresh LB and resuspended with 0.1 mL of fresh LB ready for use. *S. coelicolor* M1154 spores were collected from the SFM sporulation plates and resuspended in 2×YT medium (10 g yeast extract, 16 g tryptone, 5 g NaCl, ddH_2_O 1 L), followed by heat activation at 50℃ for 10 mins. The resuspended *E. coli* cells and heat-activated *Streptomyces* spores (equilibrated to room temperature) were combined and plated onto SFM (supplemented with 20 mM MgCl_2_) agar medium. After 16-20 h incubation at 30℃, the conjugation plate was flooded with 1 mL of kanamycin (1.5 mg/mL) and trimethoprim (1.5 mg/mL) and incubated for another 3-5 d at 30℃. Resistant exconjugants were selected to prepare the seed culture for fermentation.

### Metabolic profiling of the NPs

For small-scale metabolic analysis, *Streptomyces* strains were grown in 16 mL test tubes containing 3 mL TSBY medium with three glass beads (5 mm) at 30℃, 250 rpm for 2 d. Antibiotics were added as needed for selection. 150 μL of seed cultures were then inoculated into 3 mL of SAM medium (5% inoculum) in 24-well plates (CR1424, EnzyScreen BV, NL) and grown at 30℃, 250 rpm for 4-7 d. The condition medium was centrifuged, and 2 μL of each sample was analyzed on an Agilent 6550 iFunnel Q-TOF mass spectrometry equipped with an inline Agilent 1290 HPLC system using Xselect CSH C18 column (130 Å, 5 μm, 4.6 mm × 100 mm [Cat no. 186005289, Waters]). HPLC gradient used was: t= 0-0.5 min, 2% B; t=15 min, 20% B; t=22-25 min, 95% B; t=26-30 min, 2% B; at a flow rate of 0.6 mL/min, 30℃ with H_2_O (0.1% formic acid [FA], A) and ACN (0.1% FA, B) as mobile phases. The QTOF system was performed in positive mode with a mass scan range of 50-1000 m/z.

### Production and purification of NPs

*S. coelicolor* M1154 strains carrying different expression plasmids were grown in 100 mL TSBY medium in 250 mL Erlenmeyer flasks (×6) containing 15 glass beads for 3 d at 30℃, 250 rpm as the seed culture. The seed cultures (35 mL) were then inoculated into 2.8 L Thomson ULTR YIELD flasks (×10) containing 700 mL of respective fermentation medium and cultured at 30℃, 250 rpm for 4-7 d. *S. coelicolor* M1154/pSvn-ermEp* strain was grown in YBP medium for 4 d for the production and purification of stravidins. *S. coelicolor* M1154/pAci-kasOp* strain was grown in SAM medium for 7 d for the production and purification of acidomycin. *S. coelicolor* M1154/pDap-kasOp* strain was grown in SMM medium for 4 d for the production and purification of dapamycins. *S. coelicolor* M1154/pADSKΔsvn strain was grown in SMM medium for 4 d for the production and purification of α-Me-KAPA and in SAM for 7 d for the production and purification of 2,5-di-(2-methylhexanoic acyl)-3-methylimidazole and 2,5-dimethyl-3,6-di-(2-methylhexanoic acyl)pyrazine.

#### Purification of stravidin S2, O-Me-S2, S4 and S5

*Streptomyces* culture filtrate (6 L) was extracted with 5% (w/v) HP-20 (Diaion) resin twice. The resin was eluted with 1 mM NH_4_HCO_3_ (pH=8, 1 L), 50% MeOH in water (v/v, 3 L), and 100% MeOH (2 L) to yield three fractions S1 to S3. The active fraction S2 was loaded on an Amberlite C-120 strong cation exchange resin (H^+^ form) column and then eluted with MeOH (500 mL), water (500 mL) and 0.5 M NaCl + 0.1 M NaHCO_3_ (2 L) to give three subfractions S2-1 to S2-3. The active subfraction S2-3 was applied to a silica gel vacuum liquid chromatography (VLC) column eluted with EtOAc (3 column volumes [cv]), EtOAc/MeOH (8:2, 0.1% AcOH, 3 cv), EtOAc/MeOH (1:1, 0.1% AcOH, 5 cv), and MeOH/water/acetic acid 9:1:0.3 (v/v/v, 8 cv) to yield four fractions S-2-3-1∼4. Subfraction S2-3-3 was subjected to reverse-phase CombiFlash ISCO (RediSep Rf C18, Teledyne) and eluted with Water-ACN isocratic gradients (5%, 8%, 11% and 15% acetonitrile(ACN) supplemented with 0.1% FA, 3 cv each concentration) to yield stravidin S4 (280 mg) and S5 (71 mg). Subfraction S2-3-4 was subjected to reverse-phase CombiFlash ISCO (RediSep Rf C18, Teledyne) and eluted with Water-ACN isocratic gradients (5% and 8% ACN supplemented with 0.1% FA, 3 cv each concentration) to yield stravidin S2 (59 mg) and *O*-Me-S2 (91 mg).

#### Purification of acidomycin

*Streptomyces* culture filtrate (6 L) was extracted with 5% (w/v) HP-20 (Diaion) resin twice after adjusting its pH to 4.0. The resin was eluted with 15% MeOH in water (v/v, 0.1% FA, 3.8 L) and 90% MeOH in water (v/v, 0.1% FA, 3 L) to yield two fractions A1 and A2. The fraction A2 was applied to a silica gel VLC column eluted with Hexanes/EtOAc (4:1, 3 cv), EtOAc (0.1% AcOH, 3 cv), and EtOAc/MeOH (1:1, 0.1% AcOH, 5 cv) to yield three fractions A2-1∼3.The subfraction A2-2 was loaded on reverse-phase CombiFlash ISCO (RediSep Rf C18, Teledyne) and eluted with a Water-ACN linear gradient (5-95% ACN, 0.1% FA) to give 51 fractions A2-2-1 to A2-2-51. The subfractions A2-2-26∼29 were eluted with 50% to 52% ACN, combined and reloaded on reverse-phase CombiFlash ISCO (RediSep Rf C18, Teledyne) and eluted with a Water-MeOH linear gradient (40-100% MeOH, 0.1% FA) to give 71 fractions A2-2-26-1 to A2-2-26-71. The subfractions A2-2-26-27∼30 were combined and passed through a Sephadex LH-20 column (35 mm × 500 mm, 460 mL column volume), eluting with 85% MeOH, to yield pure acidomycin (52 mg).

#### Purification of dapamycin A

*Streptomyces* culture filtrate (4 L) was loaded on a strong cation exchange (Dowex 50WX8, H^+^ form) column after adjusting its pH to 4.0. The resin was eluted with MeOH (0.1% AcOH, 3 cv), water (0.1% AcOH, 3 cv), and 2% NH_4_OH (v/v, 10 cv) to yield three fractions DA1-DA3. The fraction DA3 was applied to a silica gel VLC column eluted with EtOAc/MeOH (9:1, 0.1% AcOH, 3 cv), EtOAc/MeOH (v/v 8:2, 0.1% AcOH, 3 cv), and MeOH (0.1% AcOH, 5 cv) to yield three fractions DA3-1∼3. The subfraction DA3-3 was recrystallized in MeOH to yield white amorphous powder. The white powder was further purified with isocratic elution on reverse-phase CombiFlash ISCO (RediSep Rf C18, Teledyne) using 5% ACN (0.1% FA) as the mobile phase to yield pure dapamycin A (160 mg).

#### Purification of dapamycin B

*Streptomyces* culture filtrate (4 L) was loaded on a strong cation exchange (Dowex 50WX8, H^+^ form) column after adjusting its pH to 4.0. The resin was eluted with MeOH (0.1% AcOH, 3 cv), water (0.1% AcOH, 3 cv), and 2% NH_4_OH (v/v, 10 cv) to give three fractions DB1-DB3. The fraction DB3 was applied to a silica gel VLC column eluted with EtOAc/MeOH (9:1, 0.1% AcOH, 3 cv), EtOAc/MeOH (v/v 8:2, 0.1% AcOH, 3 cv), and MeOH (0.1% AcOH, 5 cv) to yield three fractions DB3-1∼3. The subfraction DB3-3 was recrystallized in MeOH to yield white amorphous powder. The white powder was further purified with isocratic elution on reverse-phase CombiFlash ISCO (RediSep Rf C18, Teledyne) using 5% ACN (0.1% FA) as mobile phase to yield pure dapamycin B (122 mg).

#### Purification of α-Me-KAPA

*Streptomyces* culture filtrate (6 L) was extracted with 5% (w/v) HP-20 (Diaion) resin four times after adjusting its pH to 4.0. The resin was eluted with 15% MeOH in water (v/v, 0.1% FA, 3.8 L) and 90% MeOH in water (v/v, 0.1% FA, 3 L) to yield two fractions, KA1 and KA2. The active fraction KA1 was passed through a Sephadex LH-20 column (35 mm × 500 mm) and eluted with 85% MeOH to yield 60 subfractions (KA-1-1 to KA-1-60, 10 mL each). The active subfractions KA1-25 to 29 were combined, subjected to reverse-phase CombiFlash ISCO (RediSep Rf C18, Teledyne) and eluted with a linear gradient of 5-20% ACN (0.1% FA) to give 68 fractions KA1-25-1 to KA1-25-68. The active subfractions, eluted with 7% to 12% ACN, were combined and reloaded on reverse-phase CombiFlash ISCO (RediSep Rf C18, Teledyne) and eluted with a linear gradient of 5-30% ACN (0.1% TFA) to yield pure α-Me-KAPA (67 mg).

#### Purification of 2,5-di-(2-methylhexanoic acyl)-3-methylimidazole and 2,5-dimethyl-3,6-di-(2-methylhexanoic acyl)pyrazine

*Streptomyces* culture filtrate (6 L) was extracted with 5% (w/v) HP-20 (Diaion) resin twice after adjusting its pH to 4.0. The resin was eluted with 15% MeOH in water (v/v, 0.1% FA, 2 L) and 100% MeOH (0.1% FA, 3 L) to yield two fractions, KB1 and KB2. The fraction KB-2 was applied to a silica gel VLC column eluted with Hexanes/EtOAc (4:1, 3 cv), EtOAc (0.1% AcOH, 3 cv), EtOAc/MeOH (1:1, 0.1% AcOH, 5 cv) and MeOH (0.1% AcOH, 4 cv) to yield four fractions KB2-1∼4. The subfraction KB2-2 was subjected to reverse-phase CombiFlash ISCO (RediSep Rf C18, Teledyne) and eluted with a linear gradient of 50-100% MeOH (0.1% FA) to give 68 fractions KB2-2-1∼68. The subfractions KB2-2-23∼29 were combined and reloaded on reverse-phase CombiFlash ISCO (RediSep Rf C18, Teledyne) and eluted with a linear gradient of 40-50% ACN (0.1% FA) to yield pure 2,5-dimethyl-3,6-di-(2-methylhexanoic acyl)pyrazine (38 mg). The subfraction KB2-3 was subjected to reverse-phase CombiFlash ISCO (RediSep Rf C18, Teledyne) and eluted with a linear gradient of 30-100% MeOH (0.1% FA) to give 64 fractions KB2-3-1∼64. The subfractions KB2-3-24∼41 were combined and reloaded on reverse-phase CombiFlash ISCO (RediSep Rf C18, Teledyne) and eluted with a linear gradient of 15-55% ACN (0.1% FA) to yield pure 2,5-di-(2-methylhexanoic acyl)-3-methylimidazole (13 mg).

### Structural characterization of NPs

HR-MS of all compounds were recorded on an Agilent 6550 iFunnel Q-TOF mass spectrometry equipped with an inline Agilent 1290 HPLC system using electrospray ionization in positive mode with a mass scan range of 50-1000 m/z. Tandem MS/MS was performed on the same instrument using a step gradient of collision-induced dissociation (CID) energy of 5, 10, 15, 20, and 25 eV. 1-/2-D NMR experiments were performed on a Bruker AVIII 700 MHz instrument equipped with a cryoprobe. All NMR data are shown in **Figures S18-S67** and summarized in **Tables S2-S7**.

### Target gene deletion in WAC05950

Homologous recombination was adopted to delete genes from the chromosome of WAC05950^52^. bioAB-arm-up-F/R and bioAB-arm-dw-F/R primers were used to amplify the homologous arms from the genomic DNA of WAC05950, and then assembled into *BamH*I/*Kpn*I linearized pSUC01 plasmid through Gibson assembly, generating pSUC01-bioAB plasmid. Error-free plasmid was introduced into *E. coli* ET12567/pUZ8002 and then shuttled into WAC05950 through *E. coli*-*Streptomyces* interspecies biparental mating. WAC05950 exconjugants carrying pSUC01-bioAB plasmid integrated into the chromosome through single crossover were selected using apramycin and streaked onto SFM agar medium supplemented with apramycin (50 μg/mL). Apramycin-resistant WAC05950 exconjugant was then inoculated into 1 mL antibiotic-free TSBY medium and grown at 30℃ overnight, followed by successive passage into antibiotic-free TSBY medium for 2-4 generations. The overnight culture was then plated onto antibiotic-free SFM agar medium (0.05 μL overnight culture) for isolating single colonies. *Streptomyces* single colonies without blue pigment production were picked and streaked onto both apramycin-supplemented and antibiotic-free SFM media to select for double crossover mutants. Apramycin-sensitive and blue pigment-free colonies were screened by PCR using ΔbioAB-dF/dR primers to identify WAC05950ΔbioAB mutant. WAC05950ΔaciB was generated using an identical procedure except for the primers used for cloning, sequencing, and screening.

### MIC determination assays

Overnight liquid cultures of *E. coli*, *K. pneumoniae* and *P. aeruginosa* strains were inoculated with a single colony from a freshly streaked agar plate of the corresponding bacterial strain and cultured in MOPS minimal medium with 0.4% glucose. The bacterial culture was prepared by washing 1 mL of overnight culture in PBS pH 7.4 (×3), then diluted to OD_600_ of 0.35 and used to inoculate 1:1000 in MOPS minimal medium with 0.4% glucose in 96-well plates. Compounds were added to 96-well plates in two-fold serial dilutions. The plates were incubated at 37℃ for 18 h at 250 rpm, then OD_600_ was measured. The MIC values were determined to be the minimal concentration that fully inhibits bacterial growth. *A. baumannii* strain was grown in identical conditions, except MOPS minimal medium was supplemented with 0.6% succinate instead of glucose. *M. tuberculosis* and *M. bovis* were cultured from working stocks in biotin-free 7H9 media (Middlebrook 7H9 containing 0.2% glycerol, 10% OADC, and 0.05% Tween-80, but lacking biotin), washed once in saline, and cultured an additional 2 days in biotin-free 7H9 media. *M. smegmatis*, *M. fortuitum* and *M. abscessus* were grown on LB plates before culturing in biotin-free 7H9 media. All cultures were diluted to a final CFU/mL of ∼5 × 10^5^ cells/mL. CFU totals were confirmed to be within the range of 3-8 × 10^5^ cells/mL by plating of dilutions of the inoculum on Middlebrook 7H10 (Becton Dickinson) agar, containing 0.5% glycerol and 10% OADC. Plates were incubated stationary at 37℃ for 3 days (*M. fortuitum* and *M. abscessus*) or 7 days (*M. tuberculosis* and *M. bovis*), then OD_600_ was measured. For *M. smegmatis*, plates were incubated shaking at 37℃ for 32 h. *M. tuberculosis* and *M. bovis* were incubated with 30 µg/mL resazurin for one additional day to identify any effect on growth where compound solubility was limiting. Two biological replicates were performed for each experiment.

### Biotin operon promoter assay

*E. coli* cells harbouring the pUA66-*bioB* plasmid^53^ were prepared in the same way as for MIC experiments. Kanamycin (50 µg/mL) was added to the media to maintain plasmid selection. Cells were incubated while shaking at 37°C in a 96-well plate with a serially diluted compound. The GFP fluorescence signal, corresponding to *bioB* promoter activation, was measured at 535 nm with excitation at 489 nm using a BioTek Synergy H1 plate reader.

### Checkerboard microdilution assays

Bacteria were prepared in the same way as for MIC experiments. Each compound was serially diluted two-fold across eight concentrations in an 8 by 8 matrix on a 96-well plate. The fraction inhibitory concentration (FIC) for each compound was determined as a fraction of its MIC divided by the MIC in combination. The FIC indices (FICI) were calculated as the sum of the two FIC values for each compound in the combination. FICI values ≤0.5 were interpreted as synergistic, >0.5 and ≤1.0 as additive, >1 and ≤4.0 as indifferent, and >4.0 as antagonistic. Three biological replicates were performed for each combination.

### Expression and purification of recombinant proteins

Constructs and strains used for the recombinant expression of *E. coli* BioA, BioD, BirA, and AccB are outlined in **Table S11**. For BioA, the same plasmid construct and purification protocol were used as previously described^19^. In brief, BioA was expressed in *E. coli* AG1-pCA24N-bioA (JW0757) from the ASKA library^54^ and grown in 2 L of LB medium supplemented with chloramphenicol (80 μg/ml) at 37°C with shaking. Protein expression was induced at an OD_600_ of 0.6 by the addition of 0.1 mM IPTG and incubated for an additional 3 hours. Cells were harvested, resuspended and washed with a 0.85% saline solution, pelleted and stored at −20°C. The cells were resuspended in 25 mL lysis buffer (50 mM HEPES, pH 8, 500 mM NaCl, 100 μM PLP, 50 mM imidazole, 0.5 mg DNase, 0.5 mg RNase and protease inhibitor cocktail (Roche)). After cell disruption with a high-pressure homogenizer, the cell lysate was cleared by centrifugation at 25,000 × g for 30 min at 4°C. For protein purification, the cleared lysate was loaded onto a nickel-chelating 1-mL HiTrap affinity column (GE Healthcare, Mississauga, Canada), washed with buffer A (50 mM HEPES, pH 8, 500 mM NaCl, 100 μM PLP, 50 mM imidazole) and eluted with a linear gradient of 50–400 mM imidazole. Fractions were analyzed via SDS-PAGE for the presence of pure BioA, pooled, and desalted through a HiPrep 26/10 desalting column (GE) against the final storage buffer (50 mM HEPES, pH 8, 10% glycerol).

For the expression of BioD, BirA, and AccB, 1% of overnight cultures of respective expression strains (**Table S11**) were used to inoculate 1 L of 2x YT medium supplemented with the appropriate antibiotic and cultured at 37°C with shaking. Protein expression was induced in mid-log phase (OD_600_ = 0.6) with 0.5 mM IPTG for 4 hours at 30°C, and cells were harvested at 6,000 × g for 15 min. Protein purification buffer conditions were selected based on published literature^55–57^ with slight modifications and summarized in **Table S12**. After harvesting, the cell pellet was resuspended in 15 mL ice-cold buffer A and stored at -20°C.

All subsequent protein purification steps were carried out on ice or at 4°C. The cells were thawed, supplemented with lysozyme (250 µg/mL), DNase (50 µg/mL), and RNase (10 µg/mL) and incubated for 30 min. After cell disruption in a high-pressure homogenizer (20 kPa), the lysate was cleared by centrifugation at 25,000 × g for 30 min and subjected to Ni^2+^-affinity gravity flow chromatography. The soluble lysate was applied to 1 mL Ni-NTA-agarose resin (Qiagen), incubated for 1 h and transferred onto a column. After two consecutive wash steps with 15 mL buffer A and 15 mL buffer W to remove weakly bound material, His-tagged recombinant proteins were eluted with buffer E. After SDS-gel electrophoresis, protein-containing fractions were pooled and exchanged into storage buffer overnight at 4°C using 10K MWCO SnakeSkin tubing (Thermo, Cat. No. 68100). For BirA, the SUMO tag was cleaved post-purification by overnight incubation with 0.13 mg/mL Ulp-1 at 4°C. After the SUMO-cleavage step, protein mixtures were once more subjected to Ni^2+^-affinity chromatography to remove non-cleaved recombinant protein and the SUMO protease. The flow-through was collected, concentrated, desalted, and stored at –80°C in the above-mentioned storage buffer. Proteins were concentrated using Vivaspin 20 (MWCO 10,000, Cytiva, Cat. No. 28932360). Protein concentrations were determined using a NanoDrop ND-1000 spectrophotometer and extinction coefficients calculated via ProtParam (Expasy): BioA (63,535), BioD (38,055), BirA (47,440), AccB (4,470).

### BioA enzymatic reaction

The reaction catalyzed by BioA was carried out as described previously^58^, with slight modifications. The enzymatic reaction mixture comprised 100 mM HEPES (pH 8.5), 1 mM dithiothreitol, 25 μM KAPA and 0.5 mM SAM. The reaction was initiated by the addition of 200 nM BioA after 2 min of preincubation of the substrate mixture at 37°C, followed by a 20 min incubation. To stop the reaction, 150 μL of derivatizing solution comprising o-phthaldialdehyde/2-mercaptoethanol reagent, 0.26 M sodium borate buffer pH 9.4 and ethanol was added. The reaction was incubated at room temperature for 2 h, and the fluorescent signal, corresponding to derivatized DAPA, was measured at 470 nm with excitation at 410 nm on BioTek Synergy Neo2.

### BioD enzymatic reaction

BioD activity was evaluated using a modified PK/LDH-coupled assay^59^. The enzymatic mixture comprised 100 mM K-HEPES (pH 7.5), 20 mM NaHCO_3_, 10 mM MgCl_2_, 5 mM KCl, 50 µM ATP, 50 µM DAPA, 5 mM PEP, 0.2 mM NADH, 1:50 PK-LDH mix (Sigma-Aldrich, P0294) and and varying concentrations of dapamycin A or B. Reactions were initiated by adding 5 µM BioD, incubated at 37°C, and monitored kinetically at 340 nm for 30 min using a BioTek Synergy Neo2 plate reader to measure NADH consumption. Absorbance traces at 340 nm were fitted to linear regressions, and the resulting slopes were used to calculate reaction rates.

### BioA-BioD coupled enzymatic reaction

For a coupled BioA-BioD reaction, 100 µM of α-Me-KAPA was incubated with 0.1 µM of BioA, 1.5 µM of BioD, 20 mM NaHCO_3_, 200 µM PLP, 1 mM SAM, 100 µM ATP for 1 h at 37°C in a buffer containing 100 mM HEPES (pH 8.5), 5 mM MgCl_2_, 5 mM KCl, 1 mM DTT. The reaction was quenched with ice-cold water before LC ESI–MS analysis. 2 μL of each sample was analyzed on an Agilent 6546 LC/Q-TOF mass spectrometry equipped with an inline Agilent 1290 HPLC system using Poroshell 120 EC-C18 column (120 Å, 1.9 μm, 2.1 mm × 100 mm). HPLC gradient used was: t = 0-2 min, 2% B; t = 17 min, 20% B; t = 22-30 min, 98% B; t = 31-37 min, 2% B; at a flow rate of 0.27 mL/min, 40℃ with H_2_O (0.1% formic acid [FA], A) and ACN (0.1% FA, B) as mobile phases.

### BirA enzymatic reaction and AccB intact protein LC ESI–MS

BirA (1 µM) was incubated with 1 mM ATP and 100 µM of biotin or α-Me-biotin for 1 h at 37°C in a buffer containing 10 mM Tris-HCl (pH 8.0), 2.5 mM MgCl_2_ and 5 mM KCl. The reaction was quenched with ice-cold water before LC ESI–MS analysis. Samples were analyzed using the same method used for the BioA-BioD coupled reaction.

For a coupled reaction, AccB (10 µM) was added to the BirA enzymatic reaction and incubated for 3 h. To evaluate biotinylation or methyl-biotinylation, intact protein LC ESI–MS was performed on an Agilent 6546 LC/Q-TOF in positive ion mode with a ZORBAX StableBond 300 C3 column (300 Å, 3.5 μm, 3.0 × 150 mm). HPLC gradient used was: t = 0-1 min, 5% B; t = 21 min, 60% B; t = 26-30 min, 97% B; t = 30-31 min, 5% B; at a flow rate of 0.4 mL/min, 65℃ with H_2_O (0.1% formic acid [FA], A) and ACN (0.1% FA, B) as mobile phases. Spectra were deconvoluted, and figures were generated using UniDec software^60^.

### Overexpression system in *M. smegmatis*

We adapted a theophylline-sensitive riboswitch to overexpress FLAG-tagged biotin biosynthesis genes in *M. smegmatis* ^61^. A riboswitch cassette (riboBlock, **Table S8**) containing a strong constitutive promoter (A37TG-conN18)^62^, theophylline-responsive riboswitch^61^, and C-terminal 3×FLAG tag was ordered as a gBlock from IDT. The cassette was introduced by Gibson assembly into the backbone of pMV306hsp+LuxG13, removing the luciferase genes^63^, using primers riboBlock-F/R and MV306op-F/R to generate plasmid pRibo306. BioF, BioA, BioD, and BioB from *M. smegmatis* mc^2^155 were amplified using primers bioD-Gib-F/R, bioF-Gib-F/R, bioA-Gib-F/R, bioB-Gib-F/R. Biotin enzyme gene amplicons were assembled with theophylline-responsive vector opened with primers riboOp-F/R using Gibson assembly. Plasmids were validated by Sanger sequencing using primers 306-F/R at the McMaster Genomics Facility.

Plasmids were introduced into *M. smegmatis* mc^2^155 by electroporation using standard techniques, and transformants were selected with 50 µg/mL kanamycin. Each transformed strain was grown for 16 h in biotin-free 7H9 + OADC + Tween media at 37°C in the presence or absence of 4 mM theophylline (Sigma). Saturated cultures were washed 3× with sterile saline and resuspended in biotin-free 7H9 + OADC + Tween media with the corresponding theophylline concentration (0 or 4 mM). Sensitivity to various biotin antimetabolites was determined by an MIC assay after 2-3 days at 37°C.

### Heterologous expression of *M. smegmatis* biotin enzymes in *E. coli*

Kanamycin resistance markers from Keio deletion strains of *bioF*, *bioA*, *bioD*, and *bioB* were removed by standard methods^64^. Markerless strains were transformed with either empty vector, pRibo306, or with the cognate missing enzyme from *M. smegmatis*, pRibo306-BioF, BioA, BioB or BioD. Transformed cells were cultured in MOPS minimal media containing 0.1 ng/mL biotin at 37°C. Saturated cultures were washed 3× in sterile saline and resuspended to a final OD_600_ of 0.25 in saline. 2 mL of a 10-fold serial dilution series in saline were spotted on M9 minimal agar alone or containing either 2 mM theophylline or 4 ng/mL of biotin and cultured at 37°C for 1 day before imaging.

### Biotransformation assays of dapamycin B

*E. coli* BW25113 wild-type and ΔdppC strains were cultured following the same protocol as for MIC assays. Cultures were incubated with dapamycin B (16 µg/mL) for 24 h, after which supernatants were collected and analyzed by LC–MS using the method described for the BioA–BioD coupled assay. To quantify dapamycin A and B in the supernatants, standard curves were generated with purified compounds, and concentrations were calculated using Agilent MassHunter Quantitative Analysis software.

To investigate downstream biotransformations of dapamycins, isotope-labeled MOPS minimal media were prepared containing either ^13^C-glucose/^14^N-ammonium sulfate, ^12^C-glucose/^15^N-ammonium sulfate and ^13^C-glucose/^15^N-ammonium sulfate. Cultures were incubated with dapamycin B (64 µg/mL) for 24 h under the same conditions, and supernatants were collected and analyzed by LC–MS as above. In isotope-labeling experiments, qualitative LC–MS analysis was performed to trace incorporation of ^13^C and/or ^15^N into dapamycin-derived metabolites.

### Cell extraction and detection of α-Me-KAPA derivatives

*E. coli* BW25113 cells were prepared following the same protocol as for the MIC experiments. After washing in PBS, 250 mL of fresh MOPS minimal medium was inoculated at a 1:50 dilution and incubated at 37°C. When the OD_600_ reached 0.3–0.4, α-Me-KAPA was added to a final concentration of 4 µg/mL, and the cells were incubated for an additional 4 hours. Following incubation, the cells were pelleted by centrifugation, resuspended in 4 mL of PBS, and split evenly into four tubes. To extract α-Me-KAPA derivatives from the cells, a modified version of a previously described protocol for analyzing bacterial coenzyme A thioesters was used^65^. To each tube, 1 mL of methanol and 1 g of 0.1 mm acid-washed glass beads (Sigma-Aldrich) were added. The tubes were then shock-frozen and stored at -80°C overnight. The next day, the tubes were homogenized using a Qiagen TissueLyser III for 5 min at 30 Hz. After homogenization, the samples were centrifuged at 21,100 × g for 5 min at 4°C, and the supernatant was transferred to a clean tube. This process was repeated twice more. The supernatants from each condition were then pooled, diluted 1:2 with water, frozen, and lyophilized overnight. The lyophilized samples were resuspended in 600 µL of water, vortexed, and filtered through a 0.22-µm cellulose acetate Spin-X filter (Costar) by centrifugation at 14,000 × g for 10 min. Finally, 5 µL of each sample was injected into the LC-MS system and analyzed using the same method used for BioA-BioD couple enzyme reactions.

### *E. coli* C0244 systemic mouse infection model

All mouse experiments were performed in the Central Animal Facility at McMaster University under animal use protocol 20-12-43 as approved by the Animal Research Ethics Board. Six-to ten-week-old female CD-1 mice were used for all experiments. Systemic infection with *E. coli* C0244 was established in an immunocompetent CD-1 mouse. Mice were pre-treated with 0.2 mg/kg of streptavidin dissolved in sterile water in accordance with a previous study^21^. After 1 h, mice were infected intraperitoneally with ∼5×10^7^ CFU of bacteria suspended in 5% porcine gastric mucin PBS solution. 1 h post-infection, mice were treated intraperitoneally with vehicle or a single dose (50 mg/kg) of acidomycin, stravidin S2, dapamycin B or α-Me-KAPA. The vehicle for acidomycin was 5% DMSO, 30% PEG300 and 65% sterile water. The vehicle for the rest of the compounds was sterile water. The experimental endpoint was defined as 6 h after infection. Spleen, kidneys and liver were collected and homogenized in PBS. The blood was collected from the facial vein. The resulting samples were diluted and plated on LB agar to determine the bacterial load.

## Acknowledgements

This research was generously supported by funds to E.D.B., including Tier 1 Canada Research Chair award, a Foundation Grant from the Canadian Institutes of Health Research (CIHR) (FRN 143215) and a grant from the Ontario Research Fund (RE09-047). G.D.W. is supported by CIHR grants FDN 148463 and PJT190298. Tianjin Synthetic Biotechnology Innovation Capacity Improvement Project (TSBICIP-CXRC-065, TSBICIPCXRC-076 to M.X.), and the Hundred Talents Program of Chinese Academy of Sciences (to M.X.). M.M.T. was supported by a CIHR Canada Graduate Scholarship (CGS-D). The funders had no role in study design, data collection and analysis, decision to publish or preparation of the manuscript. We would also like to thank Prof. Wenjun Guan for gifting pSUC01 plasmid.

## Author contributions

R.G. and M.X. conceived the research, designed and carried out experiments and data analysis, and wrote the manuscript. W.W. performed compound purifications. M.A.C. acquired MIC data, designed plasmids for protein overexpression and assisted with research direction. D.H. assisted with the phylogenetic tree construction and data interpretation. J.P.D. assisted with protein purifications. M.M.T and L.A.C. assisted in the animal experiments. M.G. assisted with biotinylation assays and CORASON analysis. D.S. assisted with protein purifications and biotinylation assays. K.R. assisted with research direction. K.K. assisted with compound purifications. A.W. assisted with genomic preparations and cloning. E.D.B. and G.D.W. conceived the research and assisted with data interpretation and manuscript editing.

## Competing interests

The authors declare no competing interests.

**Correspondence and requests for materials** should be addressed to Eric D. Brown and Gerard D. Wright.

