## Supplemental Figures and Tables for "A ubiquitous *Streptomyces* biosynthetic megacluster encodes an arsenal of synergistic biotin-targeting antibiotics"

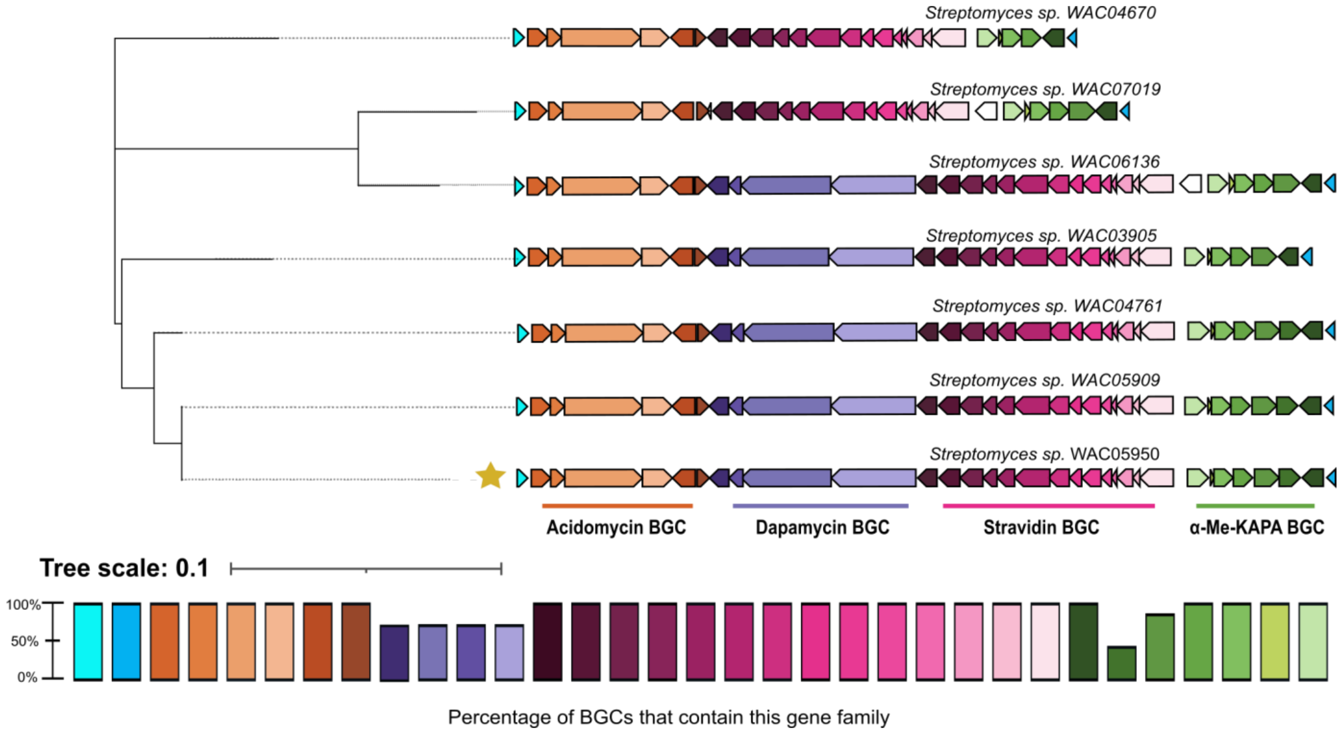


**Figure S1. CORASON-based phylogenetic analysis of megacluster-containing *Streptomyces* strains from the WAC collection.** The tree was generated using AciB, a core enzyme from the acidomycin biosynthetic gene cluster, to compare homologous clusters across strains. *Streptomyces* sp. WAC05950 was used as the reference genome (yellow star).

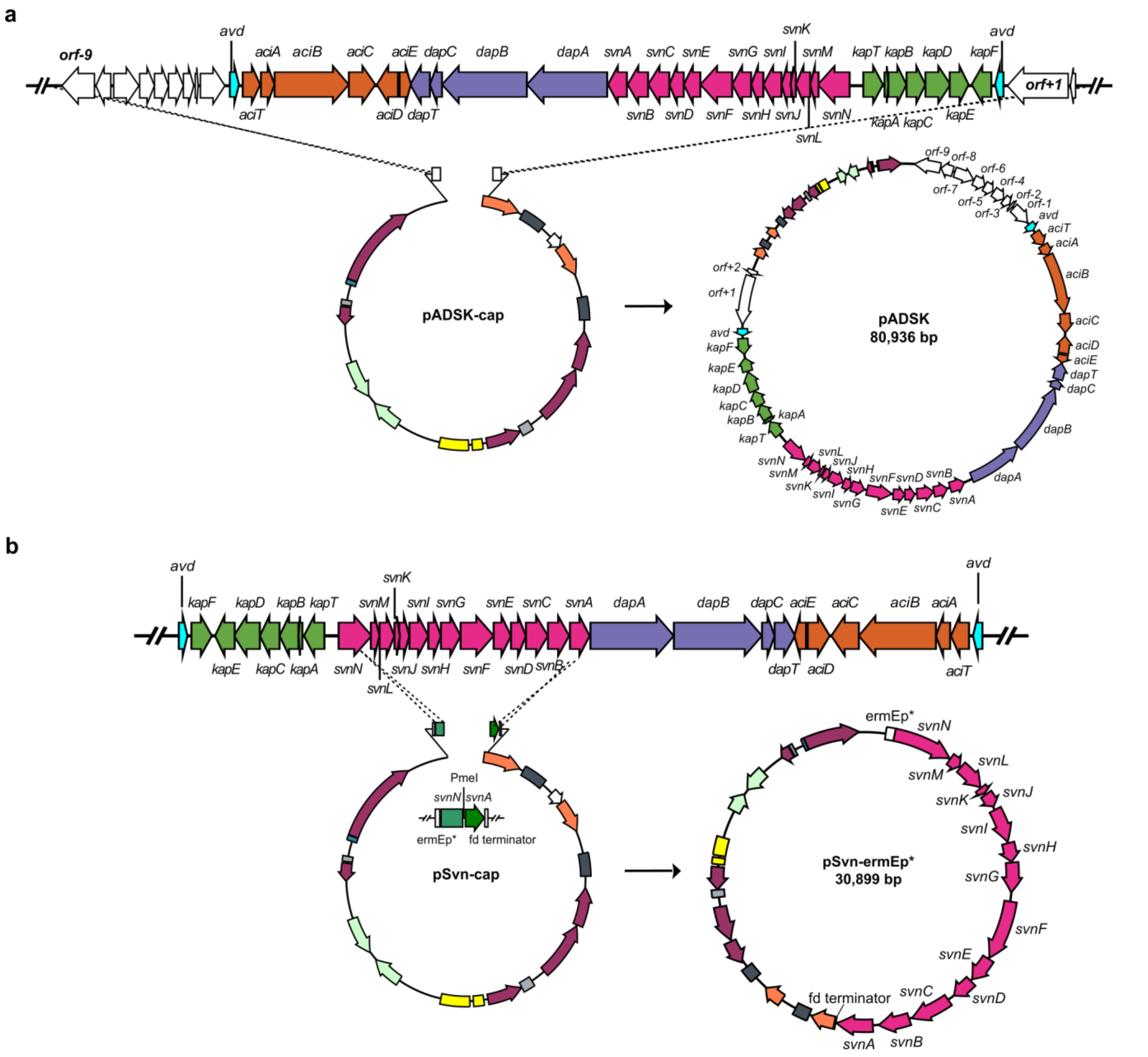


**Figure S2. TAR cloning of the megacluster and *svn* BGC. a**, Schematic representation of cloning the megacluster from the genome of WAC05950 into pCGW, resulting in pADSK. **b**, Schematic representation of subcloning and refactoring the *svn* BGC from pADSK using TAR. ermEp* promoter was introduced to overexpress the *svn* BGC, and fd terminator was introduced after *svnA* for transcription termination.

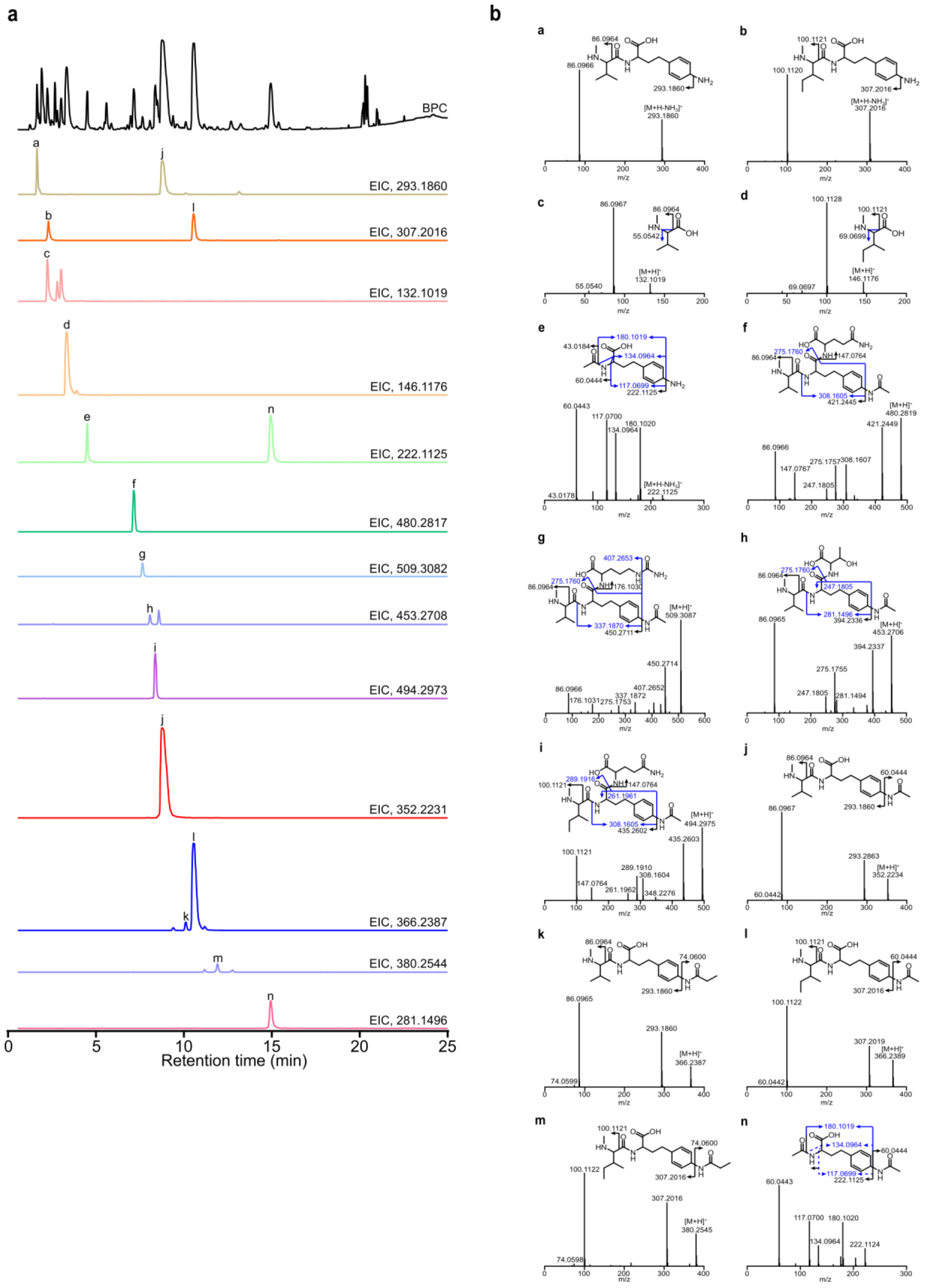


**Figure S3. MS/MS determination of stravidin analogs. a,** EICs of stravidins produced by *S. coelicolor* M1154/pADSK. **b,** MS/MS fragmentation analysis of stravidins (CID=5 eV). The black, blue, and blue dashed arrows indicate daughter ions produced by single, double, and triple bond breakages in the compounds. CID stands for collision-induced dissociation. The blue arrows indicate the daughter ions produced by breaking two bonds in the compounds, and the blue dashed arrows indicate the daughter ions produced by breaking three bonds in the compounds.

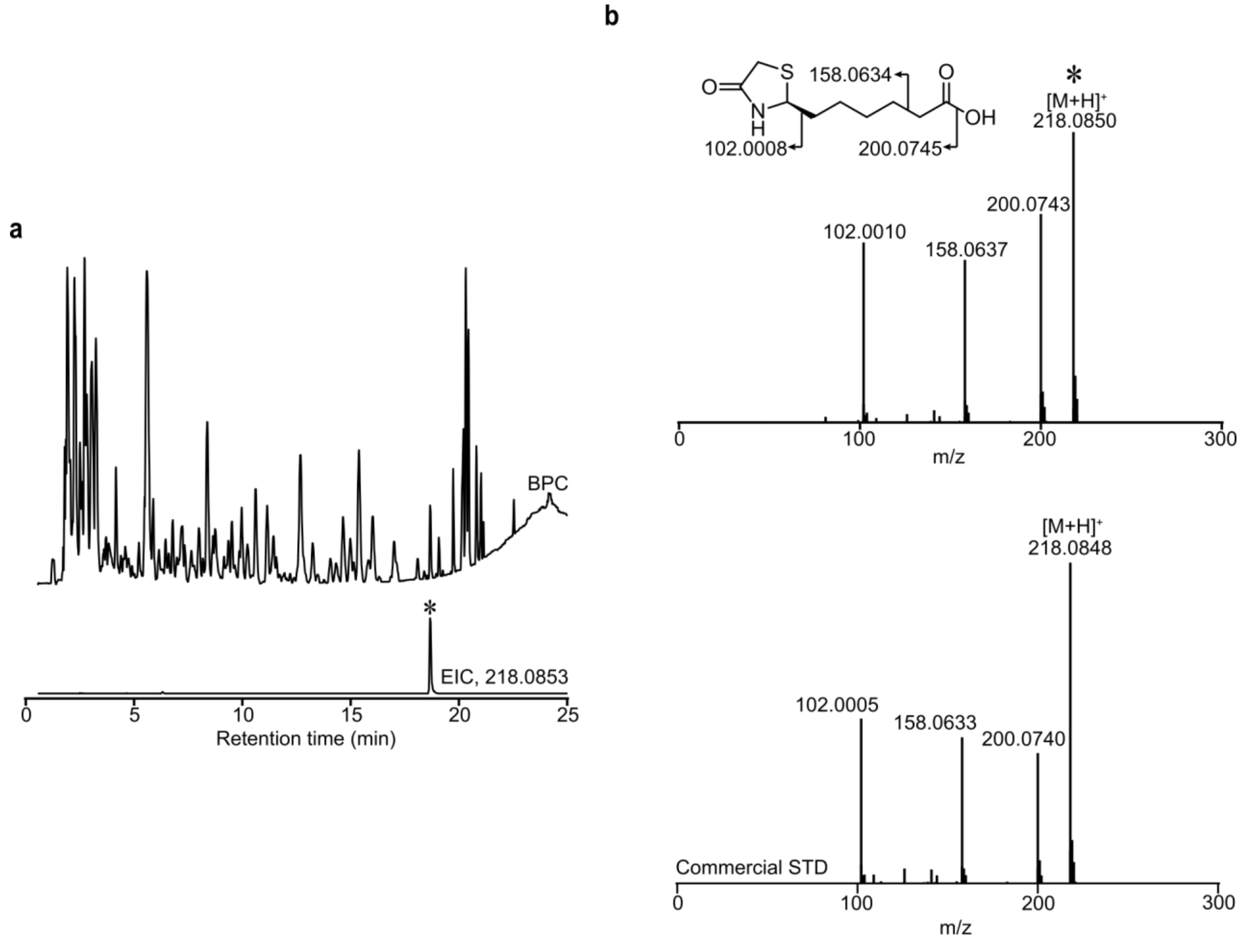


**Figure S4. MS/MS characterization of acidomycin. a,** EIC of acidomycin produced by *S. coelicolor* M1154/pADSK. The acidomycin peak is labelled by an asterisk (∗). **b,** MS/MS fragmentation of acidomycin peak in the culture media and a commercial standard (CID = 5 eV).

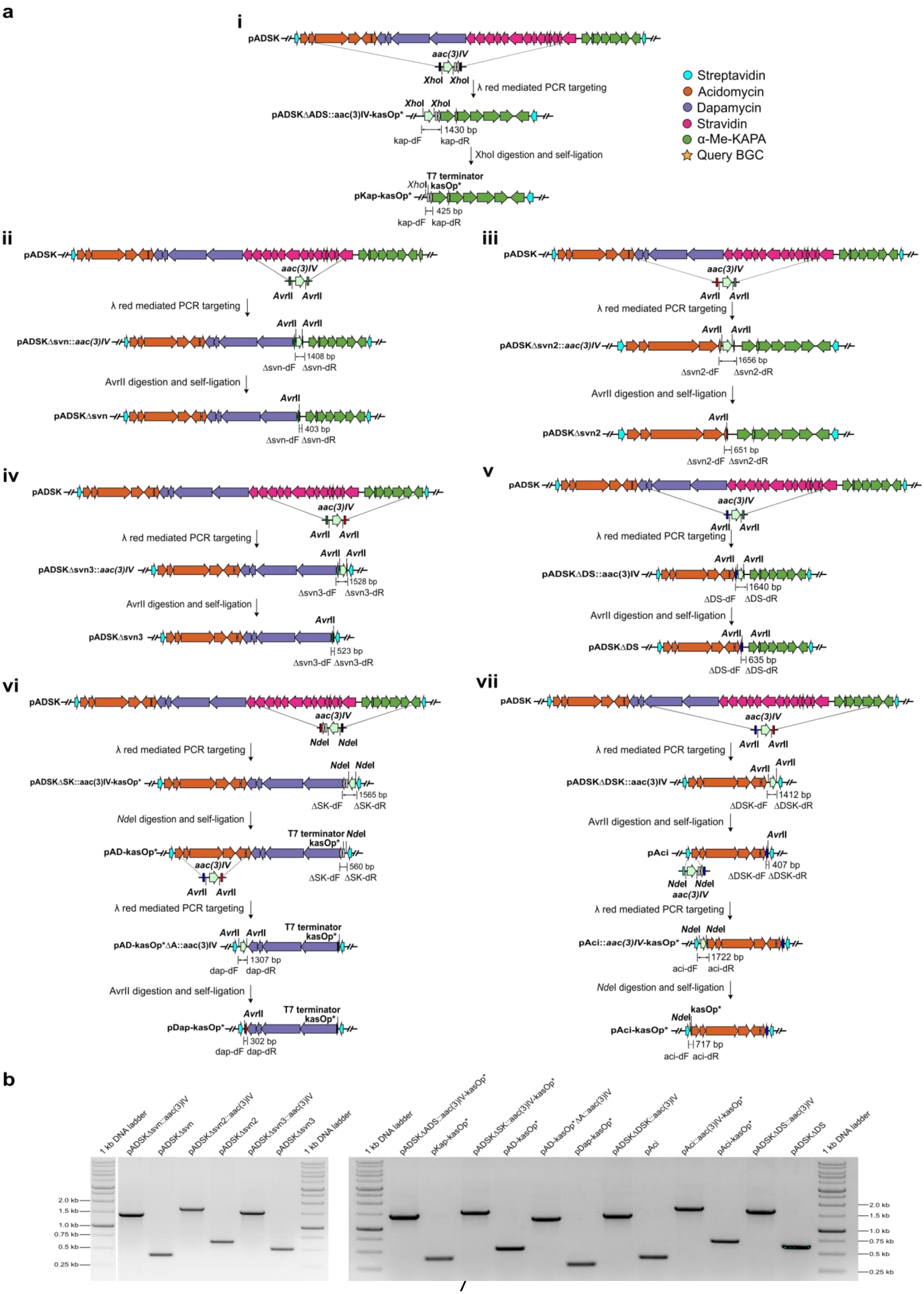

a
i
pADSK
aac(3)IV
XhoI
XhoI
λ red mediated PCR targeting
pADSKΔADS::aac(3)IV-kasOp*
1430 bp
kap-dF
kap-dR
XhoI digestion and self-ligation
T7 terminator
kasOp*
pKap-kasOp*
425 bp
Streptavidin
Acidomycin
Dapamycin
Stravidin
α-Me-KAPA
Query BGC
ii
pADSKΔsvn::aac(3)IV
1408 bp
Δsvn-dF
Δsvn-dR
AvrII digestion and self-ligation
pADSKΔsvn
403 bp
iii
pADSKΔsvn2::aac(3)IV
1656 bp
Δsvn2-dF
Δsvn2-dR
pADSKΔsvn2
651 bp
iv
pADSKΔsvn3::aac(3)IV
1528 bp
Δsvn3-dF
Δsvn3-dR
pADSKΔsvn3
523 bp
v
pADSKΔDS::aac(3)IV
1640 bp
ΔDS-dF
ΔDS-dR
pADSKΔDS
635 bp
vi
NdeI
pADSKΔSK::aac(3)IV-kasOp*
1565 bp
ΔSK-dF
ΔSK-dR
NdeI digestion and self-ligation
pAD-kasOp*
560 bp
pAD-kasOp*ΔA::aac(3)IV
1307 bp
dap-dF
dap-dR
pDap-kasOp*
302 bp
vii
pADSKΔDSK::aac(3)IV
1412 bp
ΔDSK-dF
ΔDSK-dR
pAci
407 bp
pAci::aac(3)IV-kasOp*
1722 bp
aci-dF
aci-dR
pAci-kasOp*
717 bp
b
1 kb DNA ladder
pADSKΔsvn::aac(3)IV
pADSKΔsvn
pADSKΔsvn2::aac(3)IV
pADSKΔsvn2
pADSKΔsvn3::aac(3)IV
pADSKΔsvn3
pADSKΔADS::aac(3)IV-kasOp*
pKap-kasOp*
pADSKΔSK::aac(3)IV-kasOp*
pAD-kasOp*
pAD-kasOp*ΔA::aac(3)IV
pDap-kasOp*
pADSKΔDSK::aac(3)IV
pAci
pAci::aac(3)IV-kasOp*
pAci-kasOp*
pADSKΔDS::aac(3)IV
pADSKΔDS
2.0 kb
1.5 kb
1.0 kb
0.75 kb
0.5 kb
0.25 kb

**Figure S5. Engineering and refactoring of subclusters within the anti-biotin megacluster. a**, Schematic of pADSK plasmid engineering and refactoring through λ-red mediated PCR targeting by removing DNA regions covering *aciT-svnN* (i), *svnA-svnN* (ii), *aciE-svnN* (iii), *svnA-kapF* (iv), *dapT-svnN* (v), *aciT-aciE* and *svnA-kapF* (vi), and *dapT-kapF* (vii). **b**, DNA agarose gel of the PCR products amplified from pADSK and its derivative plasmids.

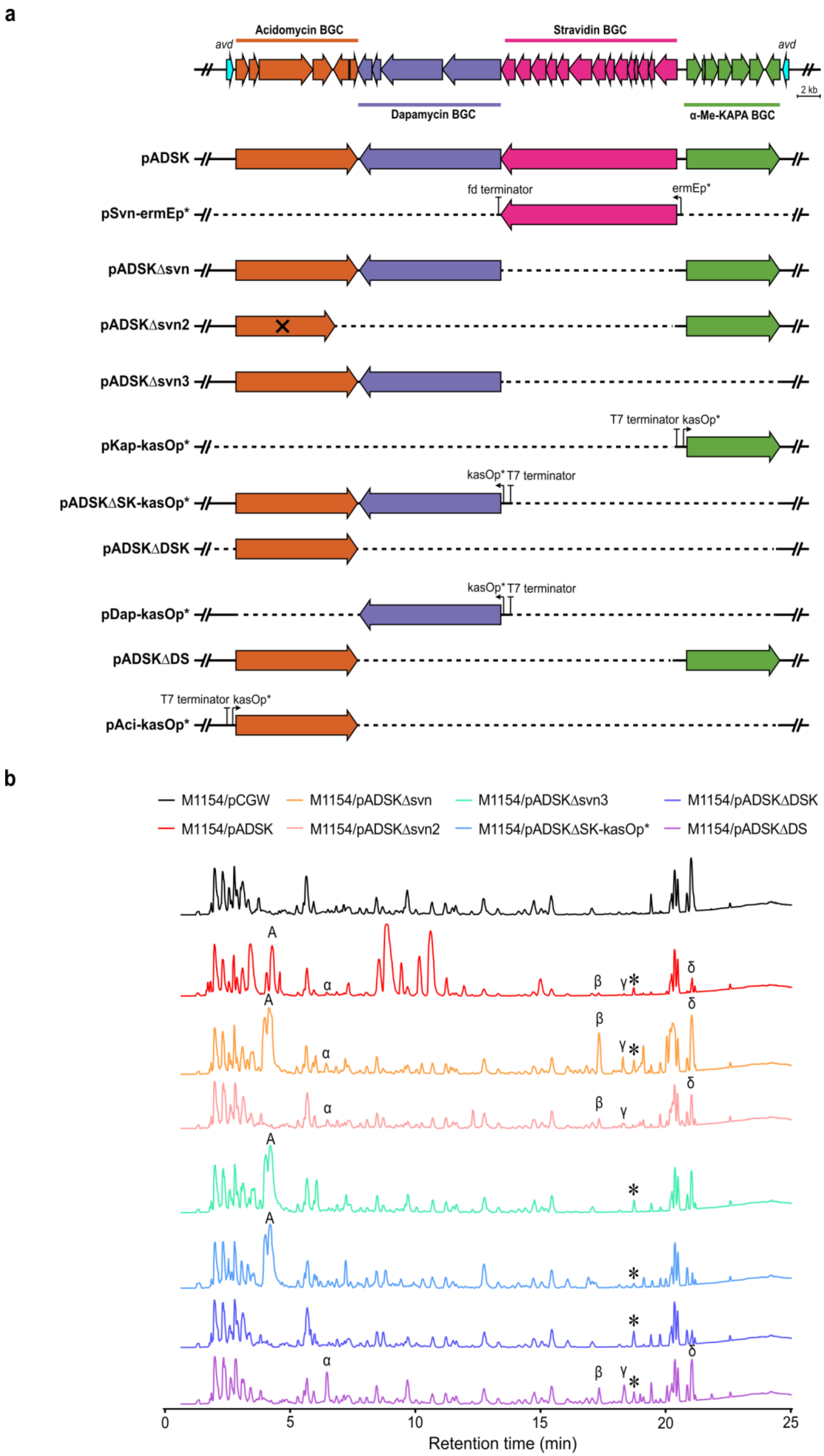
a
Acidomycin BGC
Stravidin BGC
avd
avd
2 kb
Dapamycin BGC
α-Me-KAPA BGC
pADSK
fd terminator
ermEp*
pSvn-ermEp*
pADSKΔsvn
pADSKΔsvn2
pADSKΔsvn3
T7 terminator kasOp*
pKap-kasOp*
kasOp* T7 terminator
pADSKΔSK-kasOp*
pADSKΔDSK
kasOp* T7 terminator
pDap-kasOp*
pADSKΔDS
T7 terminator kasOp*
pAci-kasOp*
b
M1154/pCGW
M1154/pADSKΔsvn
M1154/pADSKΔsvn3
M1154/pADSKΔDSK
M1154/pADSK
M1154/pADSKΔsvn2
M1154/pADSKΔSK-kasOp*
M1154/pADSKΔDS
A
α
β
γ
δ
0
5
10
15
20
25
Retention time (min)

**Figure S6. Metabolic profiling of truncated version of the anti-biotin megacluster. a,** Schematic of the engineered and refactored pADSK plasmids. Each subcluster was colored accordingly to clarify the deletion region. The dashed lines indicate the deleted regions from the pADSK plasmid. The ✕ in pADSKΔsvn2 indicates the presence of an impaired *aci* BGC. **b,** Base peak chromatogram (BPC) profile of *S. coelicolor* M1154 strain expressing pADSK derivatives. Each compound was labelled according to the BPC as: A: dapamycin A; α: α-Me-KAPA; β: 2,5-di-(2-methylhexanoic acyl)-3-methylimidazole; γ: N-Acetyl-α-Me-KAPA; δ: 2,5-dimethyl-3,6-di-(2-methylhexanoic acyl)pyrazine; and ∗: acidomycin.

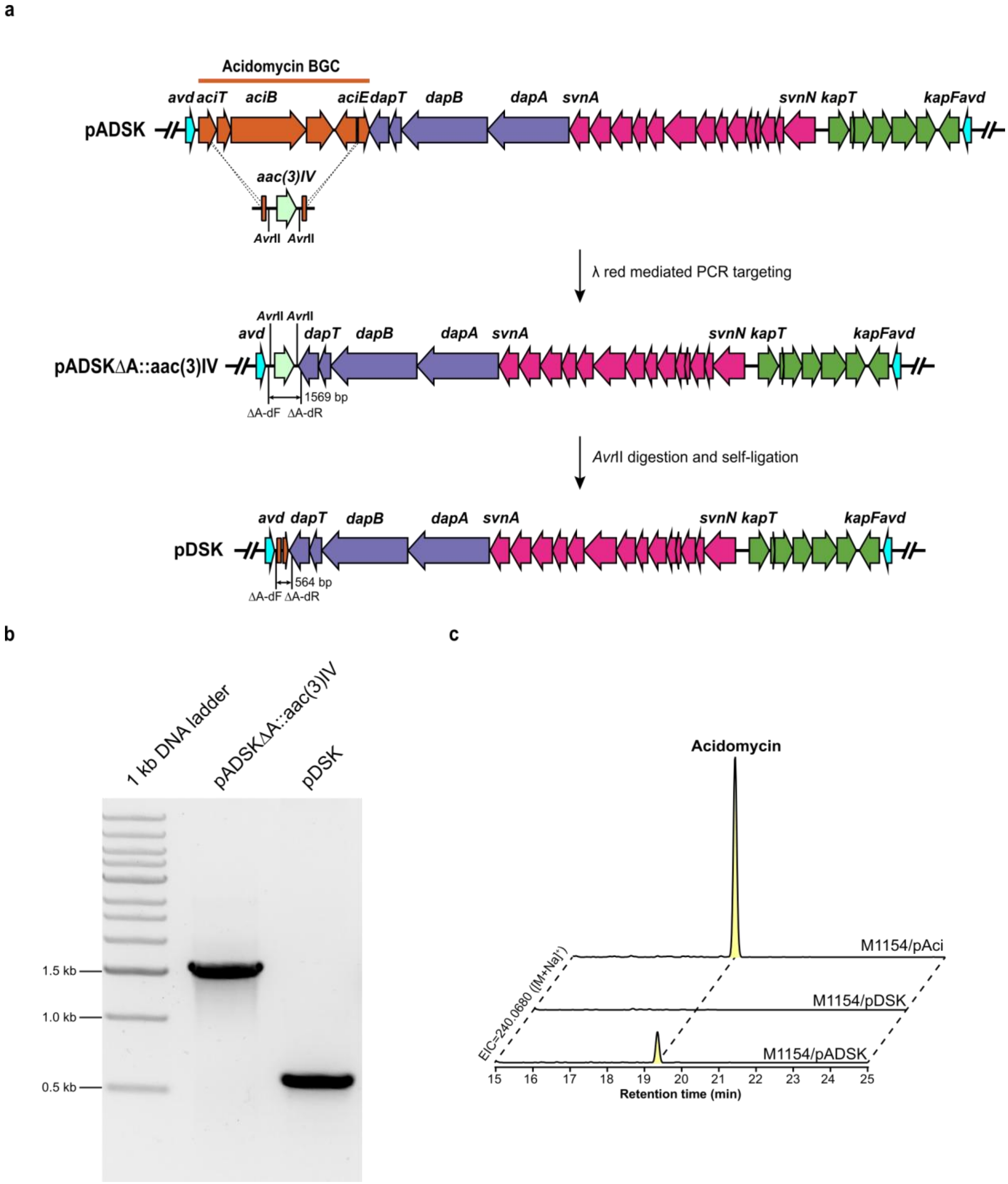


**Figure S7. Target deletion of *aci* BGC from pADSK. a**, Schematic representation of deleting *aci* BGC from pADSK using λ-red PCR targeting. **b**, Gel electrophoresis of PCR products amplified from the mutant plasmids. **c**, Extracted ion chromatogram of acidomycin ($[M+Na]^+$=240.0680) from the fermentation broth of *S. coelicolor* M1154 carrying acidomycin expression plasmids. *aci* BGC deletion abolished the production of acidomycin in the heterologous expression strain.

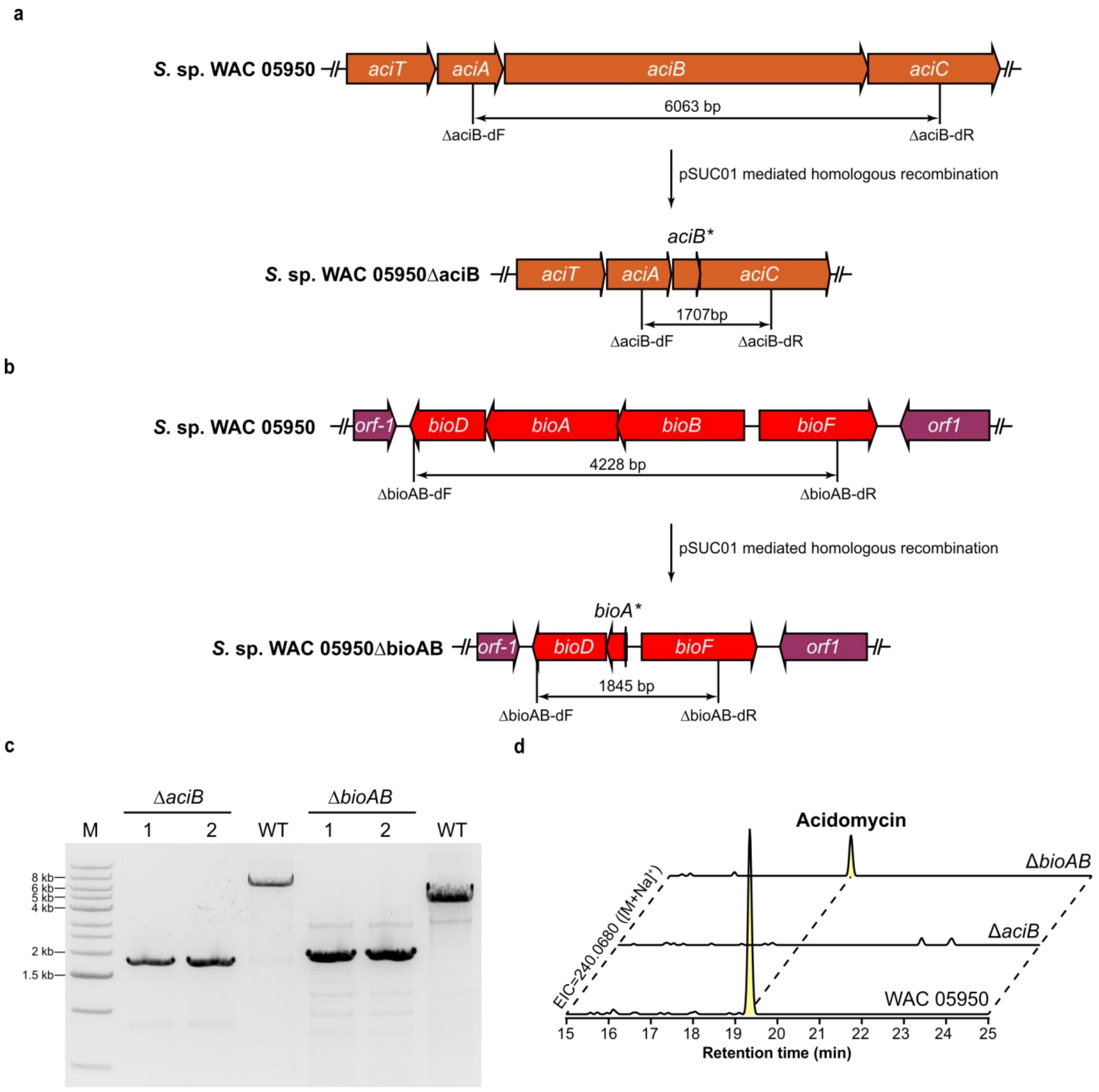


**Figure S8. Target deletion of *aciB* and *bioAB* from the chromosome of WAC05950.** Schematic representations of *aciB* (A) and *bioAB* (B) gene deletions using pSUC01 plasmid-mediated homologous recombination. C. Gel electrophoresis of PC products amplified from the genomic DNA of WAC05950 and Δ*aciB* and Δ*bioAB* mutant strains. D. Extracted ion chromatogram of acidomycin ($[M+Na]^+$=240.0680) from the fermentation broth of WAC05950 and Δ*aciB* and Δ*bioAB* mutant strains. *aciB* deletion abolished the production of acidomycin in WAC05950Δ*aciB*, while *bioAB* deletion could still produce acidomycin in WAC05950Δ*bioAB*.

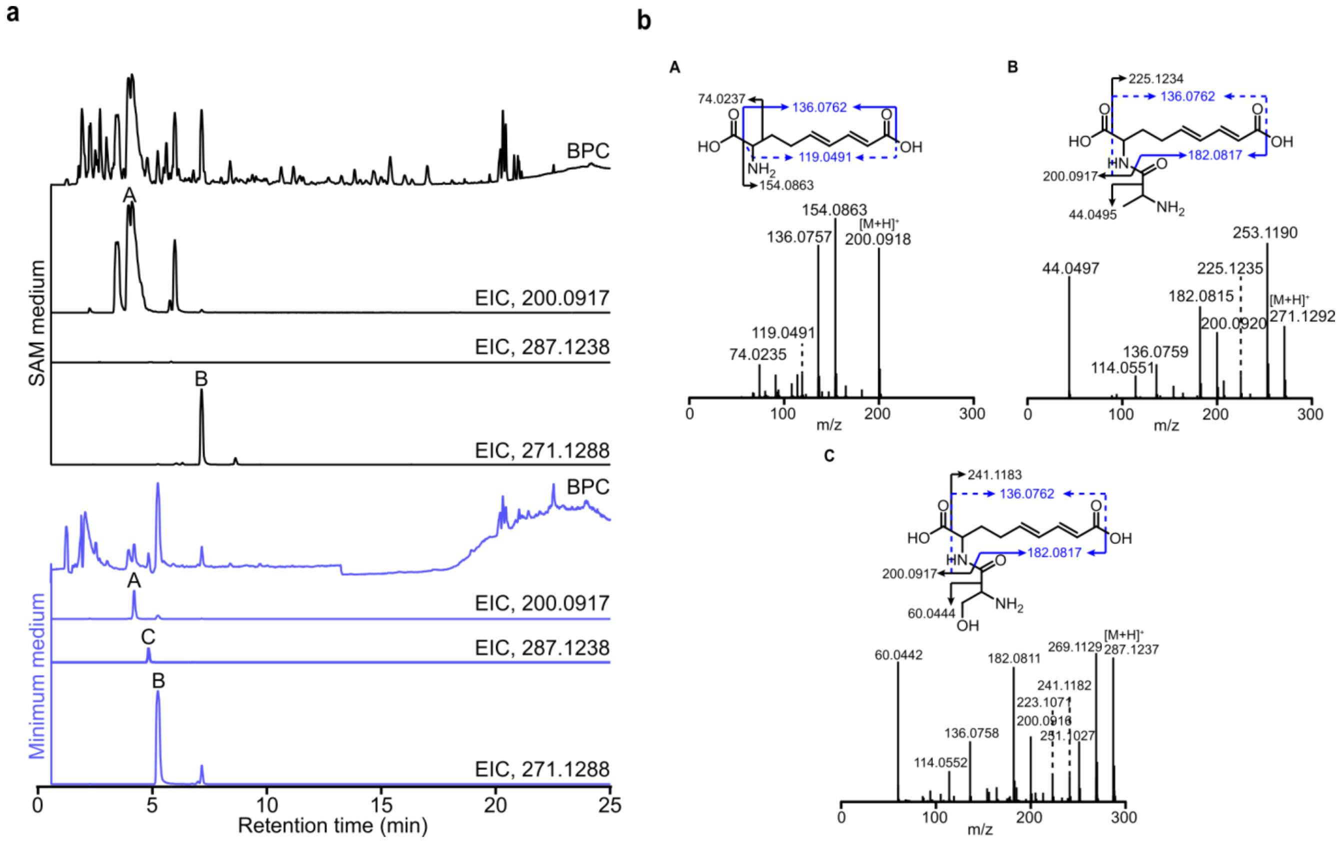


**Figure S9. MS/MS determination of dapamycin A/B/C. a,** EICs of dapamycin A/B/C produced by *S. coelicolor* M1154/pADSK grown in SAM and SMM media. Dapamycin C was only produced when grown in SMM medium. **b,** MS/MS fragmentation of dapamycin A/B/C (CID = 10 eV). The black, blue, and blue dashed arrows indicate daughter ions produced by single, double, and triple bond breakages in the compounds.

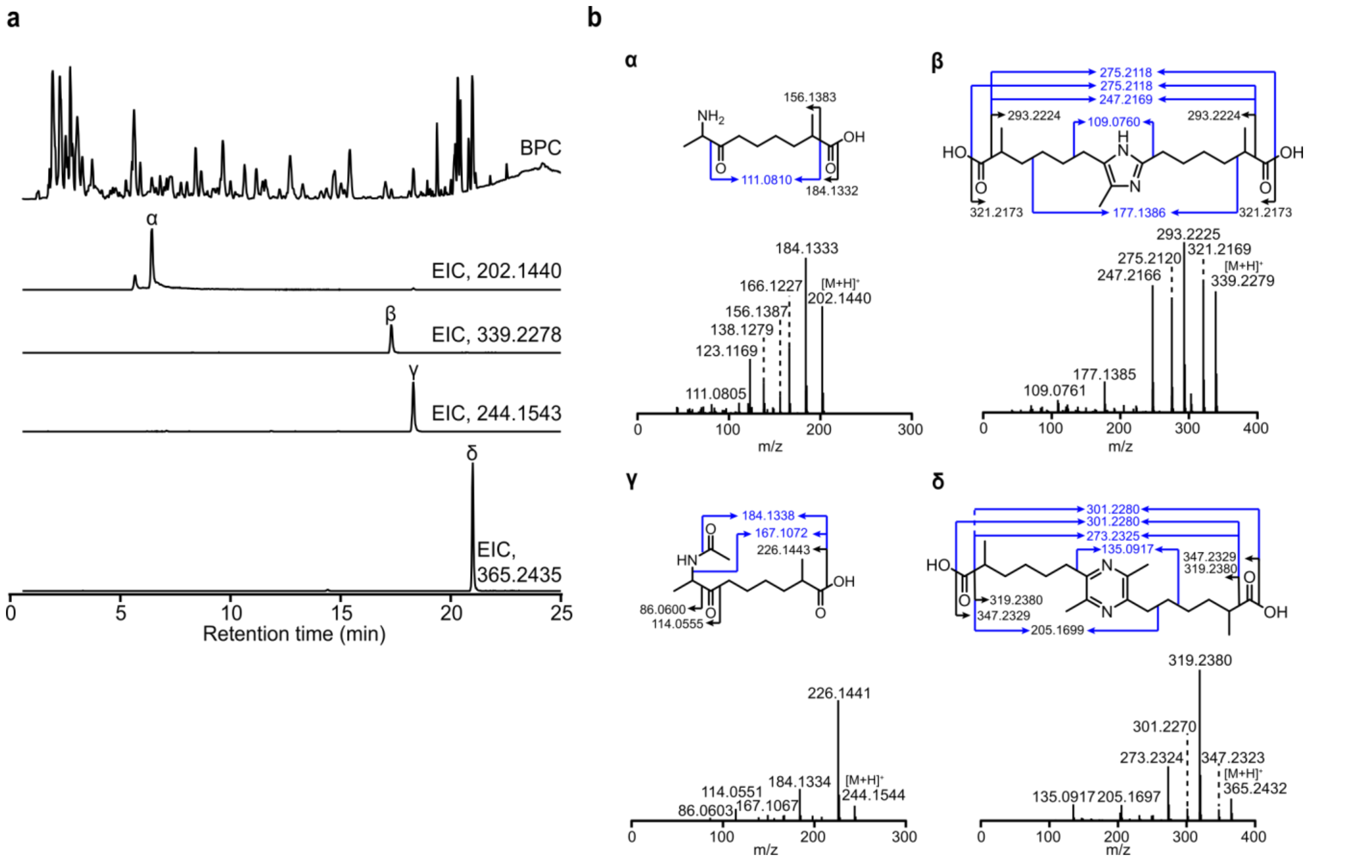


**Figure S10. MS/MS determination of α-Me-KAPA and its analogs. a,** EICs of α-Me-KAPA and its analogs produced by *S. coelicolor* M1154/pADSK. **b,** MS/MS fragmentation of α-Me-KAPA (α, CID = 10 eV), 2,5-di-(2-methylhexanoic acyl)-3-methylimidazole (β, CID = 30 eV), *N*-acetyl- α-Me-KAPA (γ, CID = 5eV), and 2,5-dimethyl-3,6-di-(2-methylhexanoic acyl)pyrazine (δ, CID = 30 eV). The black and blue indicate daughter ions produced by single and double bond breakages in the compounds.

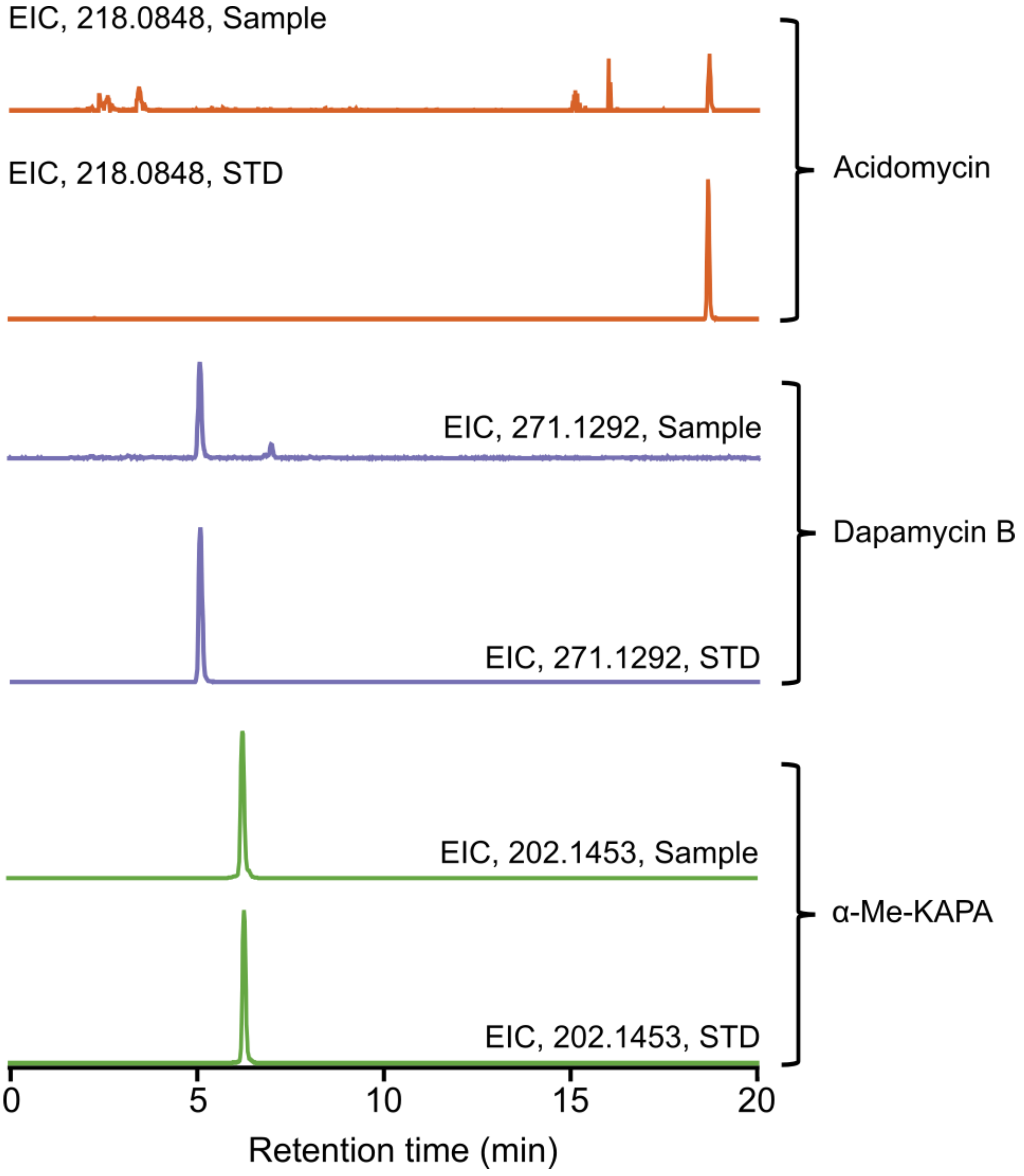


**Figure S11. Co-production of megacluster-derived NPs in the wild-type producer.** Acidomycin (EIC = 218.0848, orange), dapamycin B (EIC = 271.1292, purple) and α-Me-KAPA (EIC = 202.1453, green) are co-produced in the wild-type megacluster-containing strain *Streptomyces* sp. WAC05950 grown in SAM medium over 7 days. Each EIC was extracted from the conditioned media. Purified NPs from the heterologous expression strain acted as standards (STDs).

**a**

| Strain | MIC (μg/mL) | | | | | | | |
|---|---|---|---|---|---|---|---|---|
| | Stravidin S2 | *O*-Me-S2 | Stravidin S4 | Stravidin S5 | Acidomycin | Dapamycin A | Dapamycin B | α-Me-KAPA |
| *E. coli* BW25113 | 0.008 | 0.16 | >128 | 50 | 16 | >128 | 8* | 32 |
| *E. coli* C0244 | 0.004 | 0.16 | 100 | 25 | 32 | >128 | 128* | 32 |
| *K. pneumoniae* MKP103 | 0.064 | 1.56 | >128 | 128 | >128 | >128 | >128 | >128 |
| *K. pneumoniae* ATCC 43816 | 0.064 | 3.13 | >128 | >128 | 32 | >128 | >128 | 8 |
| *A. baumannii* ATCC 17978 | 16 | 100 | >128 | >128 | 64 | >128 | >128 | 128 |
| *P. aeruginosa* PAO1 | >128 | >128 | >128 | >128 | >128 | >128 | >128 | >128 |
| *M. smegmatis* mc$^2$155 | 0.004 | - | - | - | 2 | >128 | 0.25 | 8 |
| *M. fortuitum* ATCC6891 | >64 | - | - | - | 8 | >128 | >128 | 64 |
| *M. abscessus* ATCC19977 | >64 | - | - | - | >128 | >128 | >128 | 128 |
| *M. tuberculosis* H37Ra | 8 | - | - | - | 2 | 64 | >128 | 8 |
| *M. bovis* BCG Pasteur ATCC35734 | 4 | - | - | - | 1 | 16 | 8 | 8 |

**b**

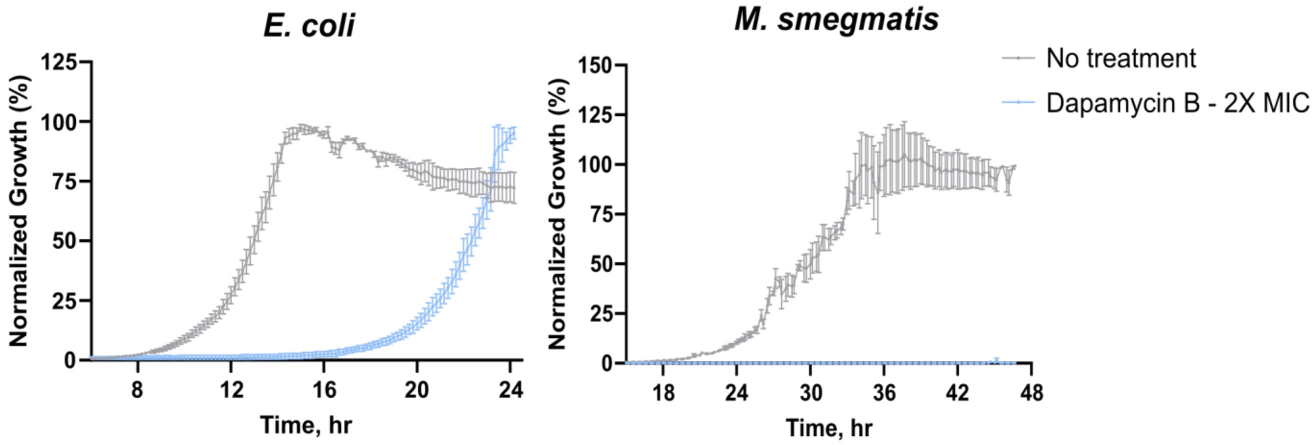


**c**

| Strain | MIC (μg/mL) | |
|---|---|---|
| | Stravidin S2 | Dapamycin B |
| Wild-type | 0.016 | 8 |
| *ΔdppC* | 0.5 | >128 |
| *ΔdtpA* | 0.016 | 8 |
| *ΔoppA* | 0.016 | 8 |

**Figure S12. Antimicrobial activity of the isolated compounds from the megacluster. a,** MIC table of the isolated compounds against Gram-negative bacteria and mycobacteria. An asterisk next to a MIC value denotes transient inhibition, with bacterial growth appearing after an 8-hour growth delay. **b,** Growth kinetics of *E. coli* BW25113 and *M. smegmatis* mc$^2$155 in the absence (grey) and presence (blue) of 2X MIC of dapamycin B (16 µg/mL and 0.5 µg/mL, respectively). Data are presented as mean ± SEM (n = 3). **c,** Comparison of dapamycin B and stravidin S2 activities in wild-type *E. coli* BW25113 and different peptide transporter KEIO mutants.

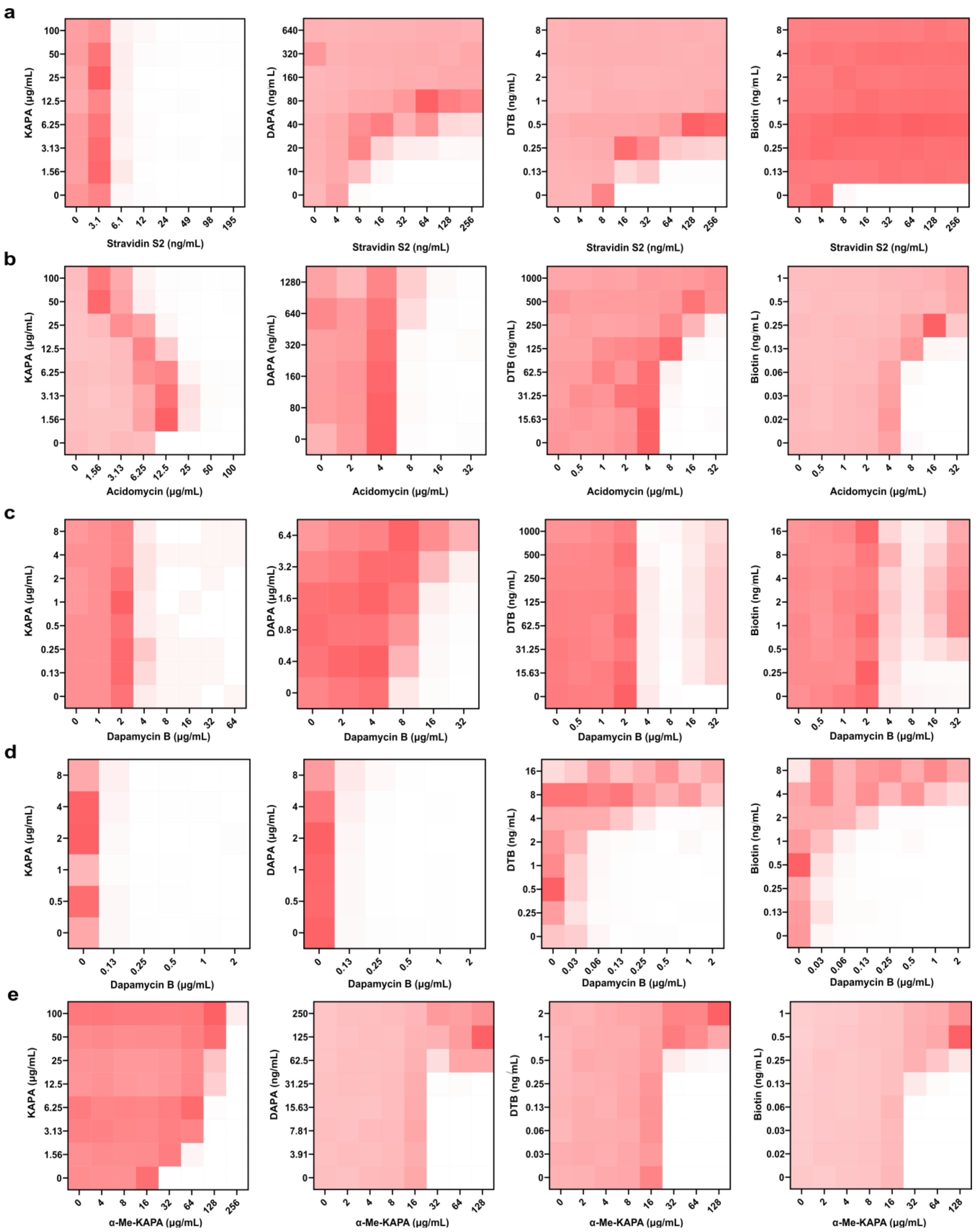


**Figure S13. Activity suppression profiles of stravidin S2, acidomycin, dapamycin B and α-Me-KAPA by biotin and intermediates of its biosynthesis. a-e,** Checkerboard broth microdilution assays of stravidin S2, acidomycin, dapamycin B and α-Me-KAPA against KAPA, DAPA, DTB and biotin in

*E. coli* BW25113 (**a-c, e**) and *M. smegmatis* mc$^2$155 (**d**). Data are representative of at least three biological replicates.

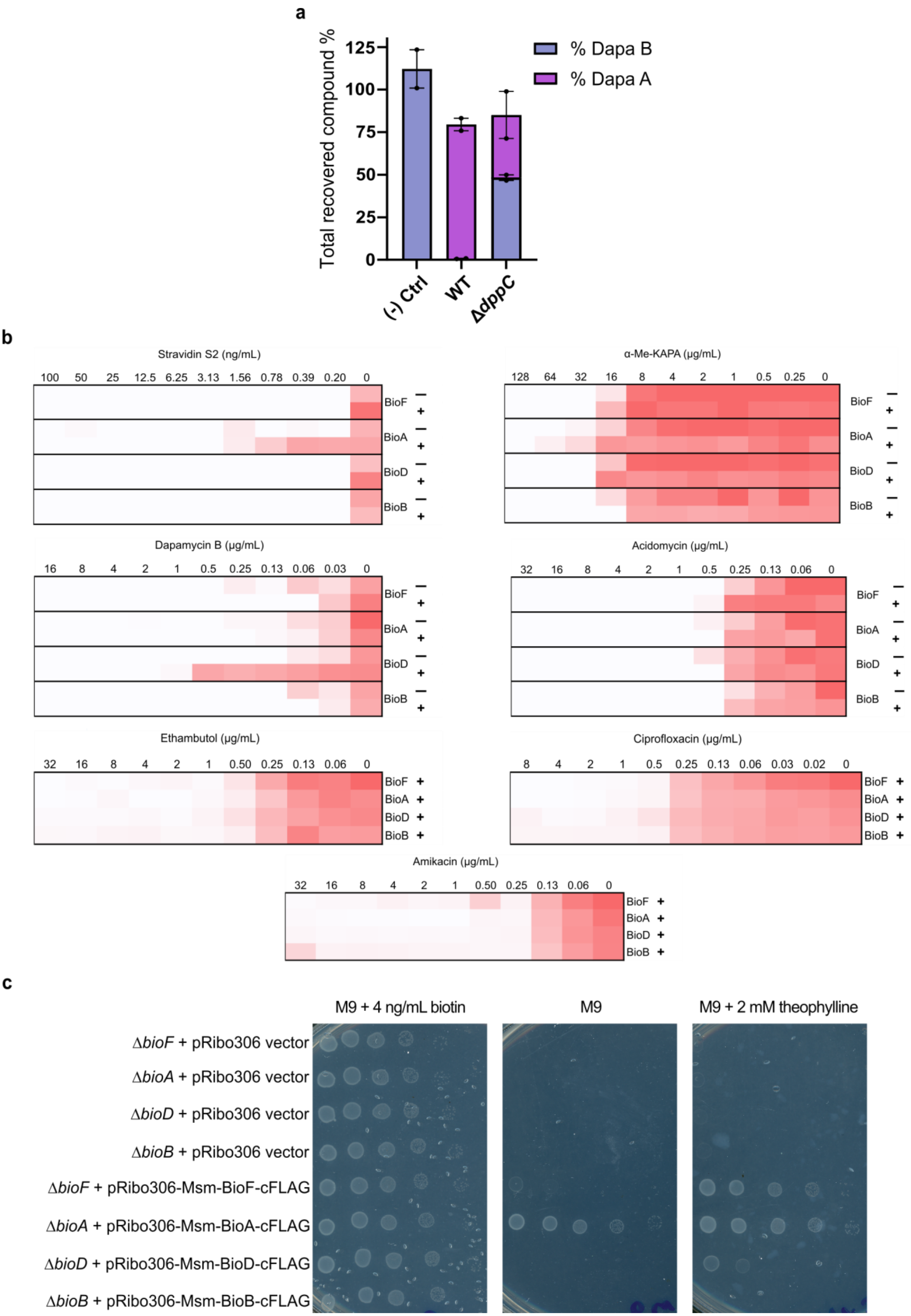
a
Total recovered compound %
% Dapa B
% Dapa A
(-) Ctrl
WT
ΔdppC
b
Stravidin S2 (ng/mL)
α-Me-KAPA (μg/mL)
Dapamycin B (μg/mL)
Acidomycin (μg/mL)
Ethambutol (μg/mL)
Ciprofloxacin (μg/mL)
Amikacin (μg/mL)
BioF
BioA
BioD
BioB
c
M9 + 4 ng/mL biotin
M9
M9 + 2 mM theophylline
ΔbioF + pRibo306 vector
ΔbioA + pRibo306 vector
ΔbioD + pRibo306 vector
ΔbioB + pRibo306 vector
ΔbioF + pRibo306-Msm-BioF-cFLAG
ΔbioA + pRibo306-Msm-BioA-cFLAG
ΔbioD + pRibo306-Msm-BioD-cFLAG
ΔbioB + pRibo306-Msm-BioB-cFLAG

**Figure S14. Cellular uptake, metabolism, and genetic evidence implicate BioD as the target of dapamycins. a,** Amount of dapamycin A and B recovered from the supernatant of *E. coli* wild-type and Δ*dppC* cells after a 24-h incubation with 64 µg/mL of dapamycin B. Dapamycin B, incubated in cell-free media, served as a control for non-cell-driven breakdown. Data are represented as two replicates. **b,** Activity of stravidin S2, dapamycin B, α-Me-KAPA and acidomycin against *M. smegmatis* overexpressing *bioFADB* in the presence or absence of theophylline. "-" and "+" indicate cultures grown without or with 4 mM theophylline, respectively. Ethambutol, ciprofloxacin, and amikacin served as negative controls. Data are shown as mean of two biological replicates. **c**, Theophylline-induced overexpression of *M. smegmatis* mc$^2$155 *bioFADB* in the corresponding auxotrophic *E. coli* BW25113 Δ*bioFADB* strains. M9 minimal agar with and without biotin supplementation served as positive and negative controls for growth of the auxotrophs, respectively.

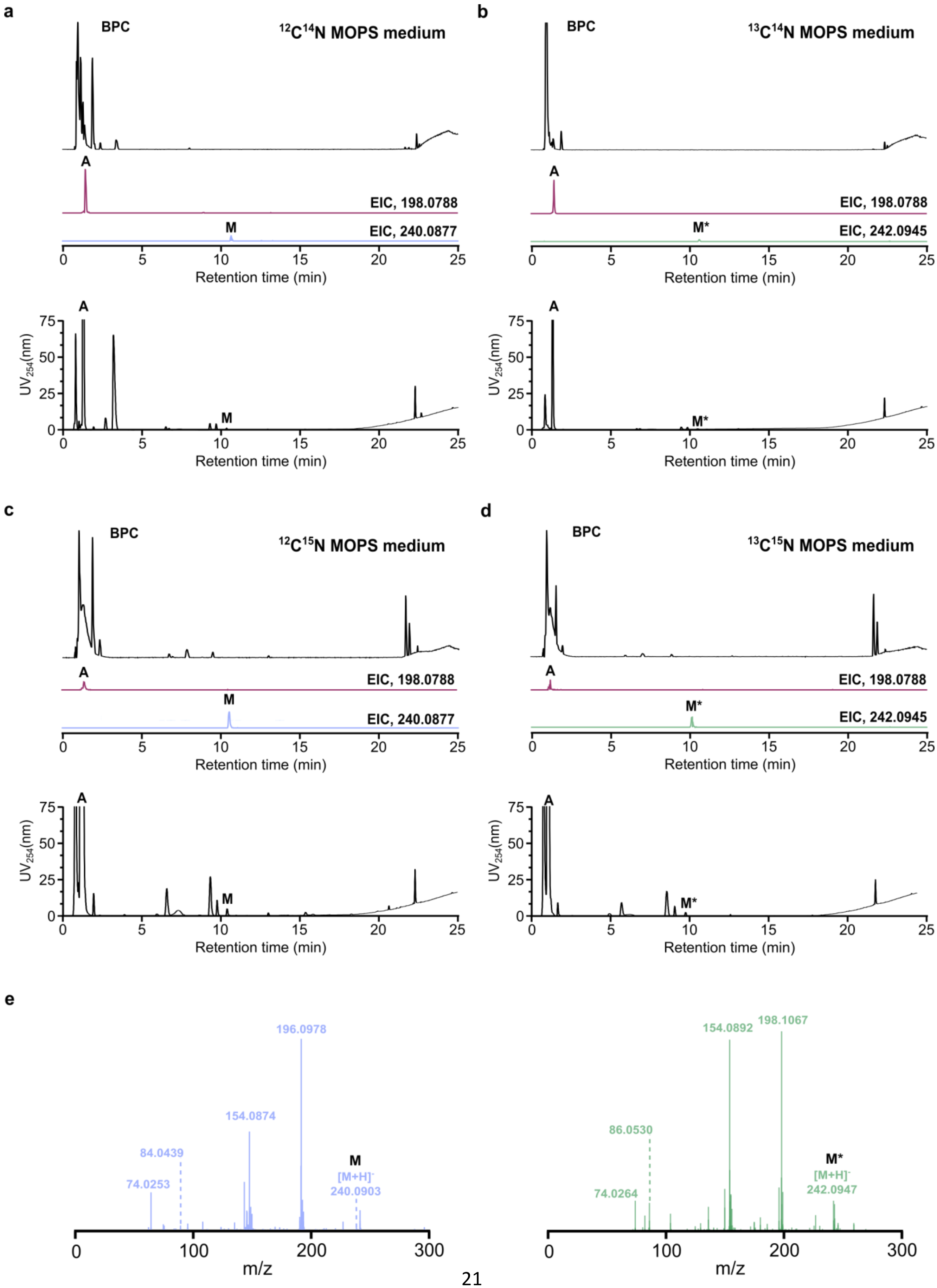
a
BPC
12C14N MOPS medium
A
EIC, 198.0788
M
EIC, 240.0877
Retention time (min)
UV254(nm)
b
BPC
13C14N MOPS medium
A
EIC, 198.0788
M*
EIC, 242.0945
c
BPC
12C15N MOPS medium
EIC, 198.0788
EIC, 240.0877
d
BPC
13C15N MOPS medium
EIC, 198.0788
EIC, 242.0945
e
196.0978
154.0874
84.0439
74.0253
[M+H]-
240.0903
154.0892
198.1067
86.0530
74.0264
242.0947
m/z

**Figure S15. Formation of a dapamycin A-derived metabolite in *E. coli*. a-d,** EICs and $UV_{254}$ of dapamycin A (A, magenta) and the metabolite (M and M*, lavender and green, respectively) produced in *E. coli* after incubation with dapamycin B in $^{12}C^{14}N$ (**a**), $^{13}C^{14}N$ (**b**), $^{12}C^{15}N$ (**c**) and $^{13}C^{15}N$ (**d**) MOPS minimal media. **e**, MS/MS fragmentation of M in $^{12}C^{14}N$ and $^{12}C^{15}N$ media (M, CID = 10 eV), and M* in $^{13}C^{14}N$ and $^{13}C^{15}N$ media (M*, CID = 10 eV).

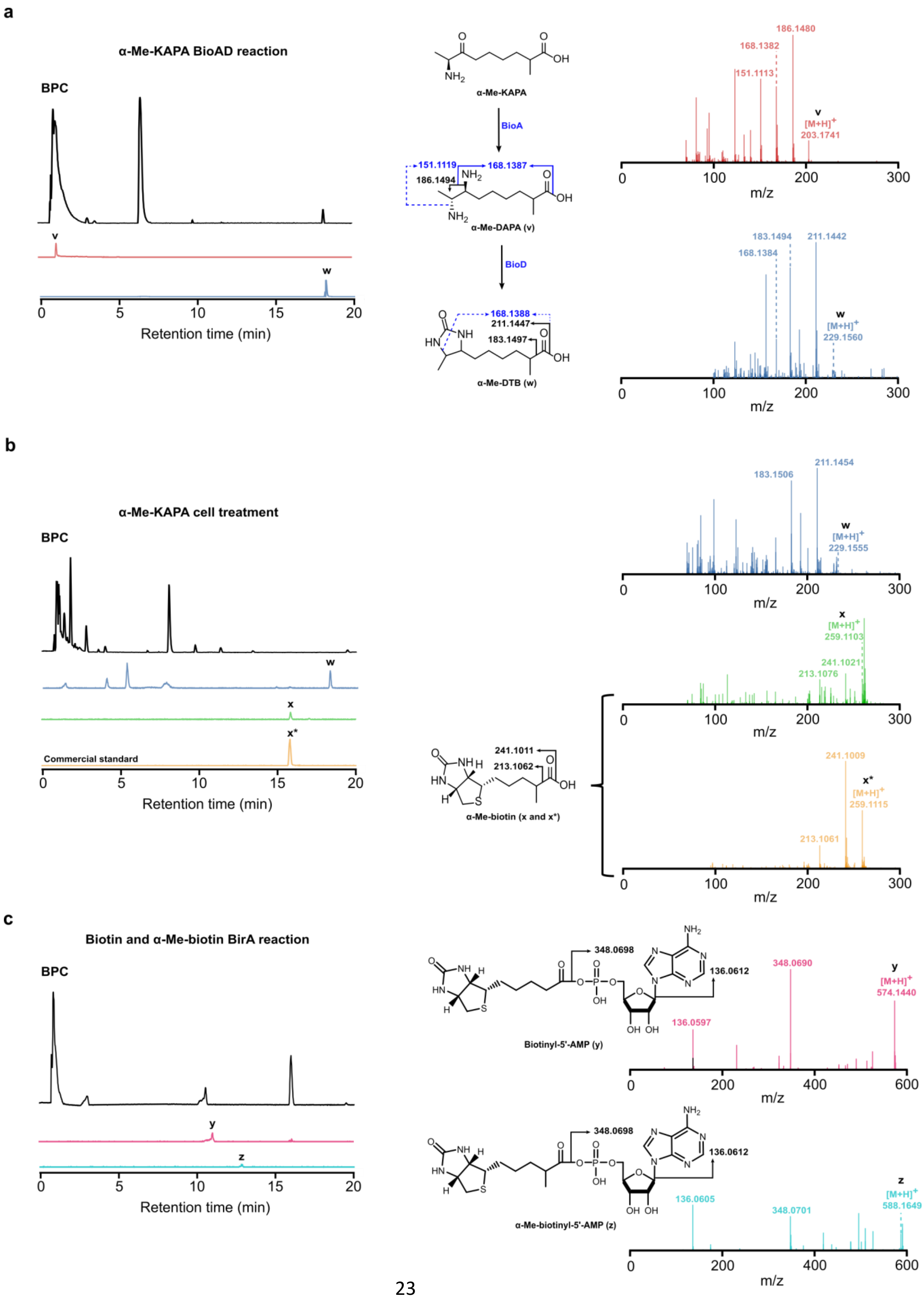
a
α-Me-KAPA BioAD reaction
BPC
v
w
Retention time (min)
α-Me-KAPA
BioA
151.1119
168.1387
186.1494
α-Me-DAPA (v)
BioD
168.1388
211.1447
183.1497
α-Me-DTB (w)
186.1480
168.1382
151.1113
v
[M+H]+
203.1741
m/z
183.1494
211.1442
168.1384
w
[M+H]+
229.1560
b
α-Me-KAPA cell treatment
Commercial standard
x
x*
211.1454
183.1506
229.1555
[M+H]+
259.1103
241.1021
213.1076
241.1011
213.1062
α-Me-biotin (x and x*)
241.1009
259.1115
213.1061
c
Biotin and α-Me-biotin BirA reaction
y
z
348.0698
136.0612
Biotinyl-5'-AMP (y)
348.0690
136.0597
574.1440
α-Me-biotinyl-5'-AMP (z)
136.0605
348.0701
588.1649

**Figure S16. α-Me-KAPA hijacks biotin biosynthesis. a,** EIC and MS/MS fragmentation of α-Me-DAPA (v, red, CID = 15 eV) produced by a BioA enzymatic reaction with α-Me-KAPA, and α-Me-DTB (w, blue, CID = 20 eV) produced by a coupled BioA-BioD. **b,** EICs and MS/MS fragmentations of α-Me-DTB (w, blue, CID = 20 eV) and α-Me-biotin (x, green, CID = 15 eV) produced in whole cells after α-Me-KAPA treatment in *E. coli* BW25113 at ¼×MIC. Commercially acquired α-Me-biotin (x*, orange, CID = 10 eV) was used as a control. **c,** EICs and MS/MS fragmentations of biotinyl-5'-AMP (y, pink, CID = 15 eV) and α-Me-biotinyl-5'-AMP (z, cyan, CID = 15 eV) produced by a BirA enzymatic reaction from biotin and α-Me-biotin, respectively.

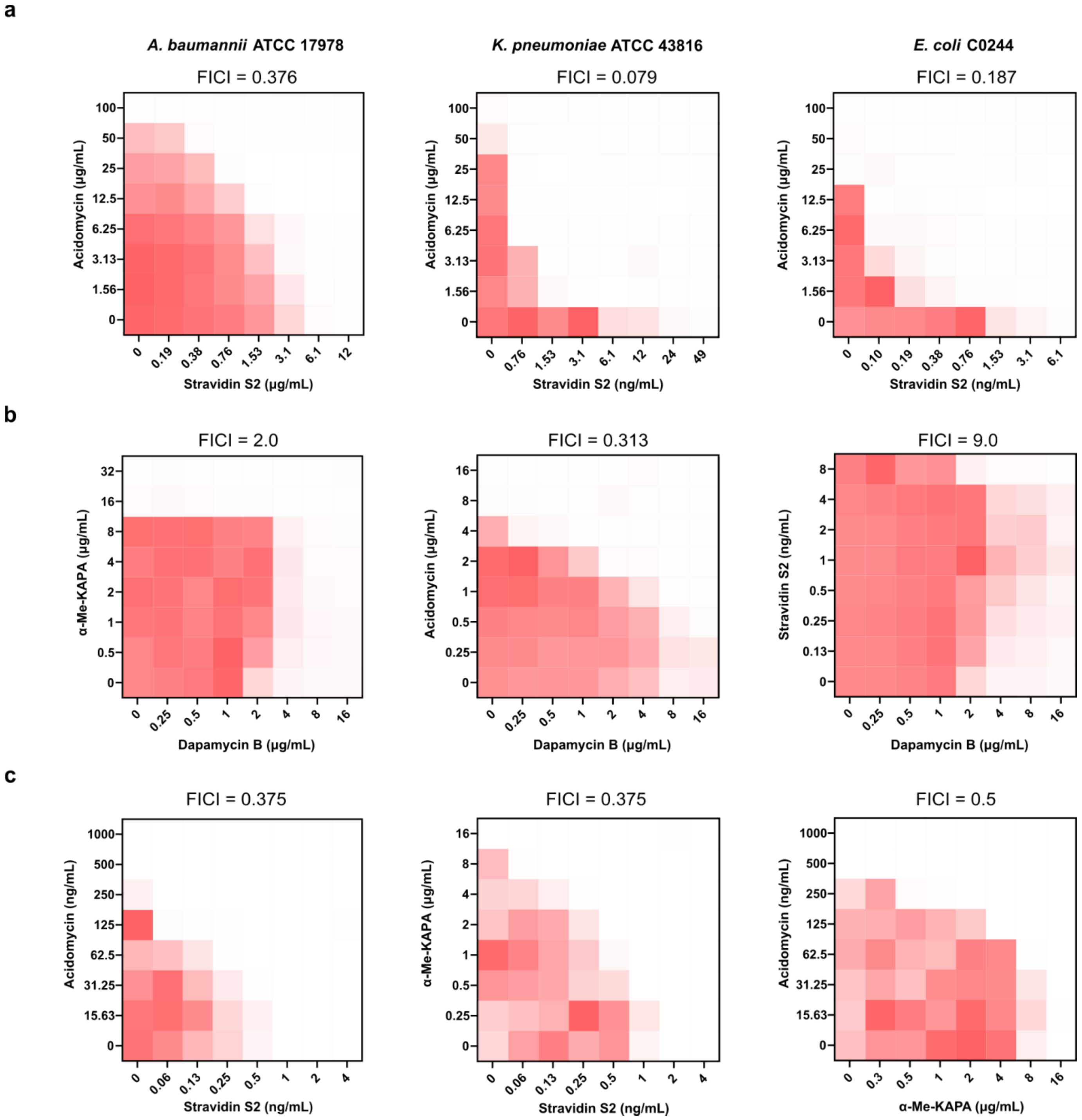


**Figure S17. Pairwise combinations of stravidin S2, acidomycin, dapamycin B and α-Me-KAPA in multiple Gram-negative species and *M. smegmatis*. a,** Checkerboard broth microdilution assays of stravidin S2 against acidomycin in *A. baumannii* ATCC 17978, *K. pneumoniae* ATCC 43816 and *E. coli* C0244. **b,** Checkerboard broth microdilution assays of dapamycin B against acidomycin, α-Me-KAPA and stravidin S2 in *E. coli* BW25113. **c,** Checkerboard broth microdilution assays of pair-wise combinations between stravidin S2, acidomycin and α-Me-KAPA in *M. smegmatis* $mc^2155$. Checkerboard and FICI data are representative of at least three biological replicates.

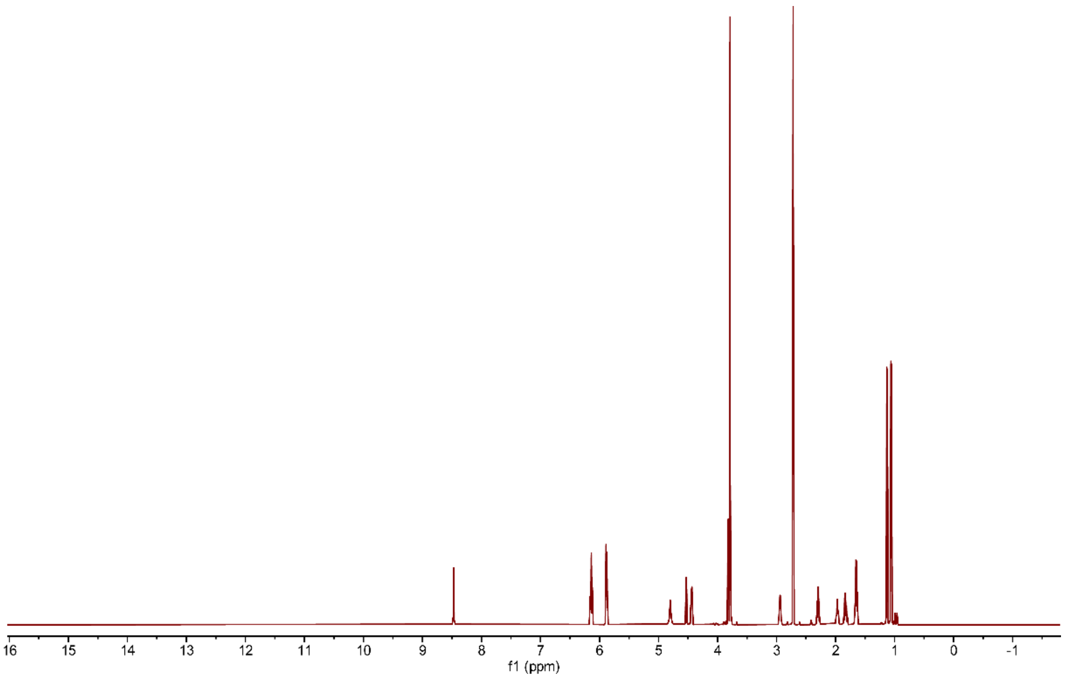


**Figure S18. $^{1}H$ NMR of *O*-Me-stravidin S2 in $D_2O$.**

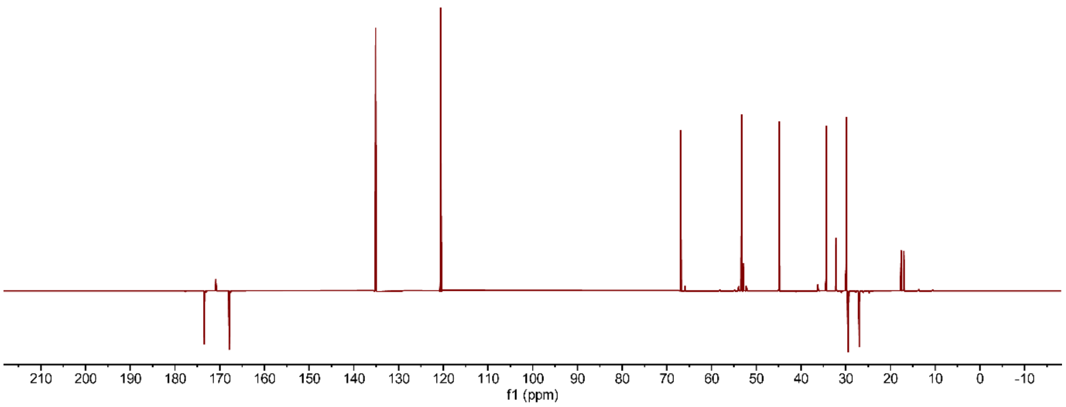


**Figure S19. $^{13}C$ DEPTQ NMR of *O*-Me-stravidin S2 in $D_2O$.**

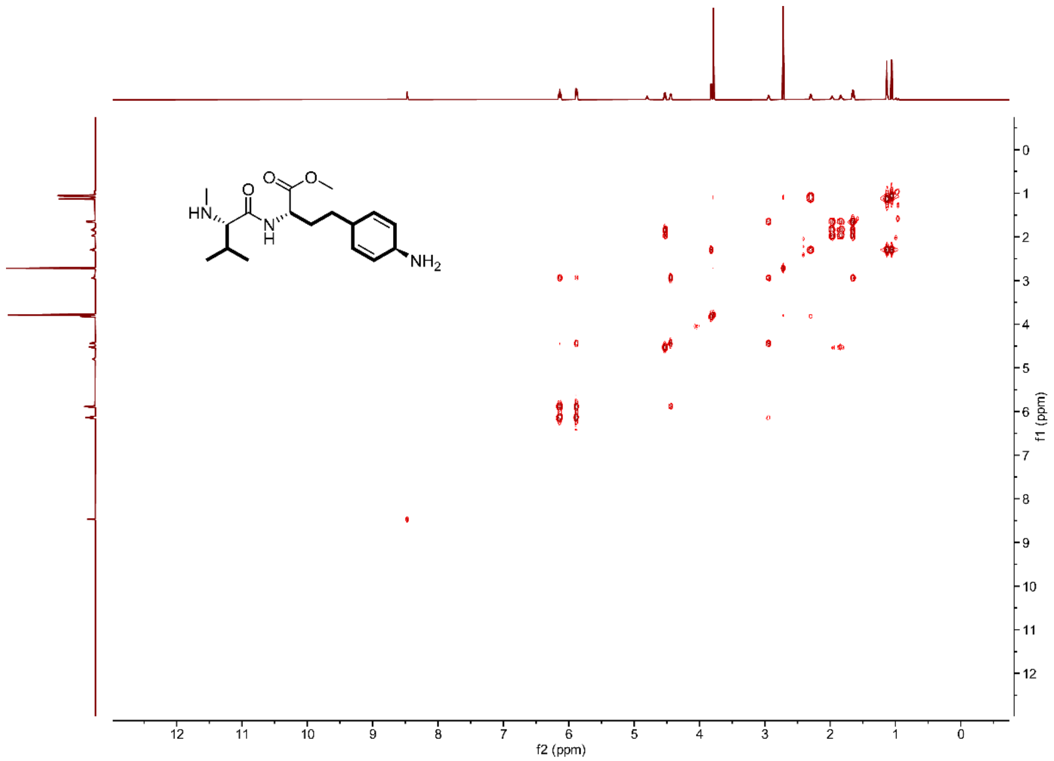

**Figure S20.** **$^1$H-$^1$H COSY NMR of *O*-Me-stravidin S2 in $D_2O$.**

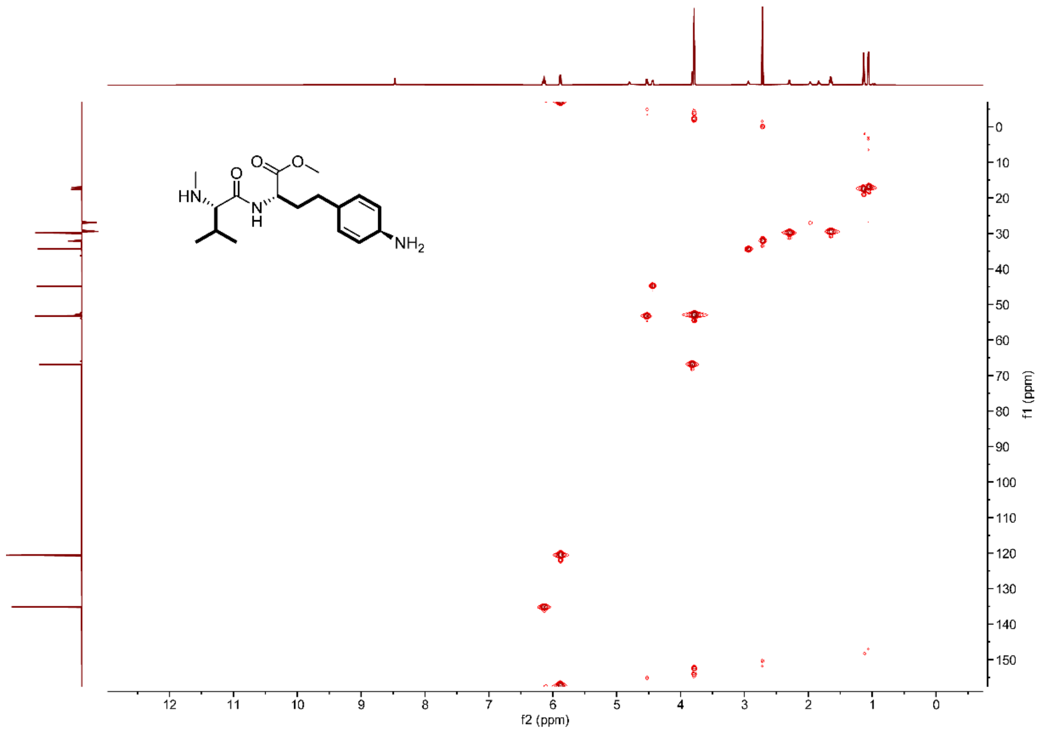

**Figure S21.** **$^1$H-$^{13}$C HSQC NMR of *O*-Me-stravidin S2 in $D_2O$.**

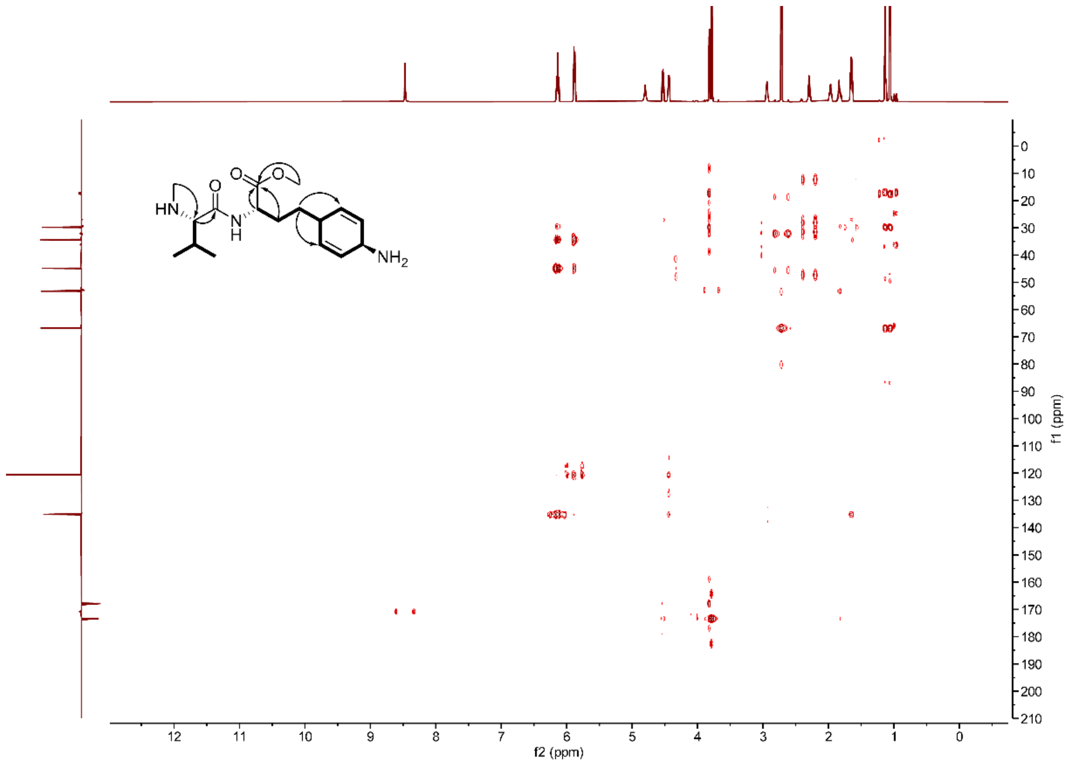


**Figure S22.** **$^{1}H$-$^{13}C$ HMBC NMR of *O*-Me-stravidin S2 in $D_2O$.**

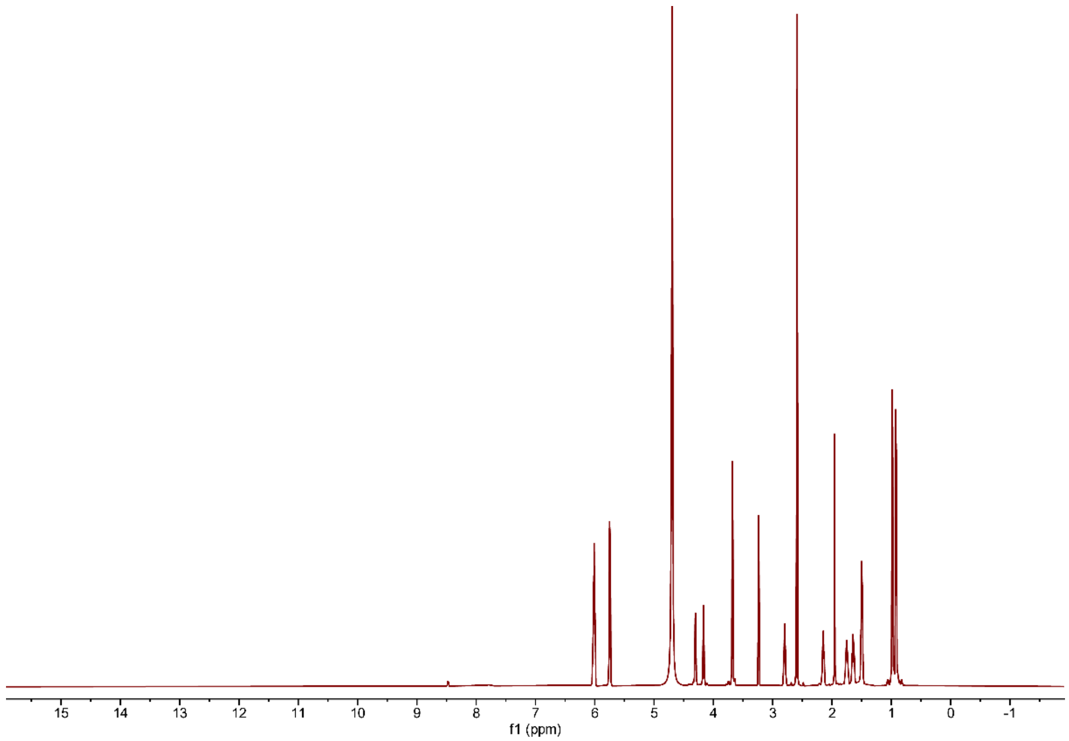


**Figure S23. $^{1}H$ NMR of stravidin S2 in $D_2O$.**

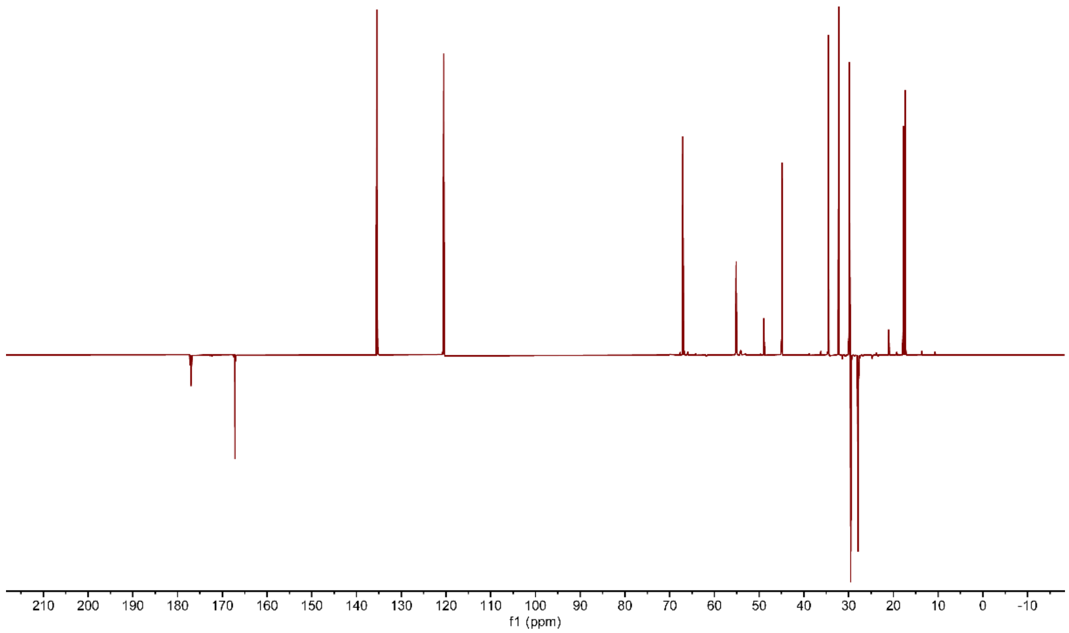


**Figure S24. $^{13}C$ DEPTQ NMR of stravidin S2 in $D_2O$.**

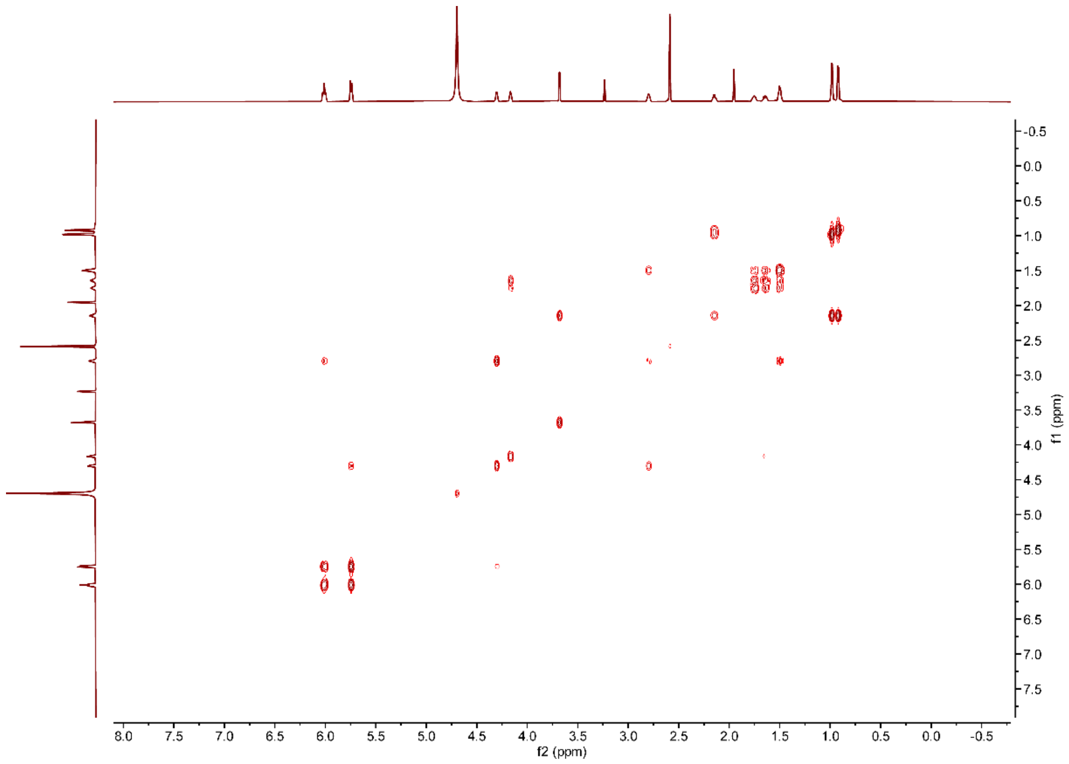


**Figure S25. $^{1}H$-$^{1}H$ COSY NMR of stravidin S2 in $D_2O$.**

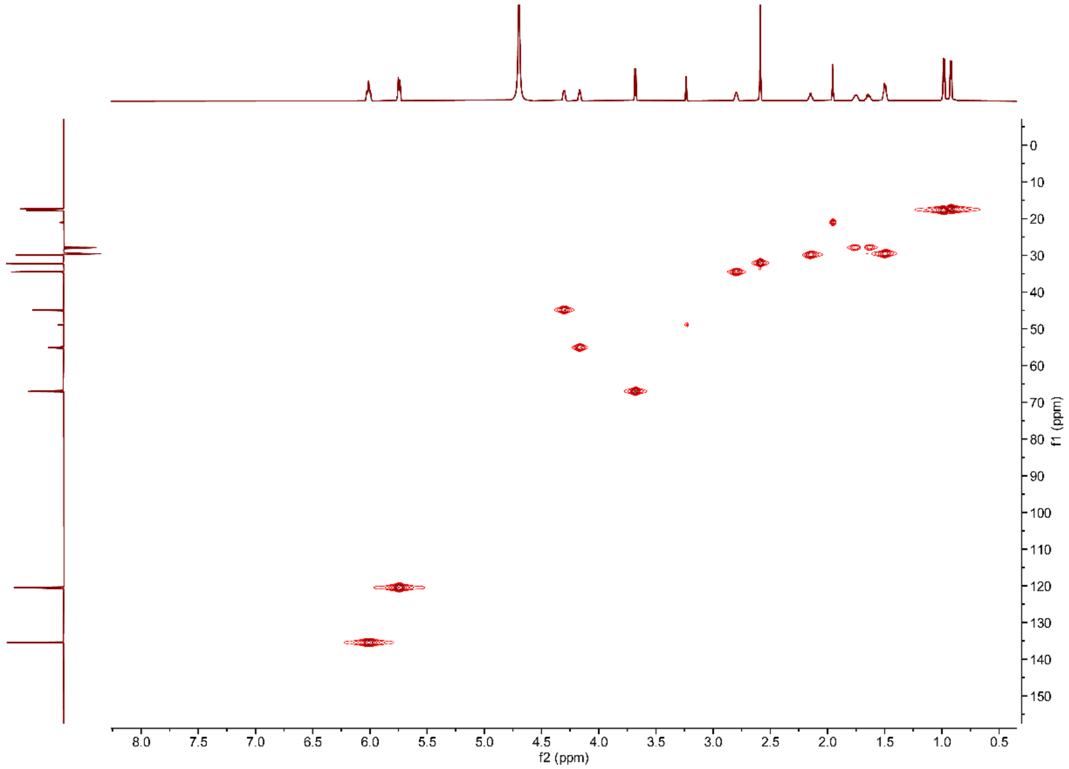


**Figure S26. $^{1}H$-$^{13}C$ HSQC NMR of stravidin S2 in $D_2O$.**

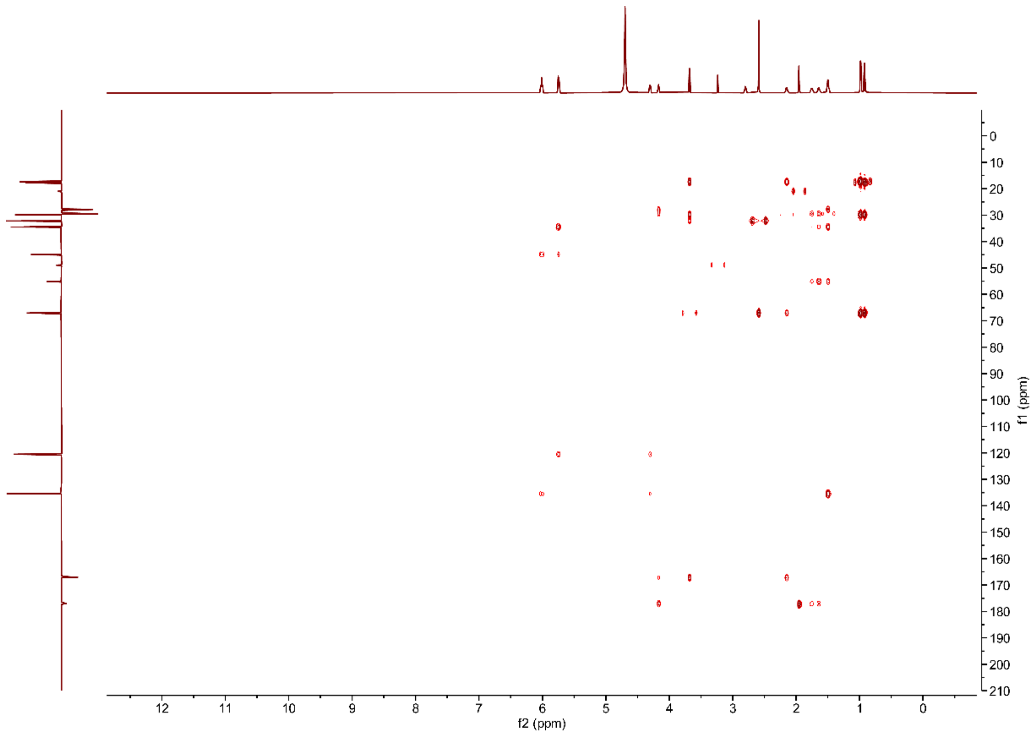


**Figure S27.** $^{1}H$-$^{13}C$ **HMBC NMR of stravidin S2 in $D_2O$.**

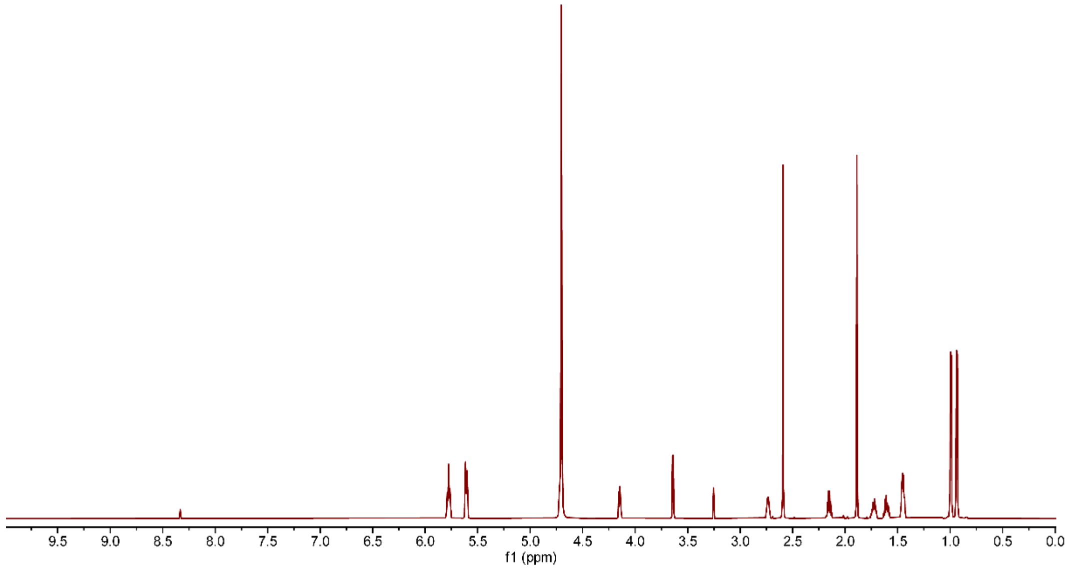


**Figure S28.** $^{1}H$ **NMR of stravidin S4 in $D_2O$.**

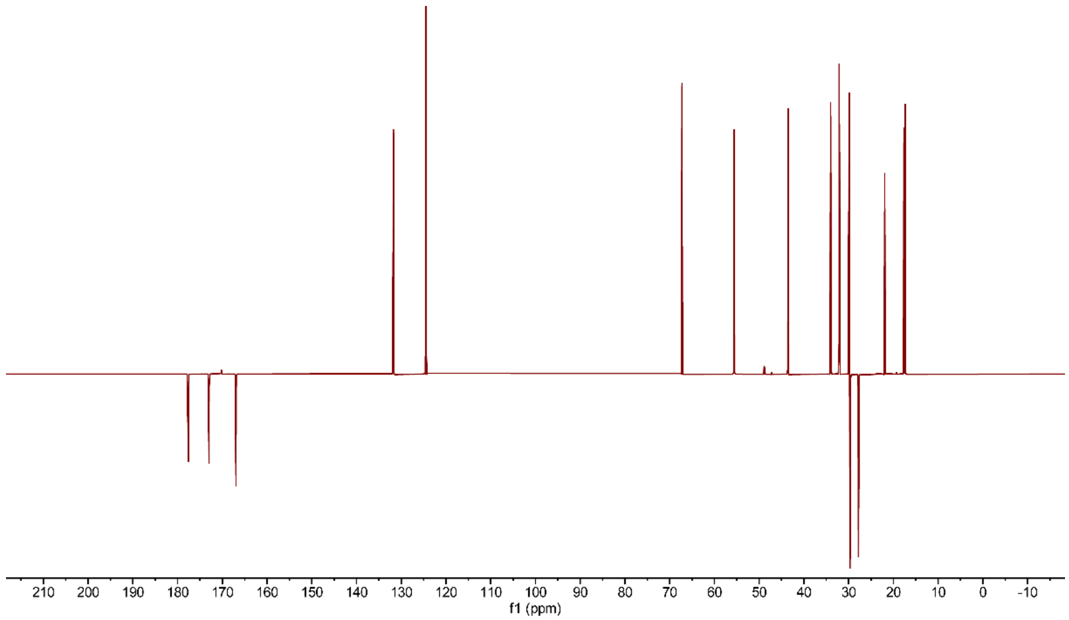


**Figure S29.** $^{13}$**C DEPTQ NMR of stravidin S4 in D$_2$O.**

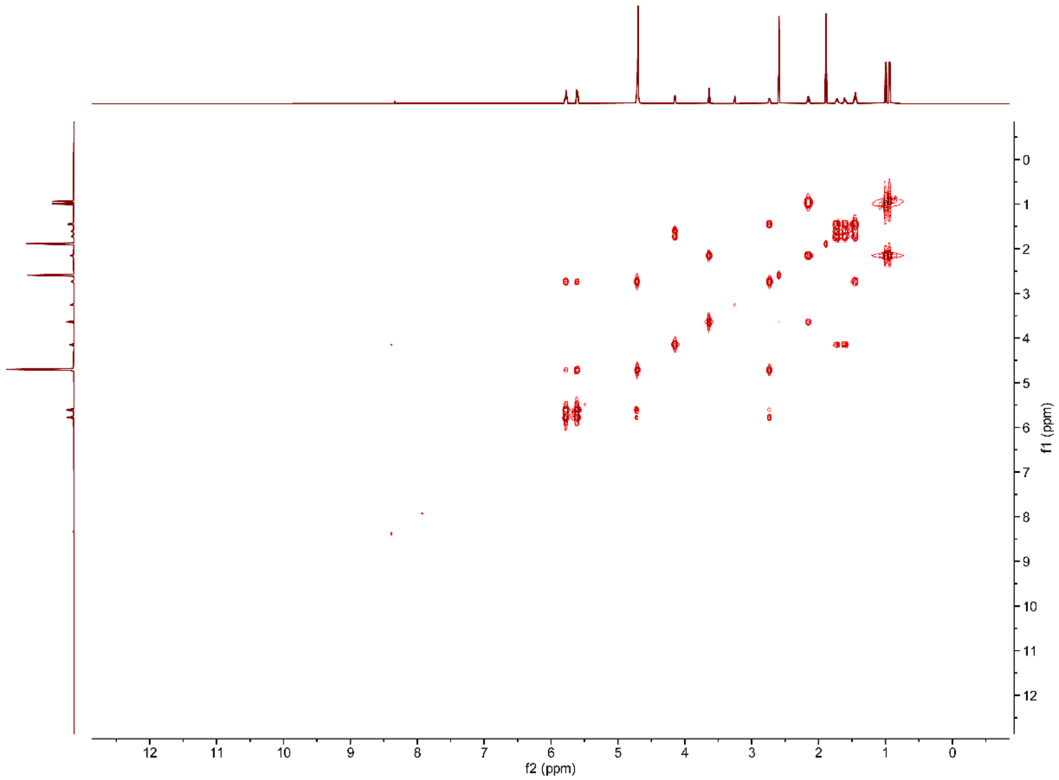


**Figure S30.** $^{1}$**H-**$^{1}$**H COSY NMR of stravidin S4 in D$_2$O.**

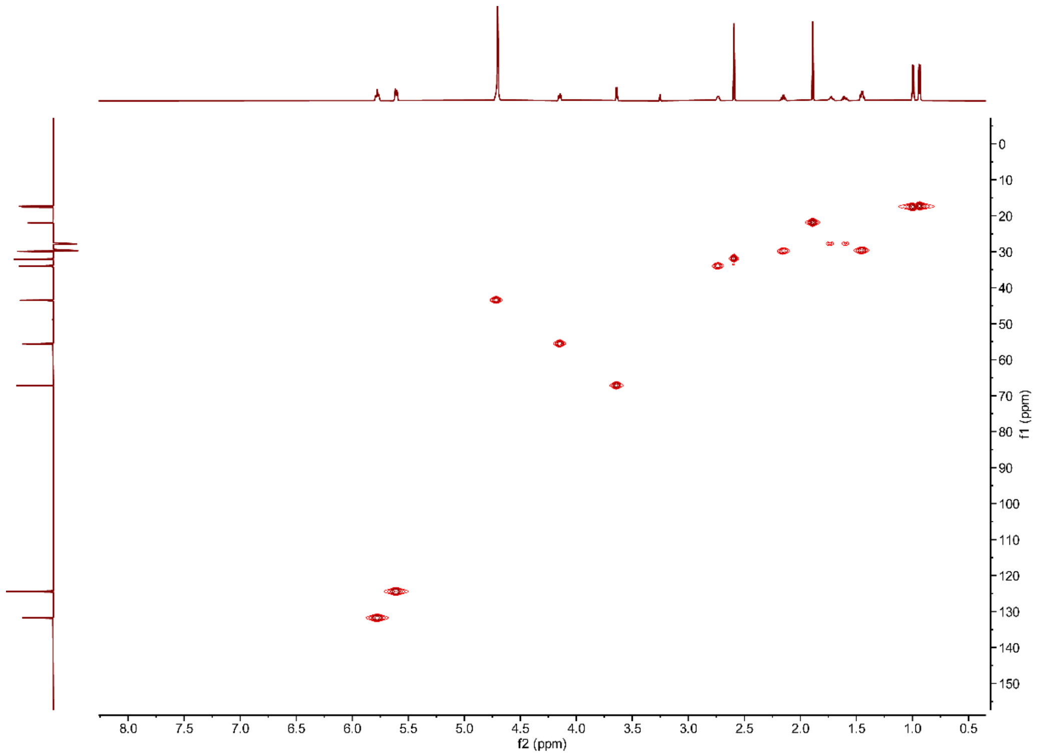


**Figure S31. [1]H-[13]C HSQC NMR of stravidin S4 in $D_2O$.**

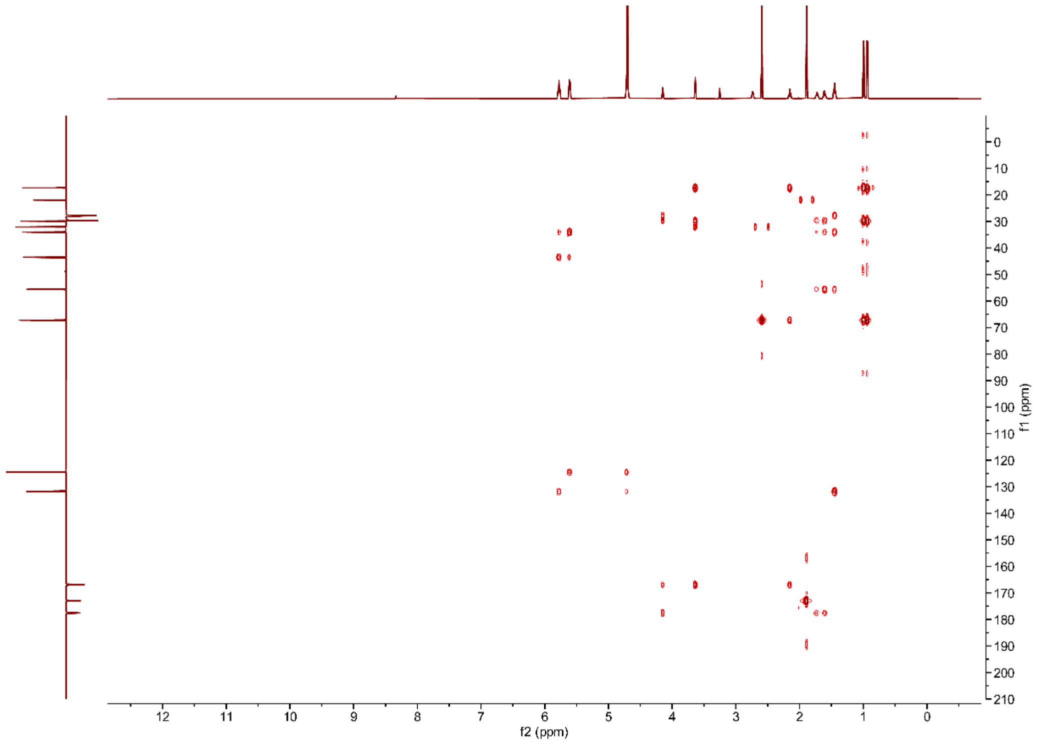


**Figure S32. [1]H-[13]C HMBC NMR of stravidin S4 in DMSO-*d6*.**

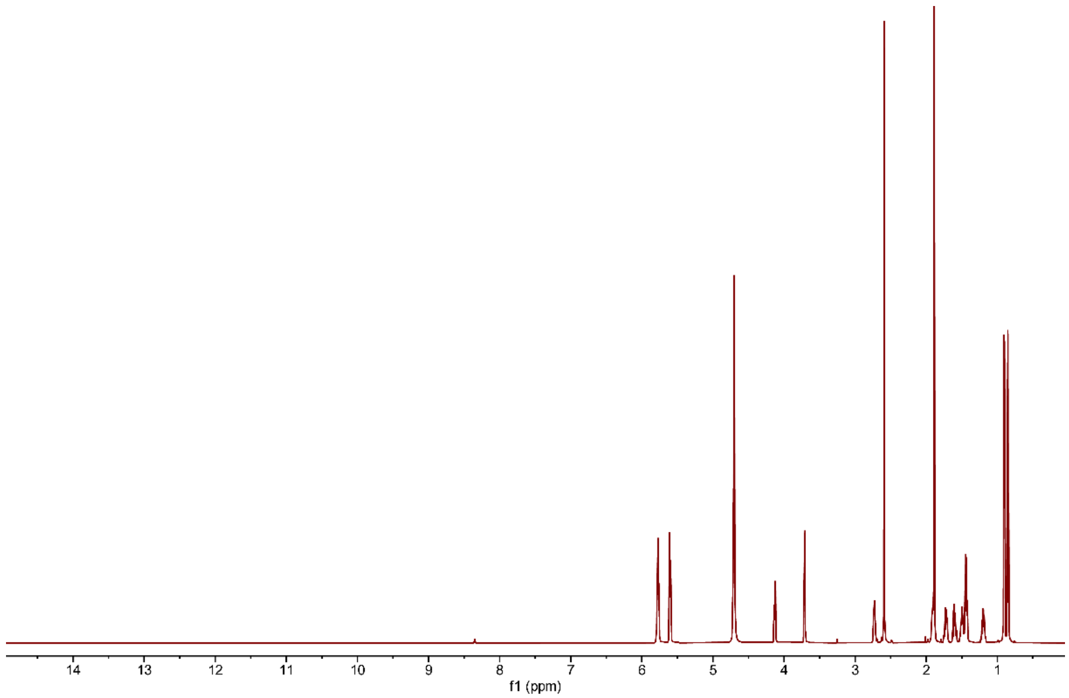


**Figure S33.** **$^1$H NMR of stravidin S5 in $D_2O$.**

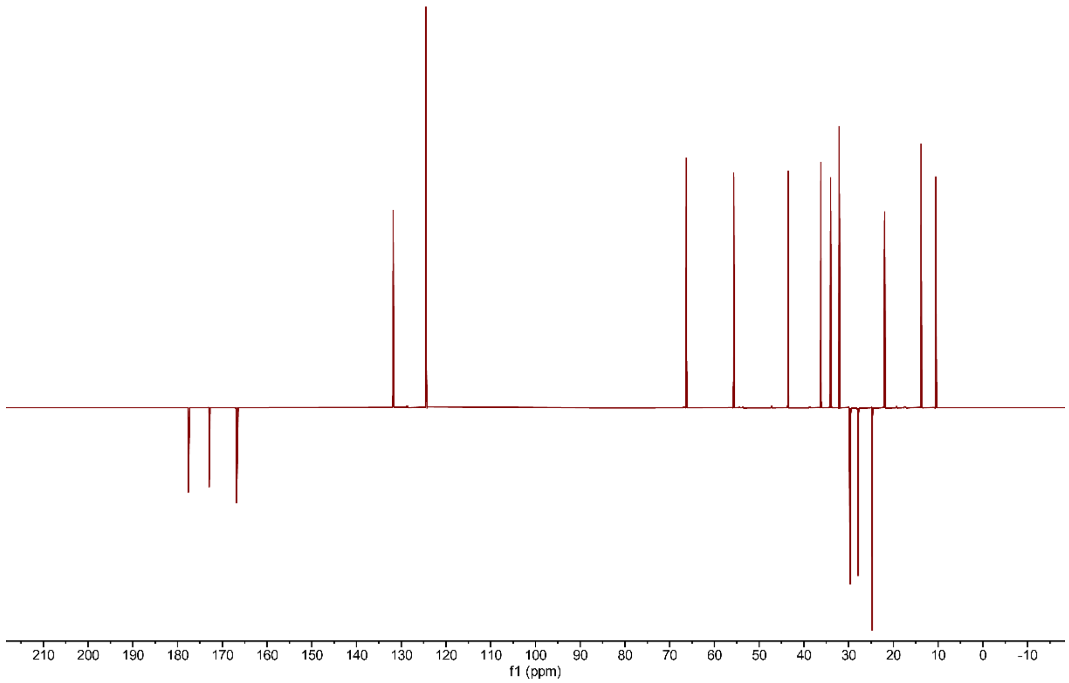


**Figure S34.** **$^{13}$C DEPTQ NMR of stravidin S5 in $D_2O$.**

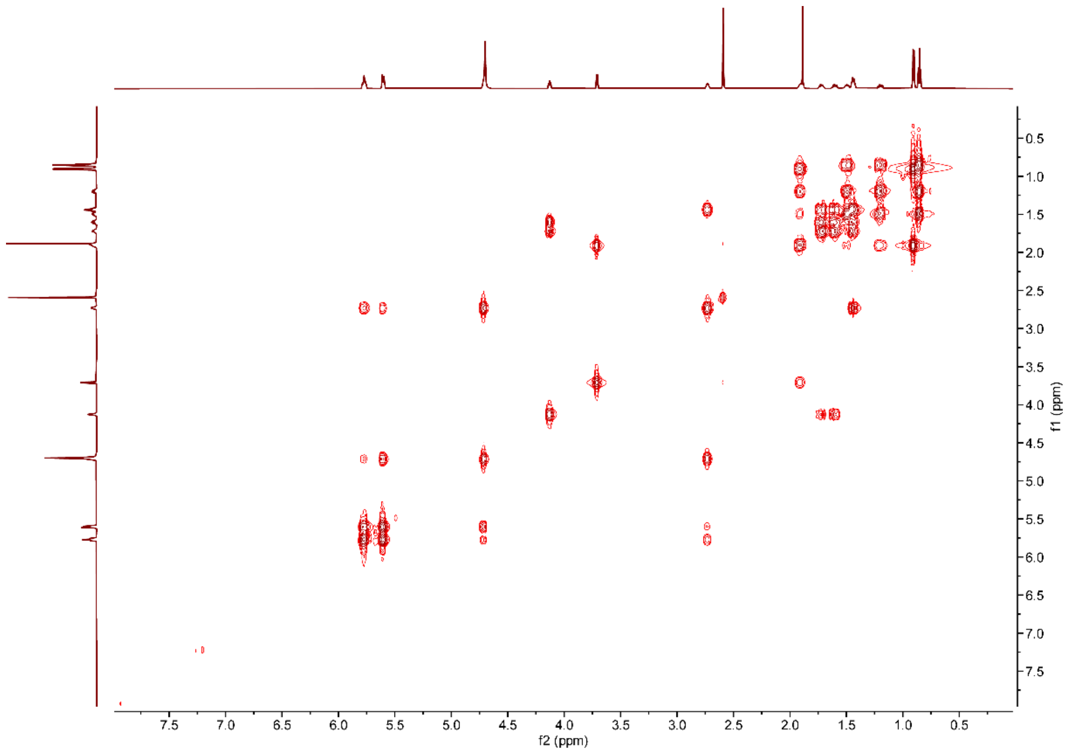


**Figure S35. $^{1}$H-$^{1}$H COSY NMR of stravidin S5 in $D_2O$.**

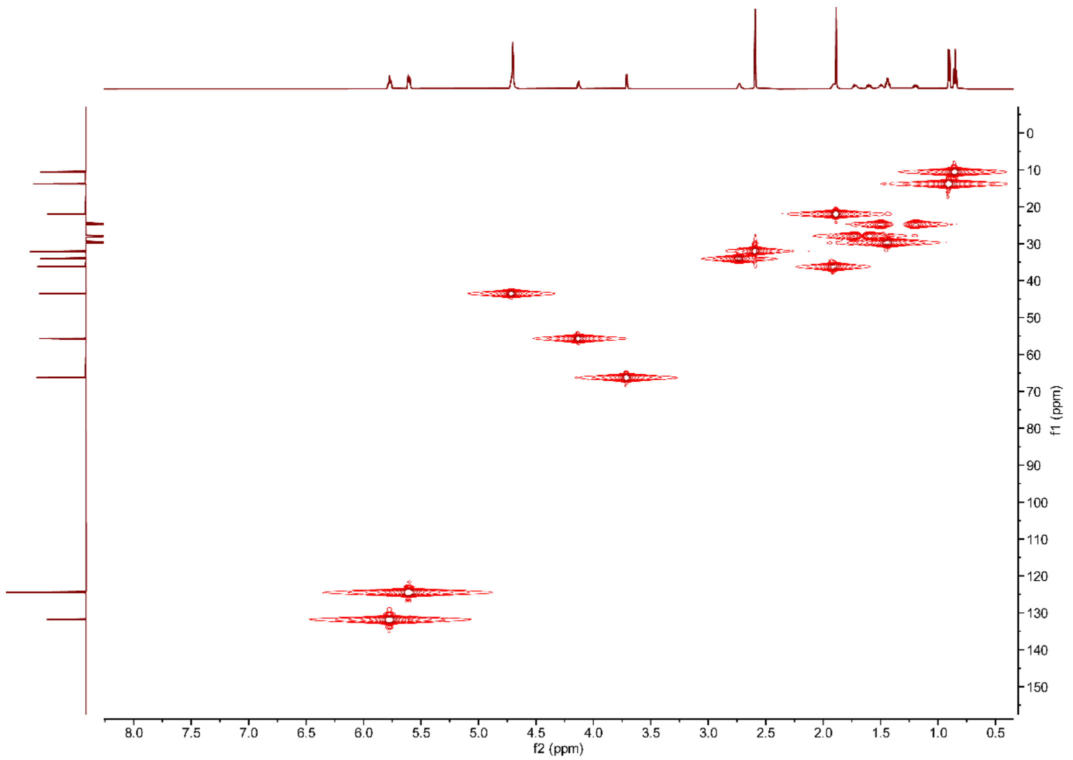


**Figure S36. $^{1}$H-$^{13}$C HSQC NMR of stravidin S5 in $D_2O$.**

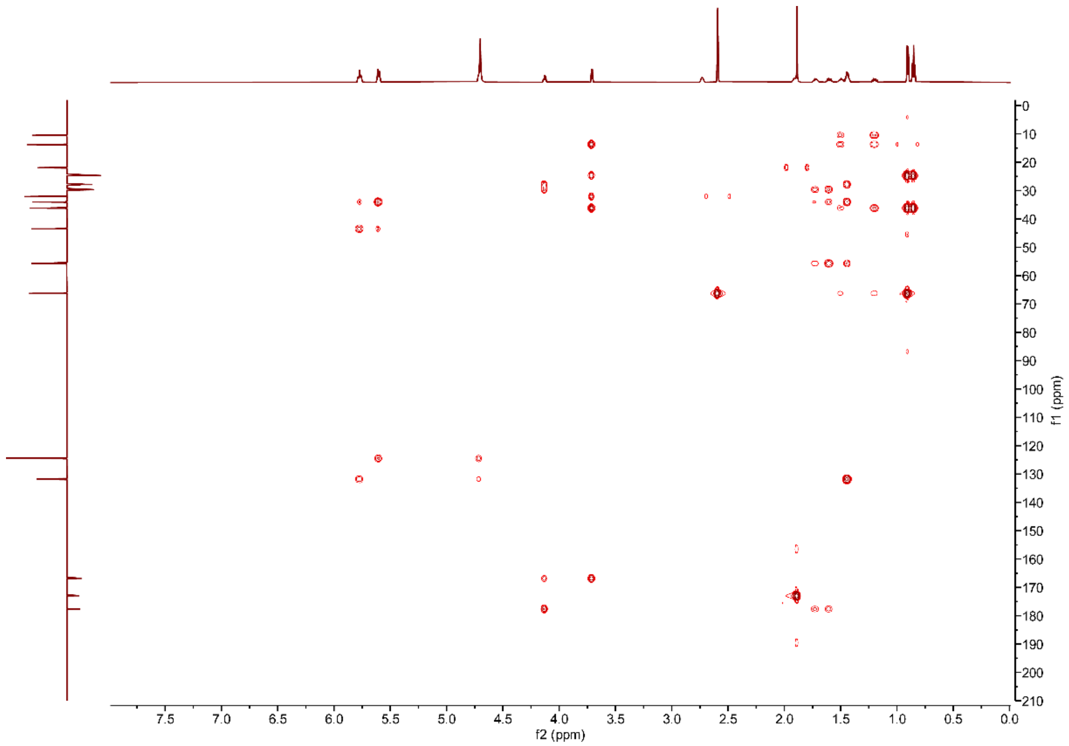


**Figure S37. $^{1}H$-$^{13}C$ HMBC NMR of stravidin S5 in $D_2O$.**

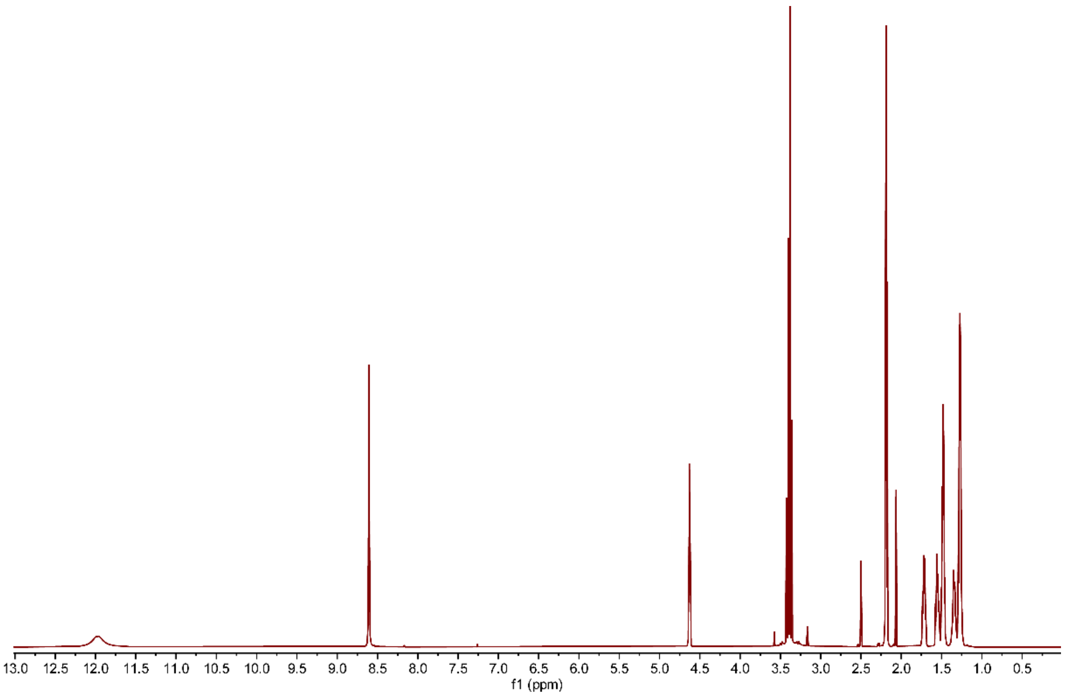


**Figure S38. $^{1}H$ NMR of acidomycin in DMSO-*d6*.**

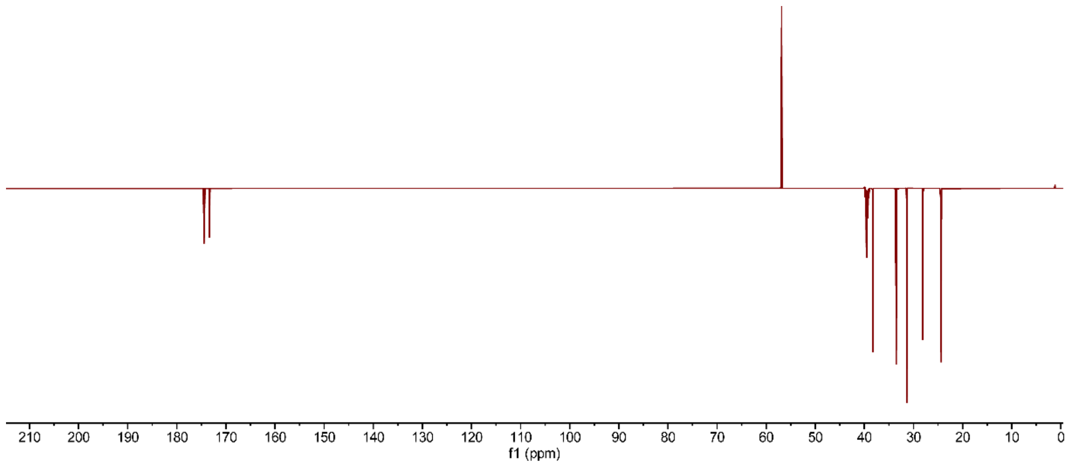


**Figure S39.** $^{13}$**C DEPTQ NMR of acidomycin in DMSO-*d6*.**

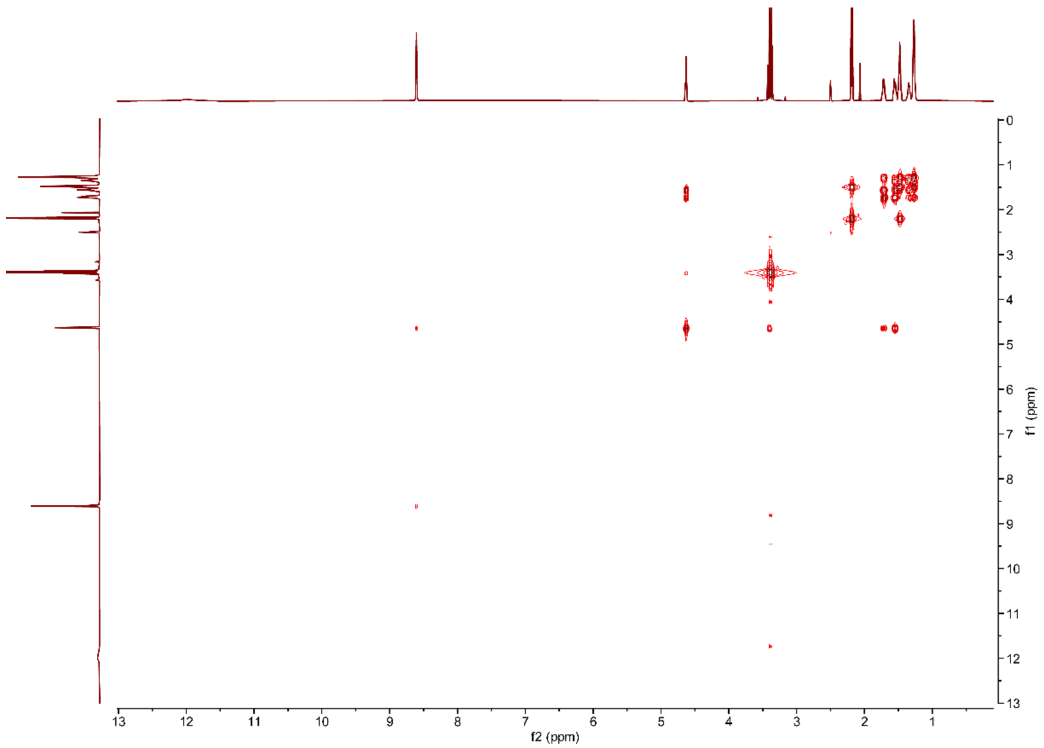


**Figure S40.** $^{1}$**H-**$^{1}$**H COSY NMR of acidomycin in DMSO-*d6*.**

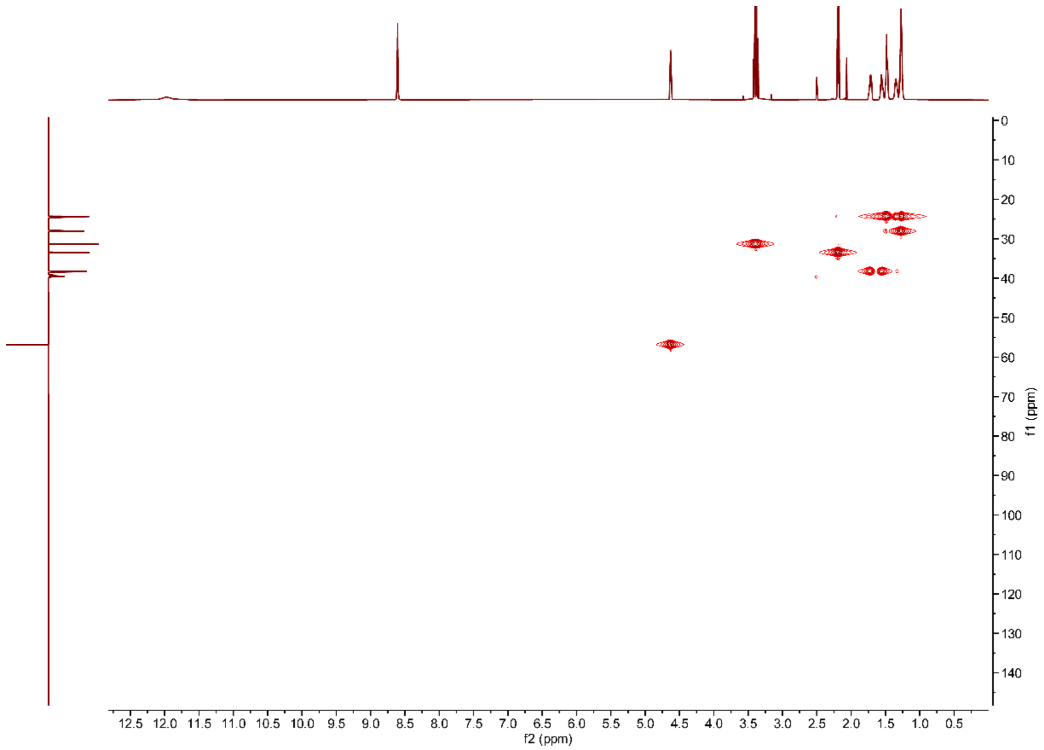


**Figure S41.** **$^{1}$H-$^{13}$C HSQC NMR of acidomycin in DMSO-*d6*.**

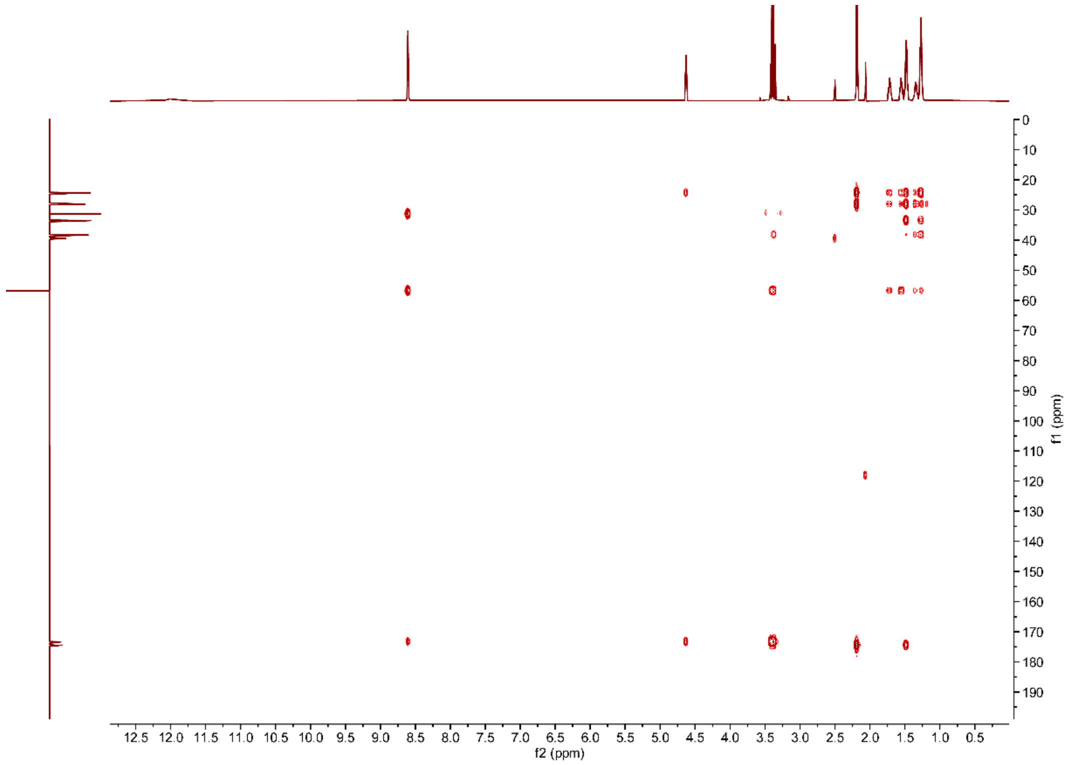


**Figure S42.** **$^{1}$H-$^{13}$C HMBC NMR of acidomycin in DMSO-*d6*.**

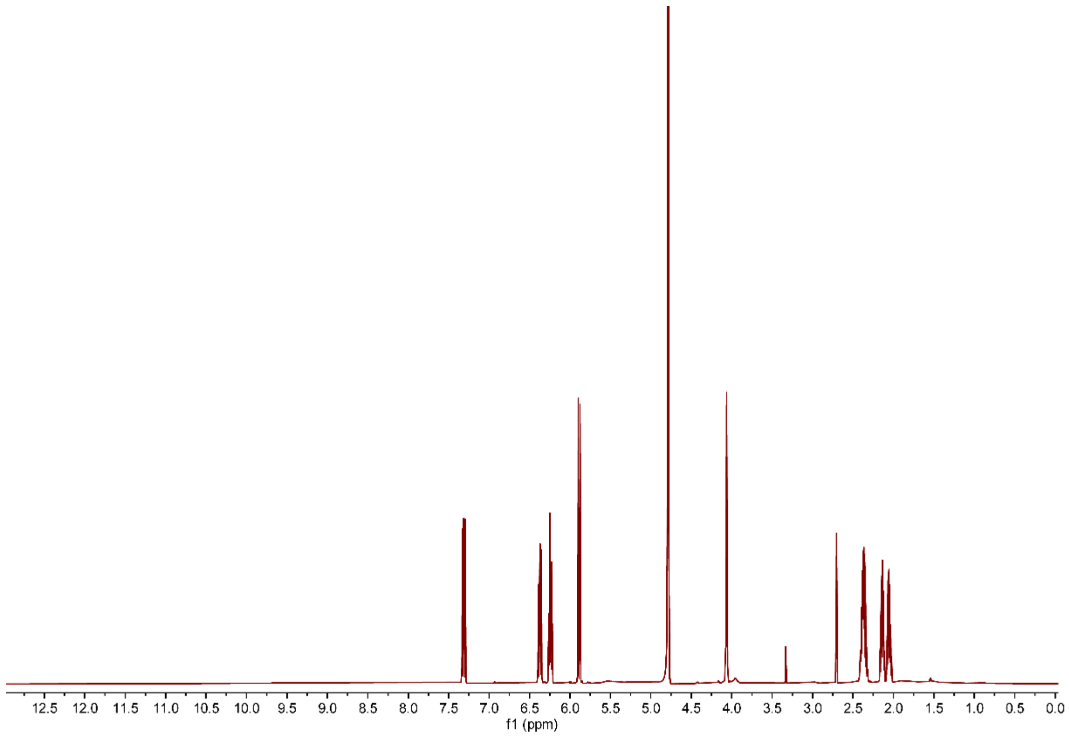


**Figure S43. $^{1}H$ NMR of dapamycin A in $D_2O$.**

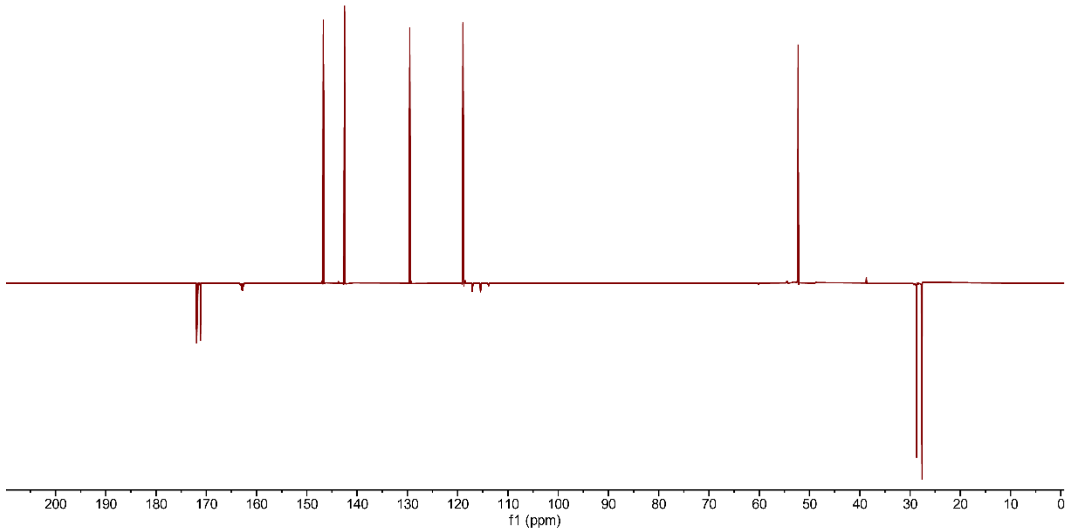


**Figure S44. $^{13}C$ DEPTQ NMR of dapamycin A in $D_2O$.**

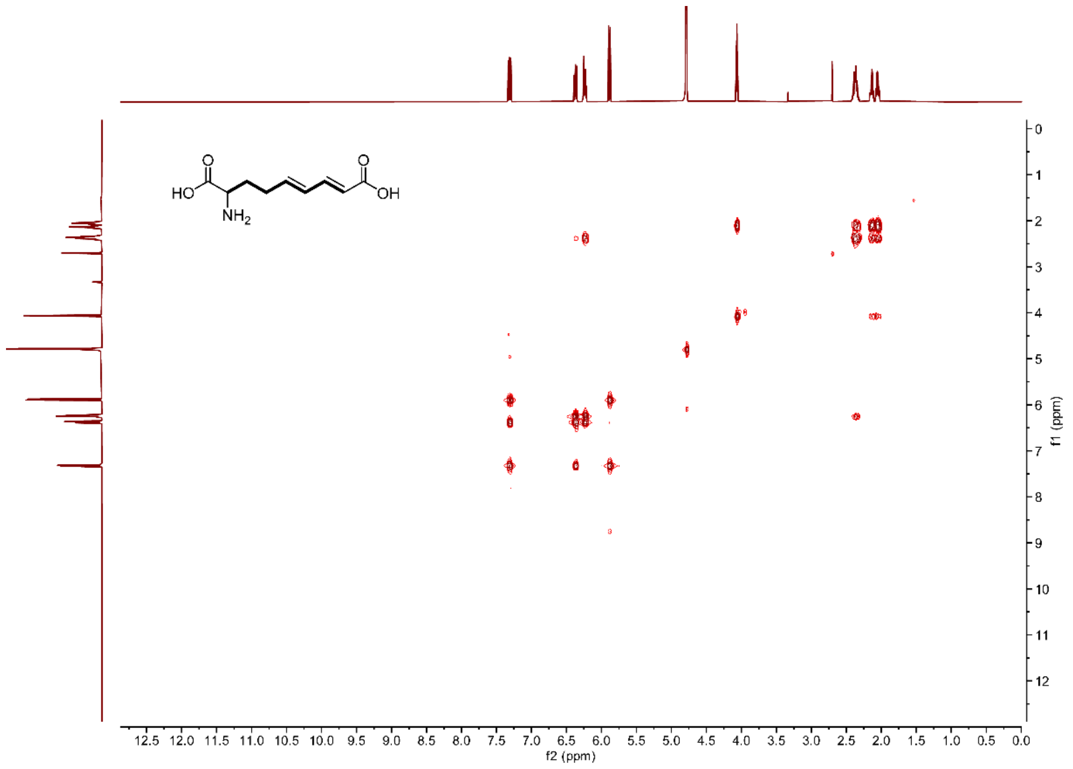


**Figure S45.** $^{1}H$-$^{1}H$ **COSY NMR of dapamycin A in** $D_2O$**.**

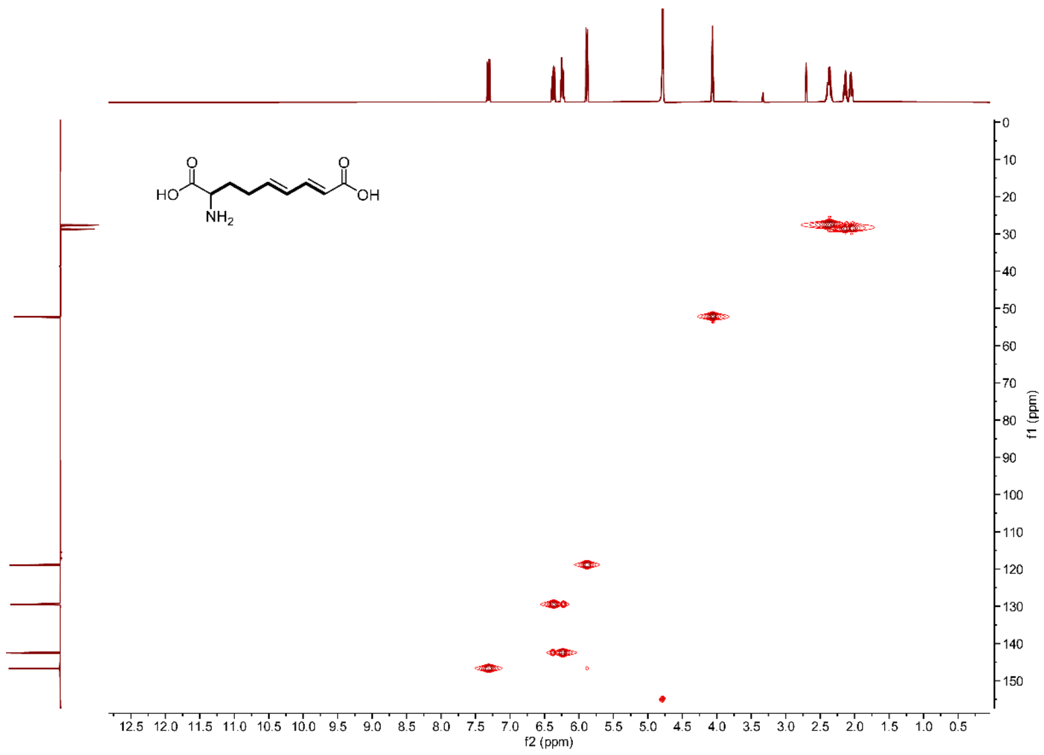


**Figure S46.** $^{1}H$-$^{13}C$ **HSQC NMR of dapamycin A in** $D_2O$**.**

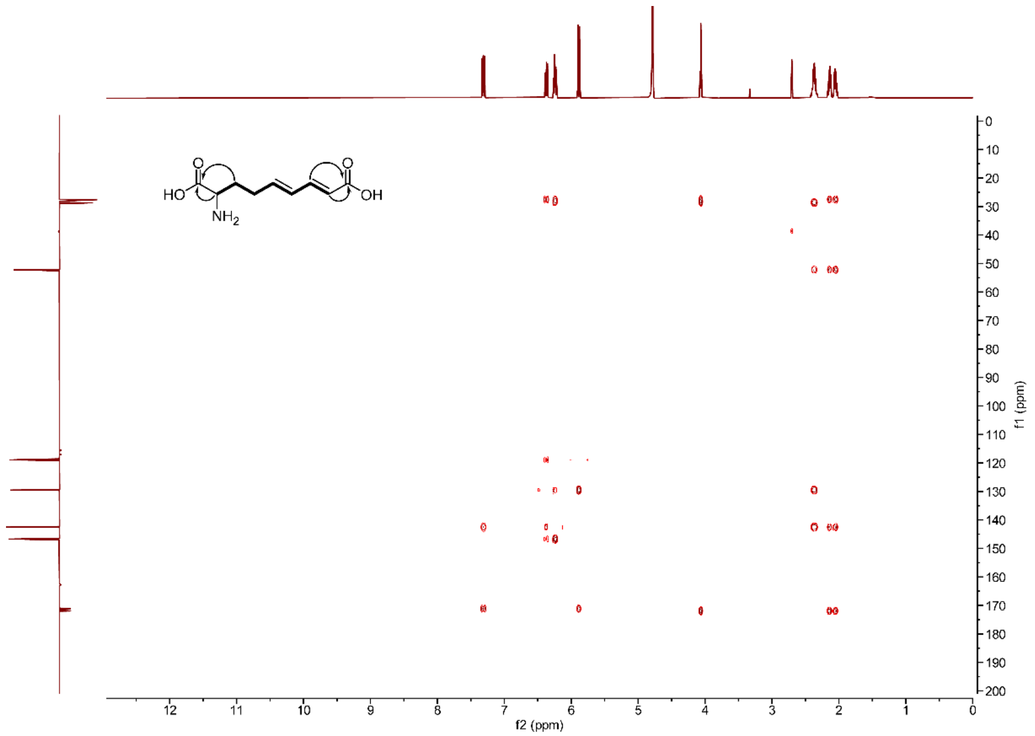


**Figure S47. $^{1}$H-$^{13}$C HMBC NMR of dapamycin A in $D_2O$.**

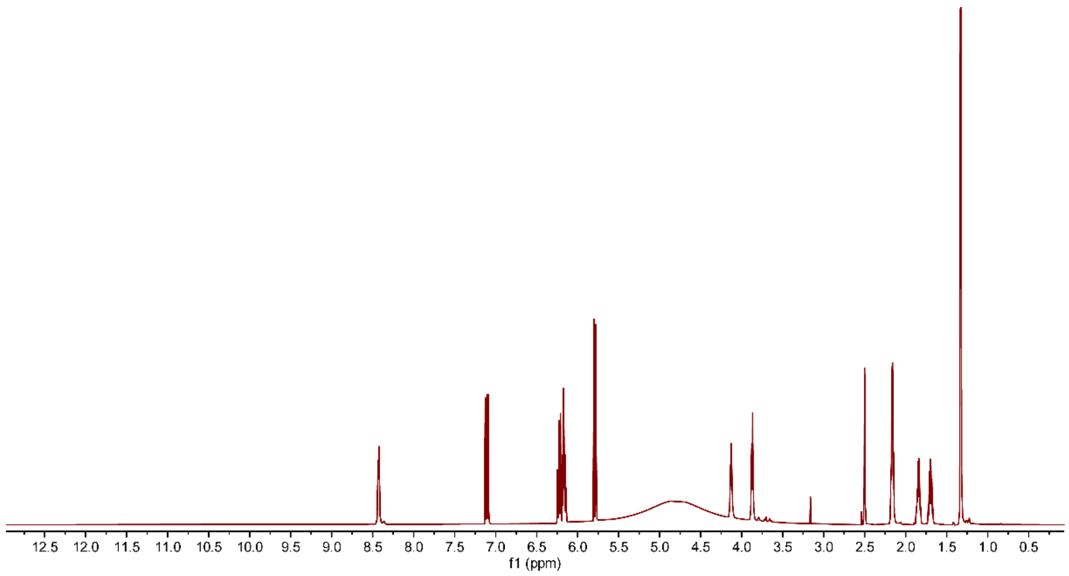


**Figure S48. $^{1}$H NMR of dapamycin B in DMSO-*d6*.**

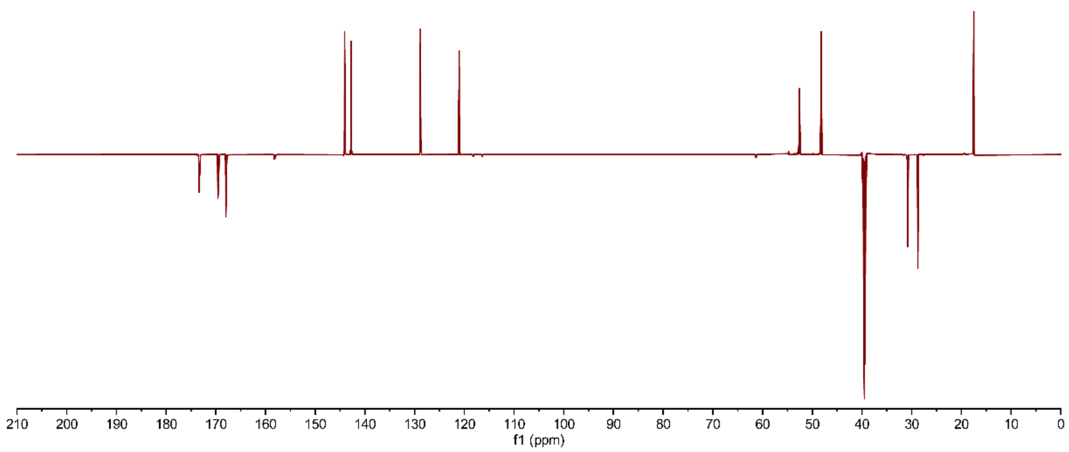


**Figure S49. $^{13}$C DEPTQ NMR of dapamycin B in DMSO-*d6*.**

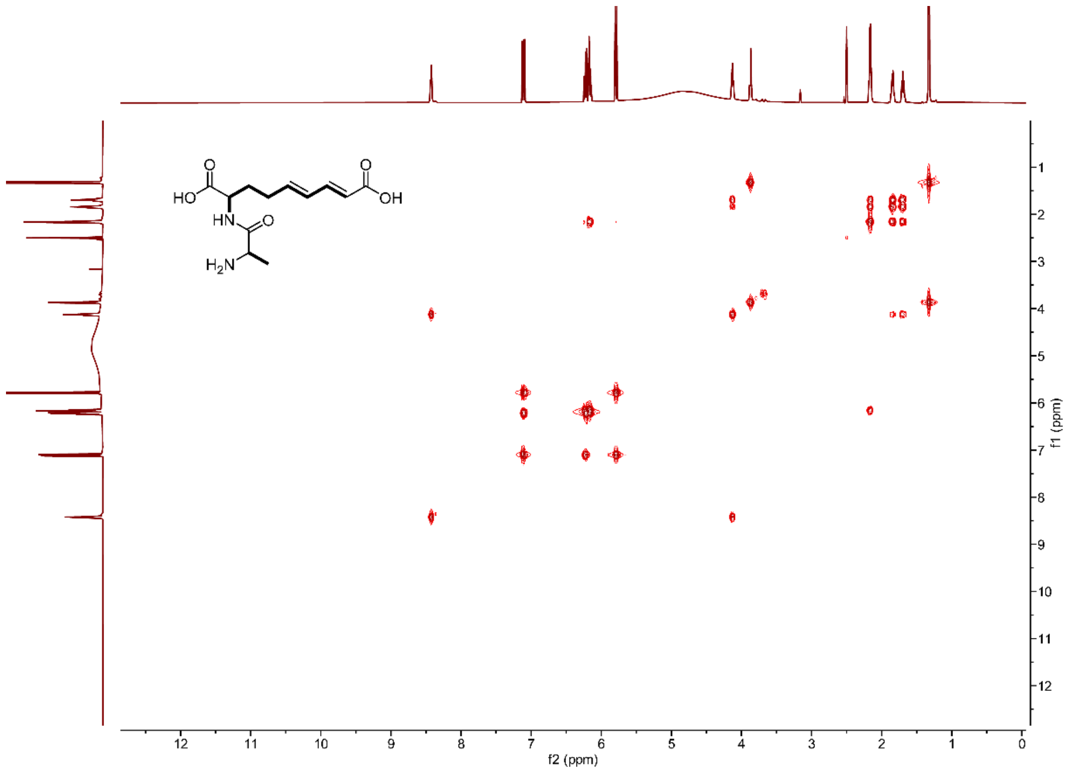


**Figure S50.** **[1]H-[1]H COSY NMR of dapamycin B in DMSO-*d6*.**

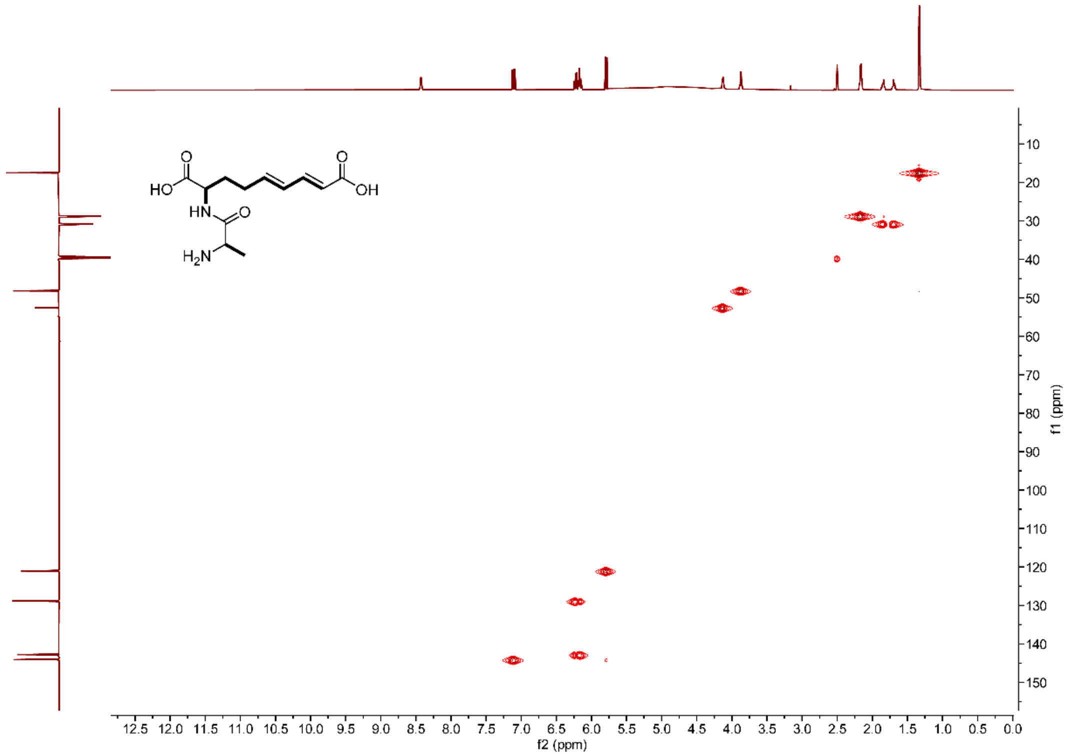


**Figure S51.** **[1]H-[13]C HSQC NMR of dapamycin B in DMSO-*d6*.**

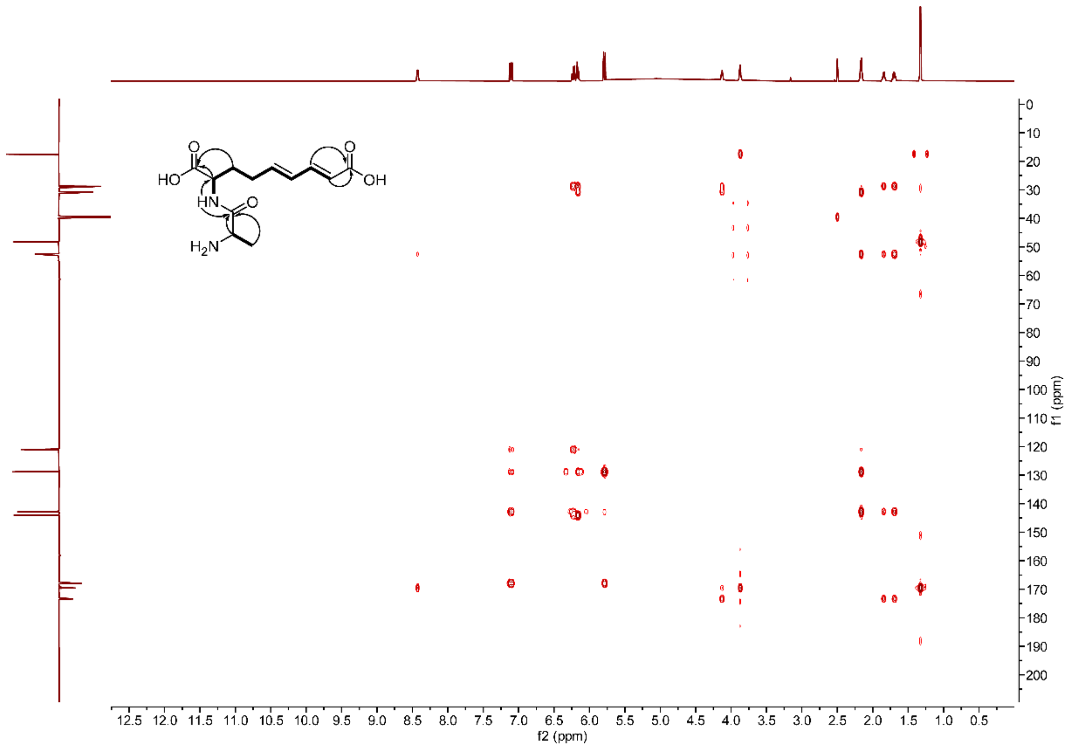


**Figure S52.** $^{1}$**H-**$^{13}$**C HMBC NMR of dapamycin B in DMSO-*d6*.**

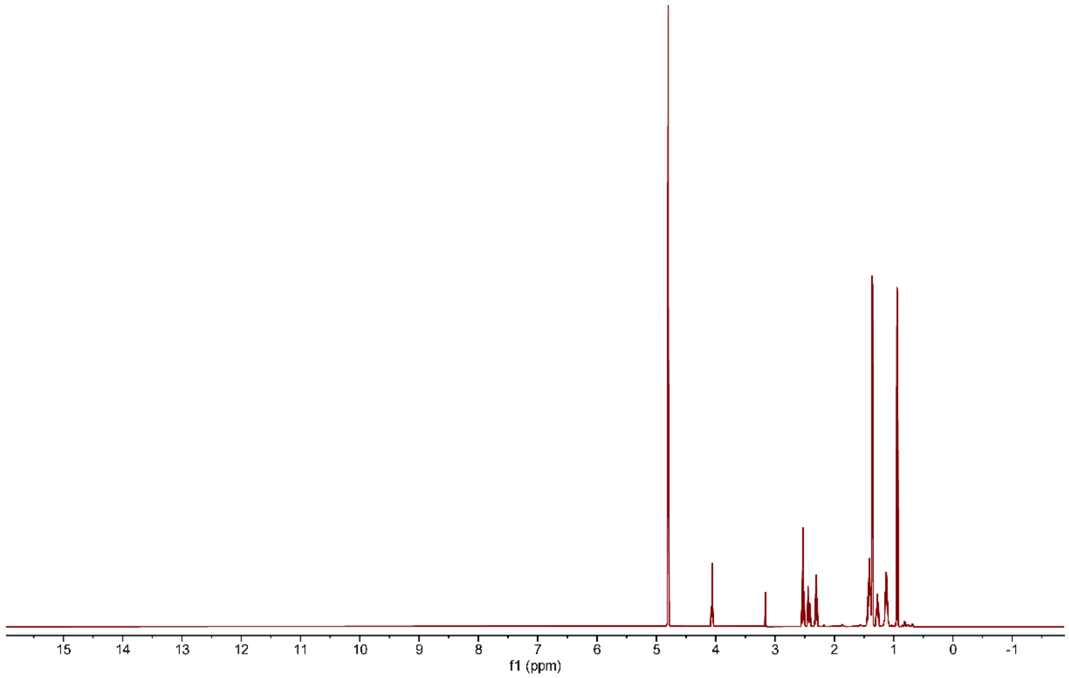


**Figure S53.** $^{1}$**H NMR of α-Me-KAPA in** $D_2O$**.**

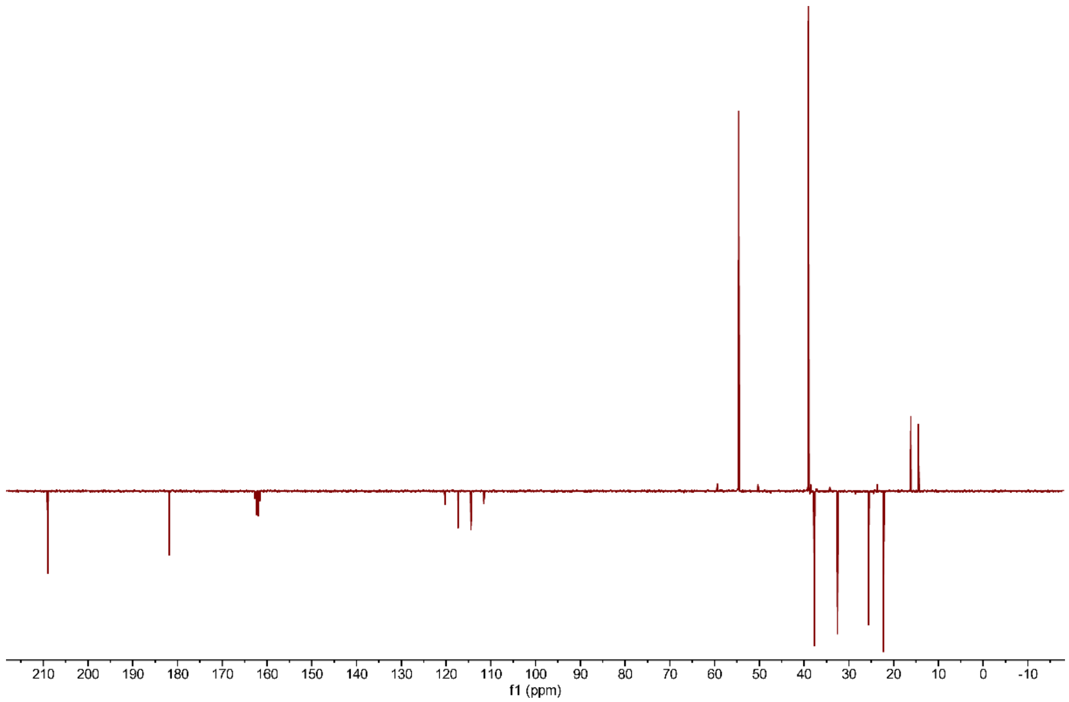


**Figure S54.** $^{13}C$ **DEPTQ NMR of α-Me-KAPA in** $D_2O$**.**

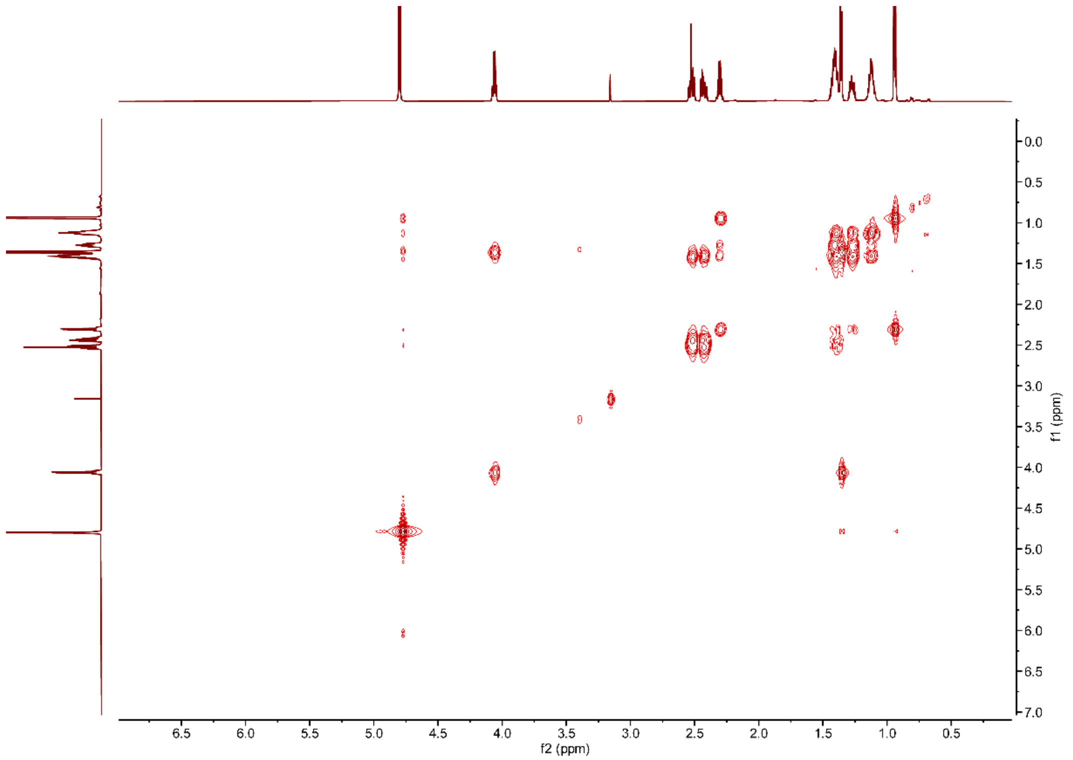


**Figure S55.** $^{1}H$-$^{1}H$ **COSY NMR of α-Me-KAPA in** $D_2O$**.**

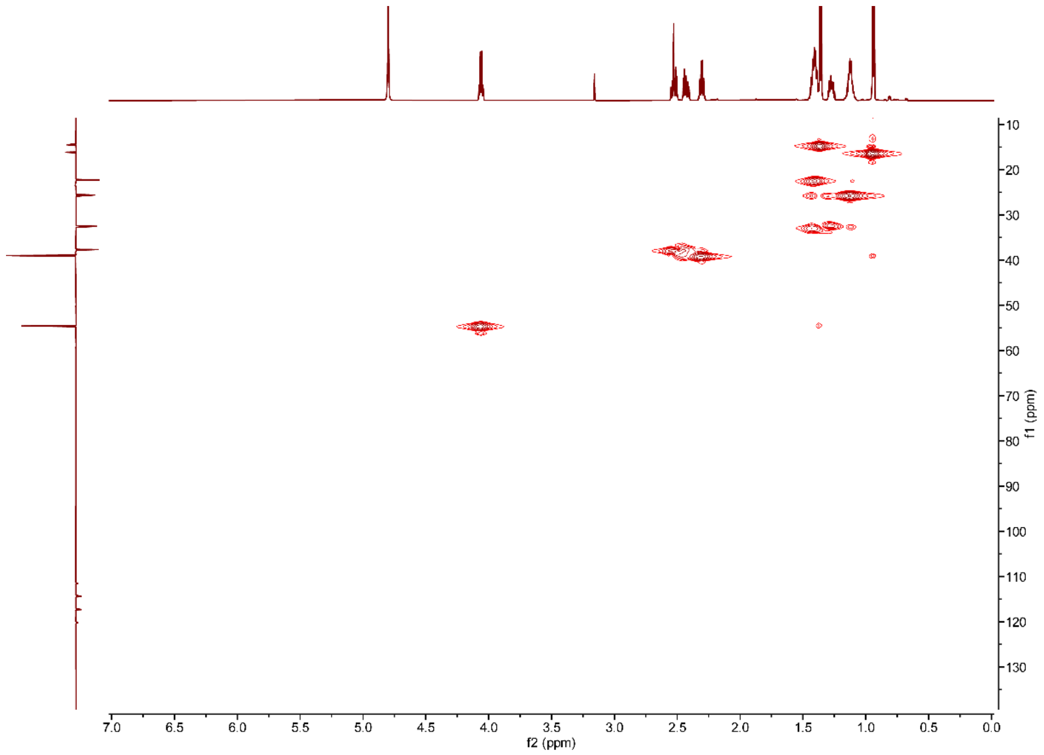


**Figure S56. [1]H-[13]C HSQC NMR of α-Me-KAPA in $D_2O$.**

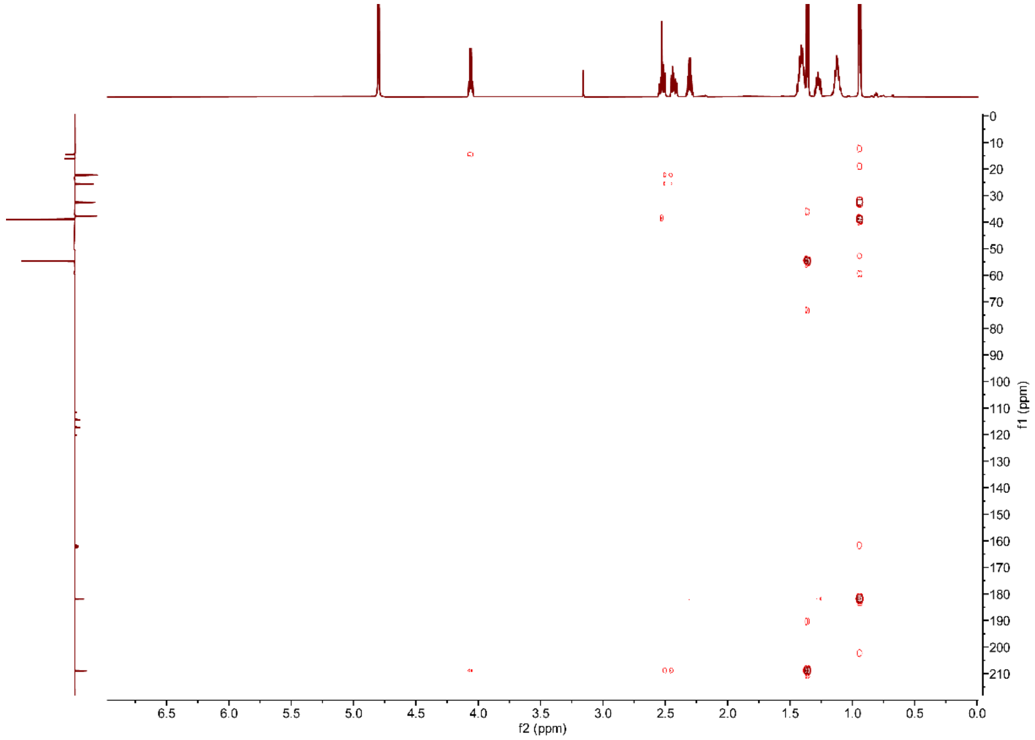


**Figure S57. [1]H-[13]C HMBC NMR of α-Me-KAPA in $D_2O$.**

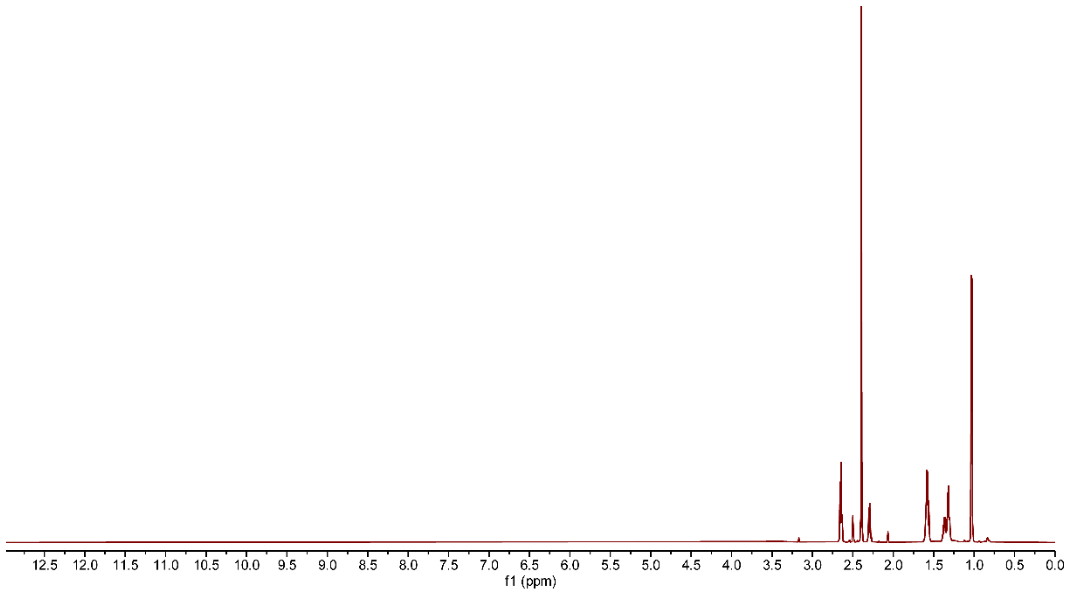


**Figure S58.** **$^{1}H$ NMR of 2,5-dimethyl-3,6-di-(2-methylhexanoic acyl)pyrazine in DMSO-*d6*.**

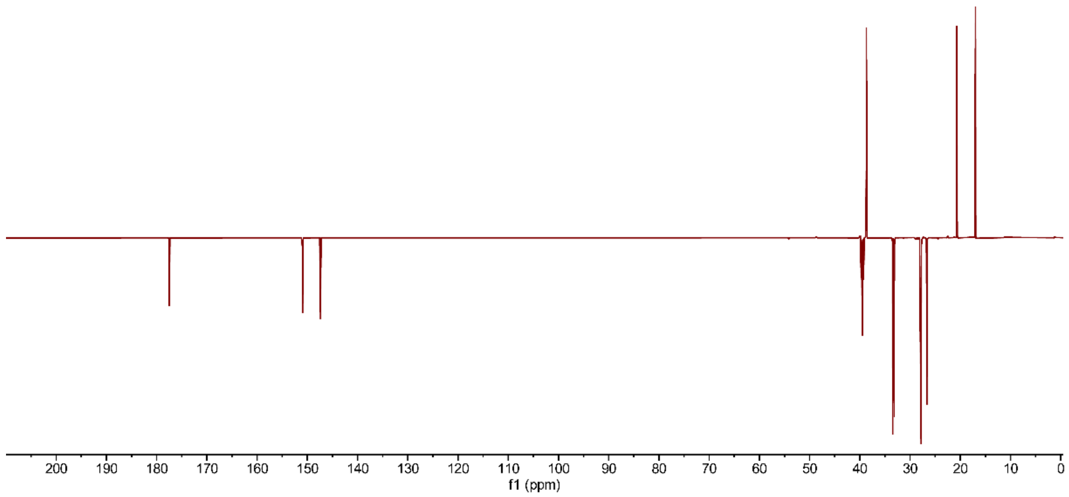


**Figure S59.** **$^{13}C$ DEPTQ NMR of 2,5-dimethyl-3,6-di-(2-methylhexanoic acyl)pyrazine in DMSO-*d6*.**

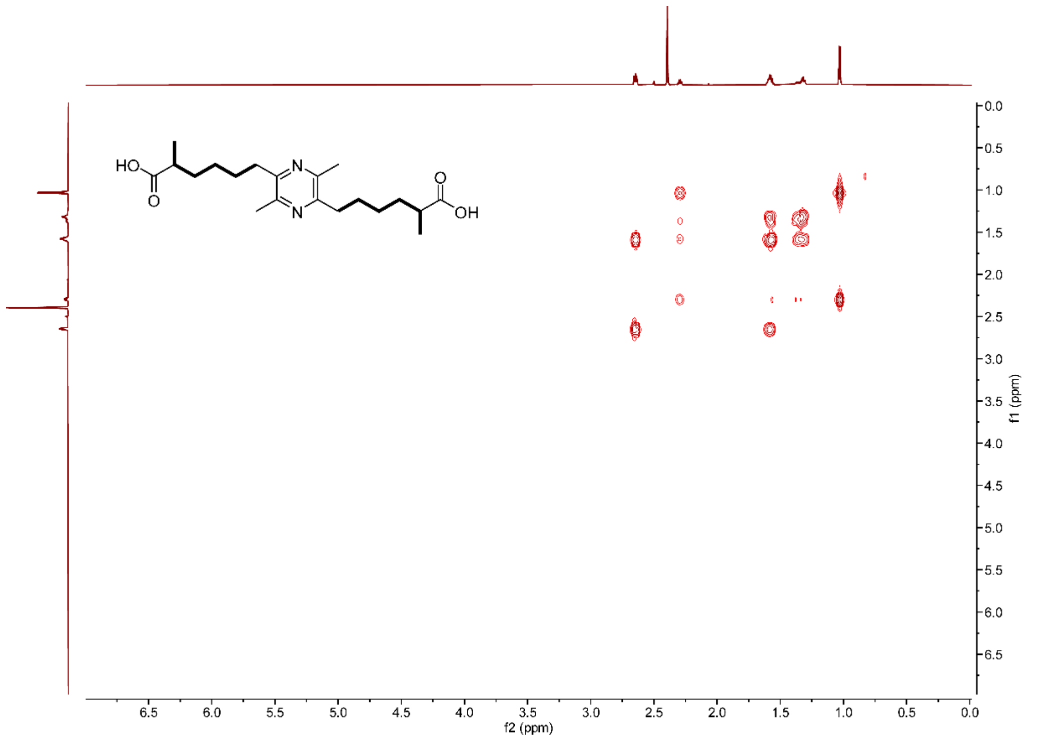


**Figure S60. $^{1}$H-$^{1}$H COSY NMR of 2,5-dimethyl-3,6-di-(2-methylhexanoic acyl)pyrazine in DMSO-*d6*.**

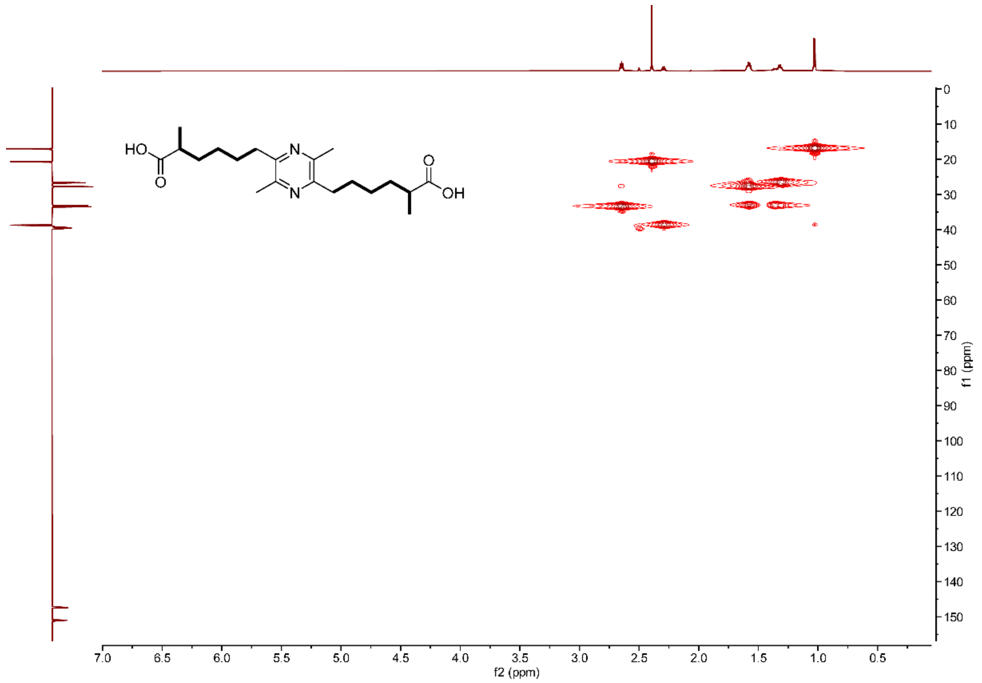


**Figure S61. $^{1}$H-$^{13}$C HSQC NMR of 2,5-dimethyl-3,6-di-(2-methylhexanoic acyl)pyrazine in DMSO-*d6*.**

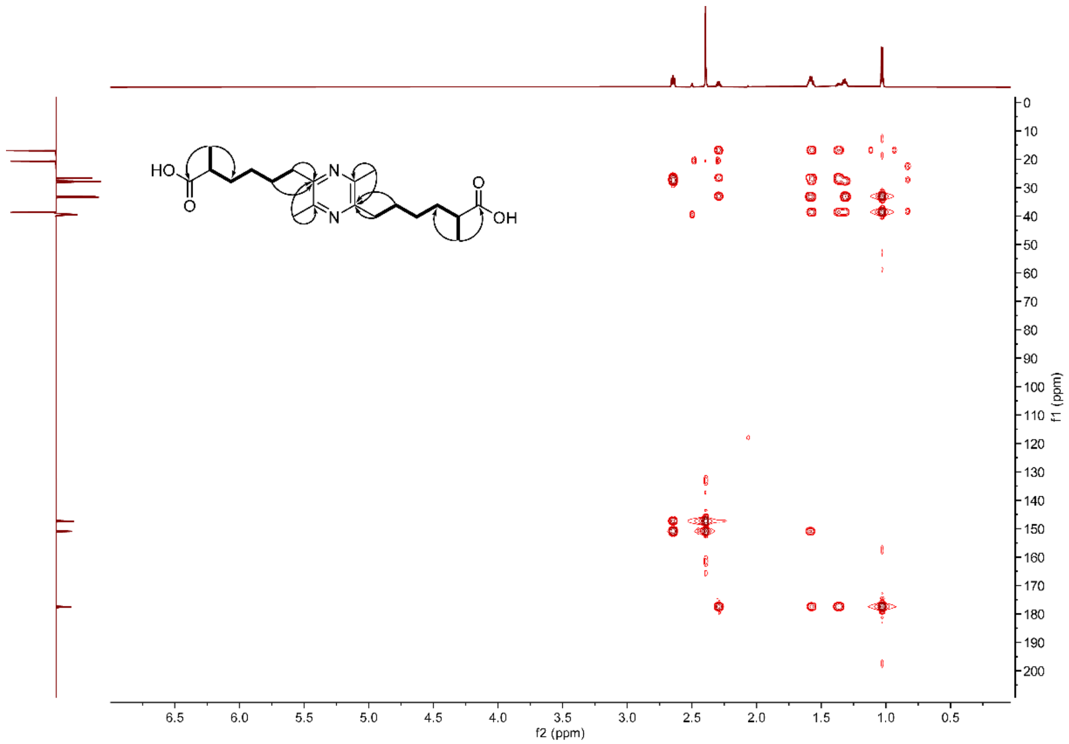


**Figure S62.** $^{1}H$-$^{13}C$ **HMBC NMR of 2,5-dimethyl-3,6-di-(2-methylhexanoic acyl)pyrazine in DMSO-*d6*.**

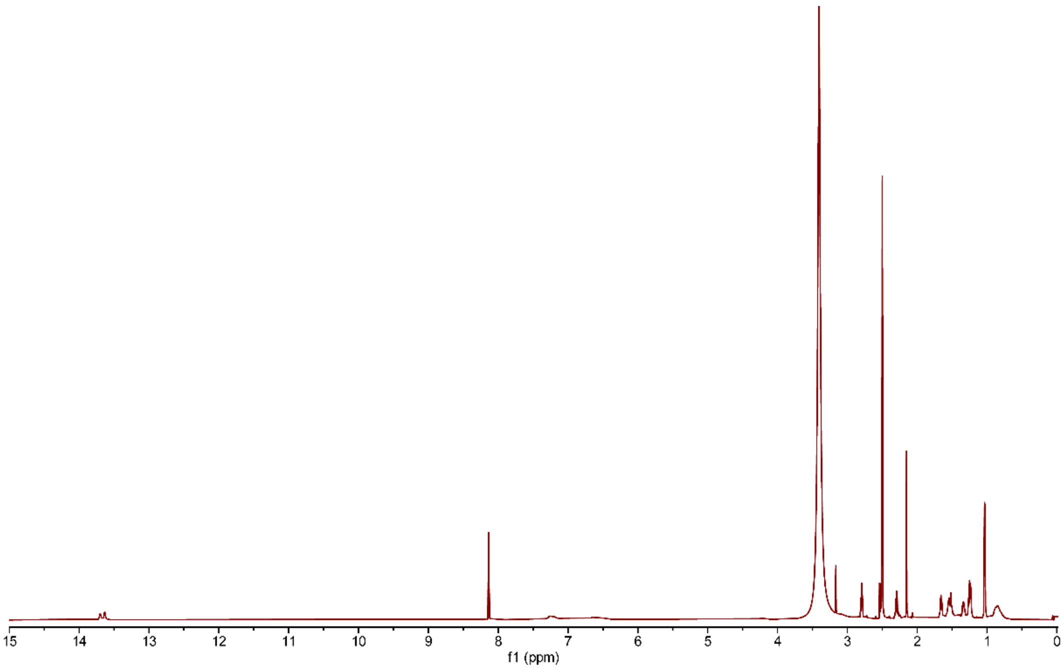


**Figure S63.** $^{1}H$ **NMR of 2,5-di-(2-methylhexanoic acyl)-3-methylimidazole in DMSO-*d6*.**

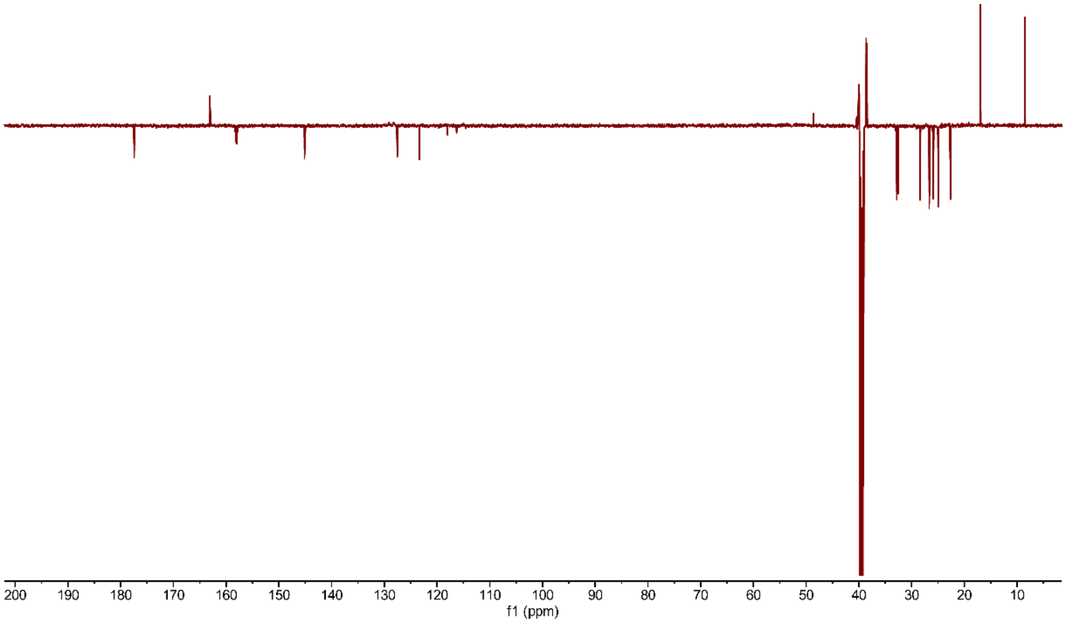


**Figure S64.** $^{13}C$ **DEPTQ NMR of 2,5-di-(2-methylhexanoic acyl)-3-methylimidazole in DMSO-*d6*.**

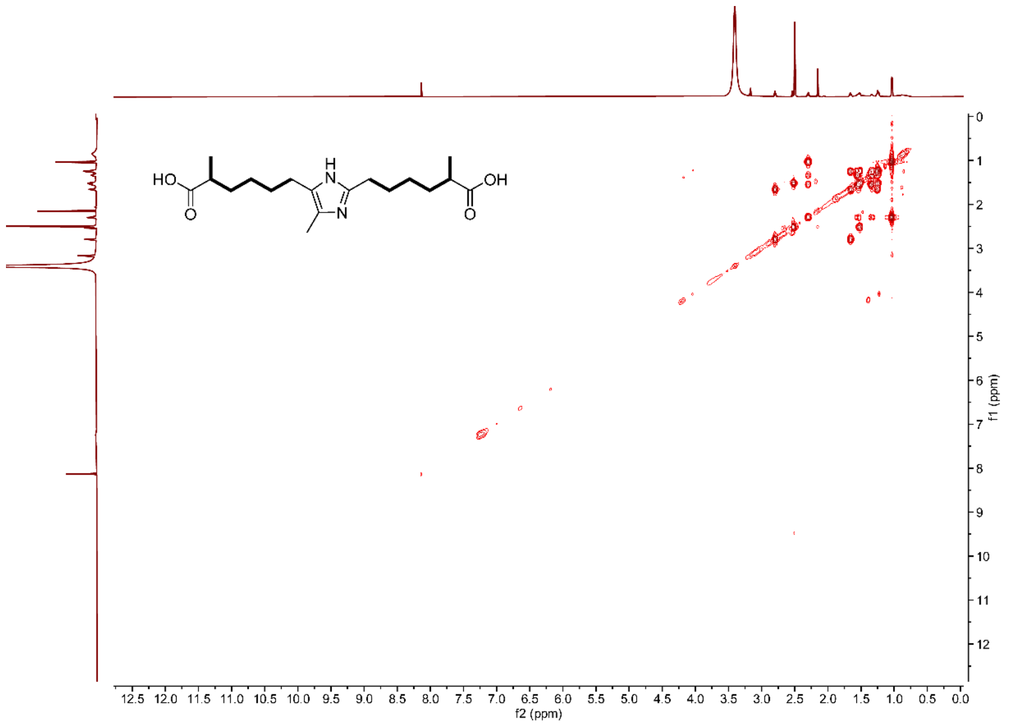

**Figure S65.** $^{1}H$-$^{1}H$ **COSY NMR of 2,5-di-(2-methylhexanoic acyl)-3-methylimidazole in DMSO-*d6*.**

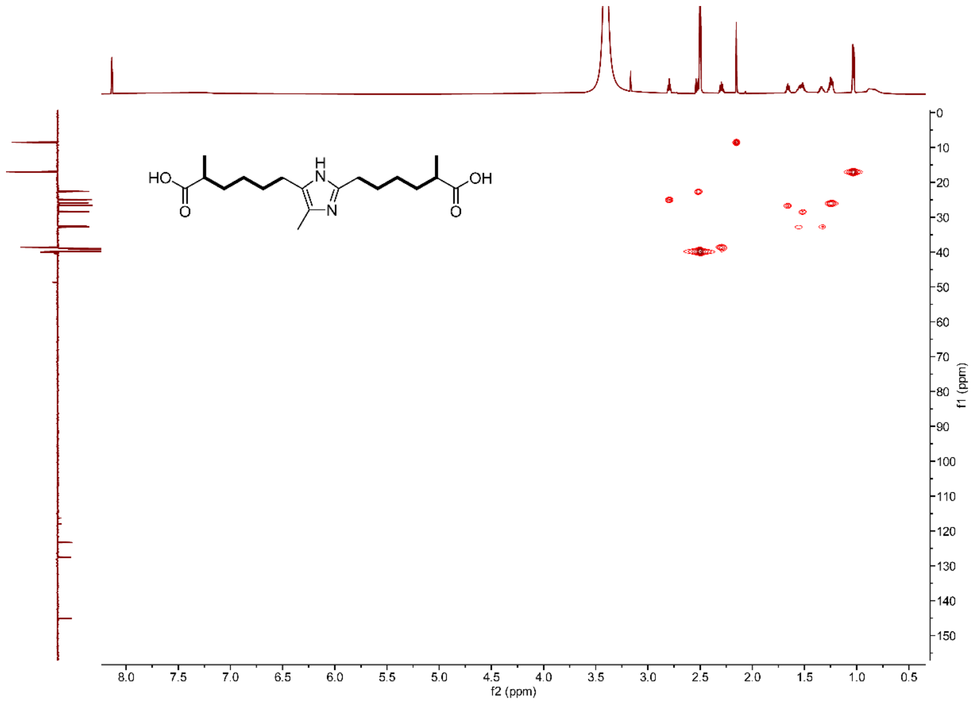

**Figure S66.** $^{1}H$-$^{13}C$ **HSQC NMR of 2,5-di-(2-methylhexanoic acyl)-3-methylimidazole in DMSO-*d6*.**

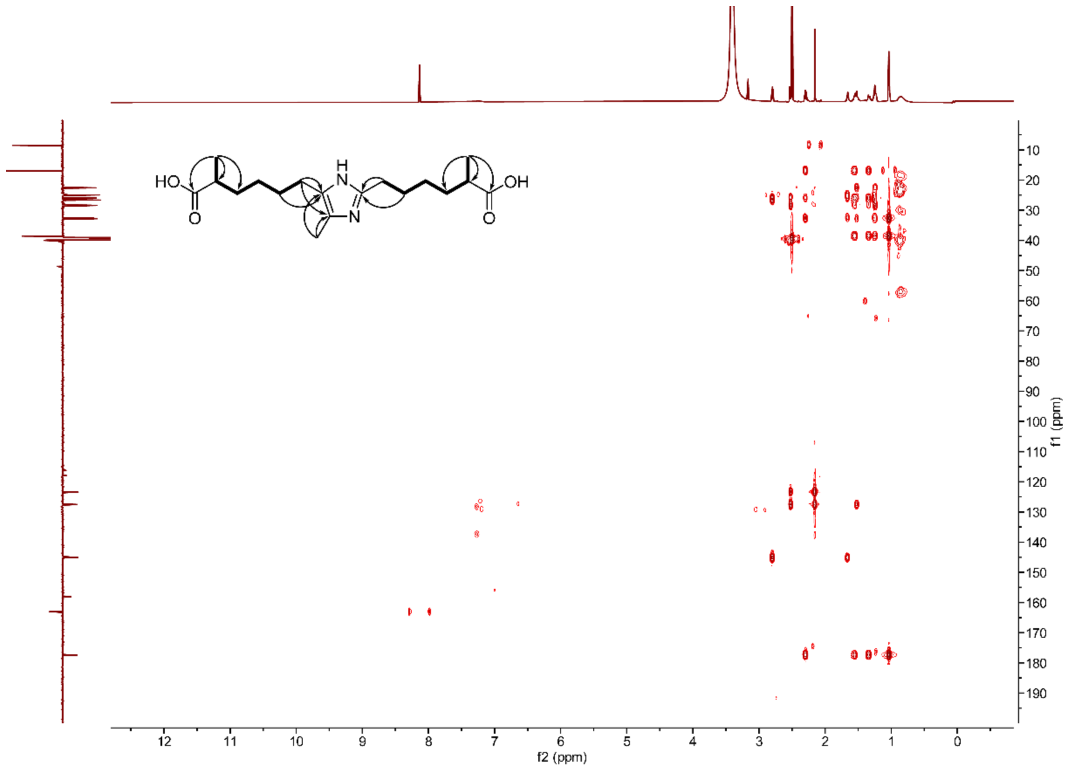


**Figure S67. $^{1}$H-$^{13}$C HMBC NMR of 2,5-di-(2-methylhexanoic acyl)-3-methylimidazole in DMSO-*d6*.**

**Table S1. Annotation of acidomycin-dapamycin-stravidin-α-Me-KAPA megacluster.**

| Genes | Protein (aa) | Homologs | Proposed functions |
|---|---|---|---|
| *orf-2* | 64 | Hypothetical protein | Unknown |
| *orf-1* | 521 | Hypothetical protein | Unknown |
| *avd* | 191 | Streptavidin | Avidin |
| *aciT* | 382 | Major facilitator superfamily transporter AraJ | Transporter |
| *aciA* | 280 | SDR family NAD(P)-dependent oxidoreductase | Oxidoreductase |
| *aciB* | 1564 | NRPS | Peptide synthase |
| *aciC* | 569 | AMP-binding protein | Acyl-CoA ligase |
| *aciD* | 451 | Cytochrome P450 | P450 |
| *aciE* | 235 | 4'-phosphopantetheinyl transferase PptT | PPTase |
| *dapT* | 422 | Major facilitator superfamily transporter | Transporter |
| *dapC* | 250 | α/β fold hydrolase GrsT | Thioesterase |
| *dapB* | 1805 | Type I PKS | Polyketide synthase |
| *dapA* | 1700 | NRPS | Peptide synthase |
| *svnA* | 405 | 3-deoxy-D-arabino-heptulosonate 7-phosphate synthase | DAHP synthase |
| *svnB* | 423 | Major facilitator superfamily transporter | Transporter |
| *svnC* | 456 | Glutathione synthase RimK | Peptide ligase |
| *svnD* | 302 | Branched chain amino acid aminotransferase IlvE | Aminotransferase |
| *svnE* | 355 | Isocitrate/isopropylmalate dehydrogenase LeuB | 3-isopropylmalate dehydrogenase |
| *svnF* | 675 | 3-isopropylmalate dehydratase | Isopropylmalate isomerase |
| *svnG* | 415 | GNAT family N-acetyltransferase | Acetyltransferase |
| *svnH* | 257 | Class I SAM-dependent methyltransferase | Methyltransferase |
| *svnI* | 386 | Isopropylmalate/homocitrate/citramalate synthases LeuA | 2-isopropylmalate synthase |
| *svnJ* | 195 | Hypothetical protein | Unknown |
| *svnK* | 106 | Type II chorismate mutase | 4-amino-4-deoxychorismate mutase |
| *svnL* | 312 | Histone deacetylase | Deacetylase |
| *svnM* | 162 | GNAT family N-acetyltransferase | Acetyltransferase |
| *svnN* | 669 | Aminodeoxychorismate synthase | Aminodeoxychorismate synthase |
| *kapT* | 408 | Major facilitator superfamily transporter AraJ | Transporter |
| *kapA* | 80 | Acyl carrier protein | ACP |
| *kapB* | 393 | β-ketoacyl-ACP synthase II FabF | Ketosynthase |
| *kapC* | 387 | 8-amino-7-oxononanoate synthase BioF | 8-amino-7-oxononanoate synthase |
| *kapD* | 513 | AMP-binding protein | Acyl-CoA ligase |
| *kapE* | 399 | Acetyl-CoA C-acetyltransferase | Acetyltransferase |
| *kapF* | 414 | Cytochrome P450 | P450 |

| | | | |
|---|---|---|---|
| *avd* | 186 | Streptavidin | avidin |
| *orf+1* | 1281 | WD40 domain repeat containing protein | Unknown |
| *orf+2* | 120 | Hypothetical protein | Unknown |

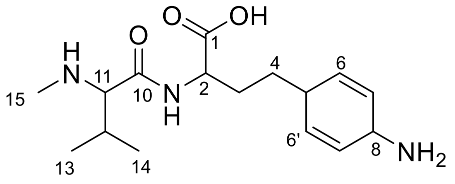

stravidin S2

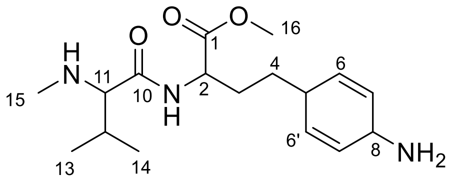

*O*-Me-stravidin S2

**Table S2. $^{1}$H and $^{13}$C NMR data of stravidin S2 and *O*-Me-S2 in $D_2O$.**

| | Stravidin S2 | | *O*-Me-Stravidin S2 | |
|---|---|---|---|---|
| No. | $\delta_H$ | $\delta_C$ | $\delta_H$ | $\delta_C$ |
| 1 | | 177.0 | | 173.4 |
| 2 | 4.27 (m) | 55.1 | 4.53 (dd, *J* = 5.1, 9.0 Hz) | 53.2 |
| 3 | 1.86 (m), 1.74 (m) | 27.9 | 1.97 (m), 1.83 (m) | 27.0 |
| 4 | 1.60 (2H, m) | 29.5 | 1.65 (2H, dt, *J* = 7.7, 8.3 Hz) | 29.5 |
| 5 | 2.90 (m) | 34.5 | 2.94 (m) | 34.4 |
| 6, 6’ | 6.11 (2H, br dd, *J* = 10.5, 11.2 Hz) | 135.5 | 6.14 (2H, br dd, *J* = 10.4, 11.2 Hz) | 135.2 |
| 7, 7’ | 5.85 (2H, br d, *J* = 10.5 Hz) | 120.5 | 5.88 (2H, br d, *J* = 10.4 Hz) | 120.6 |
| 8 | 4.41 (m) | 44.9 | 4.44 (m) | 44.9 |
| 10 | | 167.2 | | 167.9 |
| 11 | 3.78 (d, *J* = 5.7 Hz) | 67.1 | 3.82 (d, *J* = 5.8 Hz) | 66.9 |
| 12 | 2.25 (d sept, *J* = 5.8, 7.2 Hz) | 29.9 | 2.30 (d sept, *J* = 5.8, 6.9 Hz) | 29.9 |
| 13 | 1.09 (3H, d, *J* = 7.2 Hz) | 17.8 | 1.13 (3H, d, *J* = 6.9 Hz) | 17.6 |
| 14 | 1.02 (3H, d, *J* = 7.2 Hz) | 17.4 | 1.06 (3H, d, *J* = 6.9 Hz) | 17.0 |
| 15 | 2.69 (3H, s) | 32.2 | 2.72 (3H, s) | 32.1 |
| 16 | | | 3.79 (3H, s) | 52.9 |

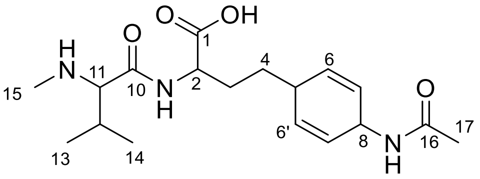


stravidin S4

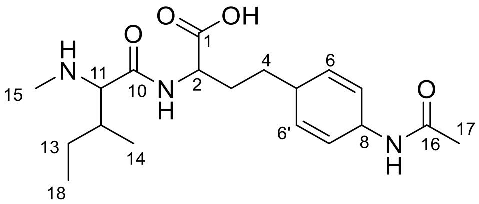


stravidin S5

**Table S3. [1]H and [13]C NMR Data of stravidin S4 and S5 in $D_2O$.**

| | Stravidin S4 | | Stravidin S5 | |
|---|---|---|---|---|
| No. | $\delta_H$ | $\delta_C$ | $\delta_H$ | $\delta_C$ |
| 1 | | 177.5 | | 177.6 |
| 2 | 4.24 (dd, *J* = 5.0, 8.1 Hz) | 55.6 | 4.22 (dd, *J* = 5.2, 7.6 Hz) | 55.7 |
| 3 | 1.82 (m), 1.70 (m) | 27.7 | 1.82 (m), 1.70 (m) | 27.9 |
| 4 | 1.55 (2H, m) | 29.7 | 1.53 (2H, m) | 29.6 |
| 5 | 2.83 (m) | 34.0 | 2.83 (m) | 34.0 |
| 6, 6’ | 5.88 (2H, br dd, *J* = 9.2, 9.6 Hz) | 131.8 | 5.87 (2H, br dd, *J* = 9.4, 9.5 Hz) | 131.8 |
| 7, 7’ | 5.70 (2H, br d, *J* = 9.2 Hz) | 124.5 | 5.70 (2H, br d, *J* = 9.4 Hz) | 124.4 |
| 8 | 4.81 (m, overlapped) | 43.5 | 4.81 (m, overlapped) | 43.5 |
| 10 | | 166.9 | | 166.8 |
| 11 | 3.74 (d, *J* = 6.0 Hz) | 67.2 | 3.81 (d, *J* = 5.7 Hz) | 66.4 |
| 12 | 2.25 (d sept, *J* = 6.0, 6.7 Hz) | 29.8 | 2.01 (m, overlapped) | 36.1 |
| 13 | 1.09 (3H, d, *J* = 6.7 Hz) | 17.6 | 1.30 (m), 1.60 (m) | 24.7 |
| 14 | 1.03 (3H, d, *J* = 6.7 Hz) | 17.4 | 1.00 (3H, d, *J* = 7.0 Hz) | 13.8 |
| 15 | 2.69 (3H, s) | 32.0 | 2.69 (3H, s) | 32.1 |
| 16 | | 173.0 | | 172.6 |
| 17 | 1.99 (3H, s) | 21.9 | 1.99 (3H, s) | 21.9 |
| 18 | | | 0.95 (3H, t, *J* = 7.4 Hz) | 10.5 |

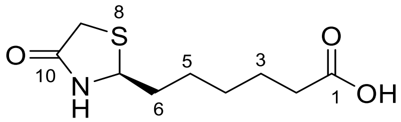


Acidomycin

**Table S4. [1]H and [13]C NMR data of Acidomycin in DMSO-$d_6$.**

| No. | $\delta_H$ | $\delta_C$ | No | $\delta_H$ | $\delta_C$ |
|---|---|---|---|---|---|
| 1 | | 174.5 | 6 | 1.55 (m), 1.72 (m) | 38.3 |
| 2 | 2.19 (2H, t, *J* = 7.3 Hz) | 33.5 | 7 | 4.63 (dd, J = 6.1, 6.2 Hz) | 56.9 |
| 3 | 1.48 (2H, quint, *J* = 7.3 Hz) | 24.4 | 9 | 3.36 (br d, J =15.5 Hz)<br>3.41 (dd, J = 1.7, 15.5 Hz) | 31.4 |
| 4 | 1.27 (2H, m, overlapped) | 28.1 | 10 | | 173.3 |
| 5 | 1.35 (m), 1.28 (m) | 24.4 | 11-NH | 8.61 (br s) | |

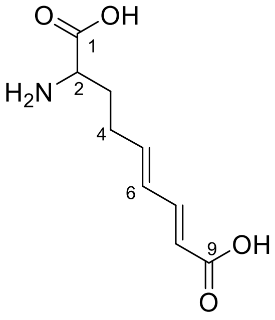

Dapamycin A

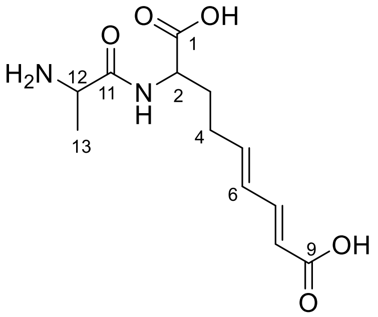

Dapamycin B

**Table S5. $^{1}H$ and $^{13}C$ NMR data of Dapamycin A in $D_2O$ and Dapamycin B in DMSO-$d_6$.**

| | Dapamycin A | | Dapamycin B | |
|---|---|---|---|---|
| No. | $\delta_H$ | $\delta_C$ | $\delta_H$ | $\delta_C$ |
| 1 | | 171.9 | | 173.4 |
| 2 | 4.06 (dd, $J$ = 6.4, 6.3 Hz) | 55.6 | 4.13 (m) | 52.6 |
| 3 | 2.13 (m), 2.05 (m) | 28.7 | 1.84 (m), 1.69 (m) | 30.8 |
| 4 | 2.37 (2H, m) | 27.6 | 2.16 (2H, m) | 28.7 |
| 5 | 6.24 (dt, $J$ = 15.1, 6.8 Hz) | 142.4 | 6.16 (dt, $J$ = 15.1, 5.6Hz) | 142.7 |
| 6 | 6.37 (dd, $J$ = 15.1, 10.9 Hz) | 129.5 | 6.23 (dd, $J$ = 15.1, 10.7 Hz) | 128.9 |
| 7 | 7.31 (dd, $J$ = 15.3, 10.9 Hz) | 146.7 | 7.11 (dd, $J$ = 15.4, 10.7 Hz) | 144.1 |
| 8 | 5.89 (d, $J$ = 15.3 Hz) | 118.9 | 5.79 (d, $J$ = 15.4 Hz) | 121.1 |
| 9 | | 171.2 | | 168.0 |
| 10-NH | | | 8.43 (d, $J$ = 8.1 Hz) | |
| 11 | | | | 169.6 |
| 12 | | | 3.87 (q, $J$ = 6.9 Hz) | 48.2 |
| 13 | | | 1.33 (3H, d, $J$ = 6.9 Hz) | 17.5 |

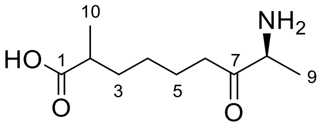


α-Me-KAPA

**Table S6. $^{1}H$ and $^{13}C$ NMR data of α-Me-KAPA in $D_2O$.**

| No. | $\delta_H$ | $\delta_C$ | No | $\delta_H$ | $\delta_C$ |
|---|---|---|---|---|---|
| 1 | | 181.8 | 6 | 2.52 (dt, *J* = 17.8, 7.0 Hz)<br>2.43 (dt, *J* = 17.8, 7.1 Hz) | 37.7 |
| 2 | 2.31 (sext, *J* = 7.0 Hz) | 39.0 | 7 | | 208.9 |
| 3 | 1.27 (m, overlapped)<br>1.41 (m, overlapped) | 32.5 | 8 | 4.06 (q, *J* = 7.4 Hz) | 54.6 |
| 4 | 1.12 (2H, m) | 25.6 | 9 | 1.36 (3H, d, *J* = 7.4 Hz) | 14.5 |
| 5 | 1.41 (2H, m, overlapped) | 22.2 | 10 | 0.94 (3H, d, *J* = 7.2 Hz) | 16.1 |

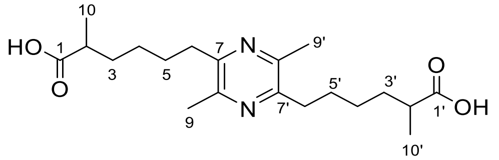

2,5-dimethyl-3,6-di-(2-methylhexanoic acyl)pyrazine

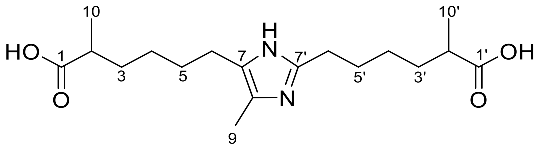

2,5-di-(2-methylhexanoic acyl)-3-methylimidazole

**Table S7. $^{1}H$ and $^{13}C$ NMR Data of 2,5-dimethyl-3,6-di-(2-methylhexanoic acyl)pyrazine and 2,5-di-(2-methylhexanoic acyl)-3-methylimidazole in DMSO-$d_6$.**

| 2,5-dimethyl-3,6-di-(2-methylhexanoic acyl) pyrazine | | | 2,5-di-(2-methylhexanoic acyl)-3-methylimidazole | | |
|---|---|---|---|---|---|
| No. | $\delta_H$ | $\delta_C$ | No. | $\delta_H$ | $\delta_C$ |
| 1, 1’ | | 177.3 | 1 | | |
| 2, 2’ | 2.29 (sext, $J$ = 6.2 Hz) | 38.7 | 2 | 2.30 (m) | 38.6 |
| 3, 3’ | 1.36 (m, overlapped)<br>1.58 (m, overlapped) | 33.1 | 3 | 1.34 (m, overlapped)<br>1.54 (m, overlapped) | 32.8 |
| 4, 4’ | 1.32 (2H, m, overlapped) | 26.7 | 4 | 1.24 (2H, m) | 25.9 |
| 5, 5’ | 1.58 (2H, m, overlapped) | 27.8 | 5 | 1.52 (2H, m, overlapped) | 28.4 |
| 6, 6’ | 2.65 (2H, t, $J$ = 7.6 Hz) | 33.5 | 6 | 2.52 (2H, m, overlapped) | 22.6 |
| 7, 7’ | | 150.9 | 7 | | 127.5 |
| 8, 8’ | | 147.4 | 8 | | 123.3 |
| 9, 9’ | 2.39 (3H, s) | 20.7 | 9 | 2.15 (3H, s) | 8.5 |
| 10, 10’ | 1.03 (d, $J$ = 7.1 Hz) | 17.0 | 10 | 1.03 (d, $J$ = 6.7 Hz) | 17.0 |
| | | | 1’ | | 177.4 |
| | | | 2’ | 2.30 (m) | 38.5 |
| | | | 3’ | 1.34 (m, overlapped)<br>1.54 (m, overlapped) | 32.8 |
| | | | 4’ | 1.24 (2H, m) | 25.9 |
| | | | 5’ | 1.66 (2H, quint, $J$ = 7.6 Hz) | 26.7 |
| | | | 6’ | 2.80 (2H, t, $J$ = 7.6 Hz) | 25.0 |
| | | | 7’ | | 145.0 |
| | | | 10’ | 1.03 (d, $J$ = 6.7 Hz) | 17.0 |

**Table S8. Primers, gblocks and sgRNA guide sequences used in this study.**

| Primers | Sequence (5′-3′) | Description |
|---|---|---|
| svn-gF | tgttctcggacatgttcg | |
| svn-gR | gtggtgatgcggatgac | |
| pCGW-sF | atccatgcttcctgaagattcc | |
| pCGW-sR | attccttggtggtacgaacatc | |
| svn-dF | actcttcacgatcagcac | |
| svn-dR | acgacttcatctgcttcc | |
| Δsvn-F | gttgccgtgcatcgggtcgcacatccagaccacgggatgcccCCTAGGtgcagctcacggtaactgat | *Avr*II site |
| Δsvn-R | cttctcgtagacaactatgactcctacacgtggaacctcttcCCTAGGaggaacttatgagctcagcc | |
| Δsvn2-F | ggtgttggccggcggctggccgtcgaggaagcgctccggacgCCTAGGtgcagctcacggtaactgat | *Avr*II site |
| Δsvn2-R | cttctcgtagacaactatgactcctacacgtggaacctcttcCCTAGGaggaacttatgagctcagcc | |
| Δsvn3-F | gttgccgtgcatcgggtcgcacatccagaccacgggatgcccCCTAGGtgcagctcacggtaactgat | *Avr*II site |
| Δsvn3-R | gacagtgtggacctgttcgaccaggggtatgcgctggacccgCCTAGGaggaacttatgagctcagcc | |
| ΔDS-F | ccgcagcgggatcagcaggacgatcccggcgcccgccagcacCCTAGGtgcagctcacggtaactgat | *Avr*II site |
| ΔDS-R | cttctcgtagacaactatgactcctacacgtggaacctcttcCCTAGGaggaacttatgagctcagcc | |
| ΔSK-F | cggtccgcgcagag | Amplified from pUC18-k*TaSK |
| ΔSK-R | cgtctcacagggtttcttc | |
| ΔDSK-F | ccgcagcgggatcagcaggacgatcccggcgcccgccagcacCCTAGGtgcagctcacggtaactgat | *Avr*II site |
| ΔDSK-R | gacagtgtggacctgttcgaccaggggtatgcgctggacccgCCTAGGaggaacttatgagctcagcc | |
| aci-F | ggtgatgtgatcatggcgcaaaacctctgttggcctgcaactCATATGtgcagctcacggta | Amplified from pUC18-k*TaSK |
| aci-R | tgcgaacgctccgagcgcgagcgtcgagatcttgcgcaacacaactcccccagtcctg | |
| dap-F | acgagcggctacgtcgtcaccggtgtgctgccgcagctgagcCCTAGGtgcagctcacggtaactgat | *Avr*II site |
| dap-R | cttgtagaccgcctccttcgcacagaacagcagccggtcccaCCTAGGaggaacttatgagctcagc | |
| kap-F | ttgctttgccgatgttactt | Amplified from pUC18-aTk*k |
| kap-R | gacacgtccagttcacc | |
| Δsvn-dF | gcacatccagaccacg | |
| Δsvn-dR | acagggatgcatattcgg | |
| Δsvn2-dF | gaggggatcagcgtca | |
| Δsvn2-dR | tcggaagtgtcagaactg | |

| | | |
|---|---|---|
| Δsvn3-dF | catgttggtccatctcgg | |
| Δsvn3-dR | ctcacagggtttcttccc | |
| ΔDS-dF | tgatcaggtacatgttgtgc | |
| ΔDS-dR | acagggatgcatattcgg | |
| ΔSK-dF | cagaccggccttgagta | |
| ΔSK-dR | cgtctcacagggtttcttc | |
| ΔDSK-dF | tgatcaggtacatgttgtgc | |
| ΔDSK-dR | cgtctcacagggtttcttc | |
| aci-dF | cggatcgtcacccagt | |
| aci-dR | gcgatgatgagcggtc | |
| dap-dF | gttgtgggagtgtggaac | |
| dap-dR | gccggatcgaacgtga | |
| kap-dF | ctctctcgctttcctcat | |
| kap-dR | gctgatgacaaagccgt | |
| pUC18-k*TaSK-F | gacgcgtctcgtgcggacgaattcgtaatcatggtcatagctg | |
| pUC18-k*TaSK-R | gcggaccgtccgtcgaccgaattcactggccgtcgttttac | |
| pUC18-aTkk*-F | cggtgaactggacgtgtcgaattcgtaatcatggtcatagctg | |
| pUC18-aTk*k-R | atgaggaaagcgagagagaattcactggccgtcgttttac | |
| M13F | gtaaaacgacggccagt | |
| M13R | Caggaaacagctatgac | |
| ΔA-F | GGCTACGTCGTCACCGGTGTGCTGCCGCAGC TGAGCGAGGAACCTAGGtgcagctcacggtaactgat | *Avr*II site |
| ΔA-R | CTTGTAGACCGCCTCCTTCGCACAGAACAGC AGCCGGTCCCACCTAGGaggaacttatgagctcagcc | |
| bioAB-arm-up-F | ctcttcgcctaggaattcaagcttggatccatcacctccgtcccat | |
| bioAB-arm-up-R | gggtccatggacctgctgaacaccatcggcgtggttcag | |
| bioAB-arm-dw-F | gtggtccagctgaaccacgccgatggtgttcagcaggtccat | |
| bioAB-arm-dw-R | tgattacgaatttctagaccatgggacttcatcagcctggag | |
| aciB-arm-up-F | ctcttcgcctaggaattcaagcttggatccaactgtccgtctcactgt | |
| aciB-arm-up-R | ccagtagttgaagccgatctccgggcggtcgtagtcactgt | |
| aciB-arm-dw-F | ggggtcaacagtgactacgaccgcccggagatcggcttca | |
| aciB-arm-dw-R | tgattacgaatttctagaccatggctccagcggtacgagat | |
| pSUC01-sF | actacgaggtgctgagg | |
| pSUC01-sR | cggctcgtatgttgtgt | |
| pSUC01bioAB-sF1 | gaccaccaggaggacat | |
| pSUC01bioAB-sR1 | ggagccgctggtcat | |
| pSUC01bioAB-sF2 | tgtacggacggctcac | Sequencing primers |
| pSUC01bioAB-sR2 | gcgggcggaagcat | |
| pSUC01aciB-sF1 | gccacgatcaactcct | |
| pSUC01aciB-sR1 | gagatcagcgtcgggta | |
| pSUC01aciB-sF2 | gccgaacttcgccta | |
| pSUC01aciB-sR2 | gaactgctccagcgt | |
| ΔA-dF | ggagtgtggaacggact | |
| ΔA-dR | agcggtggcacaacat | |
| ΔbioAB-dF | ggaaggactccgccgaccag | Diagnostic primers |
| ΔbioAB-dR | gccgtgttgaccaggtgcct | |
| ΔaciB-dF | gccacgatcaactcct | |
| ΔaciB-dR | gaactgctccagcgt | |
| MV306op-F | tatcctgcaatcaagcttatcgatgtcg | |
| MV306op-R | cgttgcgctcggtcgttc | |

| | | |
|---|---|---|
| riboBlock-F | gaacgaccgagcgcaacg | |
| riboBlock-R | cgacatcgataagcttgattgcaggata | |
| bioD-Gib-F | ctaaggaggcaacaagatgacggtcctcgcggtcaccggc | |
| bioD-Gib-R | gtagtccgaaccacctccgctcaggctcgtgatccagtctg | |
| bioF-Gib-F | ctaaggaggcaacaagatgacgcgcgcaggtcttt | |
| bioF-Gib-R | gtagtccgaaccacctcctgcccgggccgtggcgagcacgtcggt<br>cag | |
| bioA-Gib-F | ctaaggaggcaacaagatggccgacctgaccccg | |
| bioA-Gib-R | gtagtccgaaccacctccggttaatgcacgcgcgacgc | |
| bioB-Gib-F | ctaaggaggcaacaagatgttcgaggtgttccgagatcttttc | |
| bioB-Gib-R | gtagtccgaaccacctcccagggtggcgttgagcgc | |
| riboOp-F | ggaggtggttcggactacaag | |
| riboOp-R | catcttgttgcctccttagcag | |
| 306-F | ccctgattctgtggataacc | |
| 306-R | gcctggcagtcgatcgta | |
| Synthesized gblocks | | |
| ADSK-gbk | *cctcccatggtataaatagtg****gcgccagcagctgatccgcaagcccgtccccaag***<br>***cccgctccccgagacctgg***ttta**aaccgcatccacgagaacaacaagttctgca**<br>**agggcaacagccgtctccccgacaaaa***atgtcgaaagctacatataagga* | *Pme*I site |
| ermEp*-svnN-svnA-fd terminator-gbk | cccatggtataaatagtggcggtaccagcccgacccgagcacgcgccggcacgcc<br>tggtcgatgtcggaccggagttcgaggtacgcggcttgcaggtccaggaaggggga<br>cgtccatgcgagtgtccgttcgagtggcggcttgcgcccgatgctagtcgcggttgat<br>cggcgatcgcaggtgcacgcggtcgatcttgacggctggcgagaggtgcgggga<br>ggatctgaccgacgcggtccacacgtggcaccgcgatgctgttgtgggcacaatcg<br>tgccggttggtaggatcgtctagaacaggaggccccatgtgatagacctgcggatcc<br>ttctcgtagacaactatgactcctacacgtggaacctcttccagctgatctggaaggtg<br>gcgggtgtgccgcccgtcgtcgtgcgcaacgacgagacgacggcggaagaactg<br>ctcggccaggacttcacccacgtcgtgatctccccgggcccgggcaccgcggcca<br>gggacgaggacttcgggctgtgccgggaactcctggagcgggccacggtgcccgt<br>cctcggagtgtgtctcggccaccagggcctggccctcgccttcggcggggacgtgc<br>gccacgcgccggagaccgtgcacgggcagaccagcgggatcacccacaccggg<br>accggcgtcttcgccggcatcccgcagggcttccgcgcggtgcgctaccactccct<br>caccgtcaccgagccgctgccggagcacctcgaggcgaccgcctggtccgagga<br>cggcgtcctgatgggcctgcgccaccgcgaacgccccctgcacggcgtgcagttc<br>catcccgaatcggtggacagcgagtacggcgaggagatcgtcgccaacttcctggg<br>catcacgcccgcgctccgctggagcgaaGTTTAAACgatcggggcgggcc<br>ttcctctcctcgacgcactggccctggatcggcgaacgcacccgctccgcccgtgg<br>cgcccacgtacgactgctcgccacggtggacaacccggtcgcctgcaaggtcggg<br>ccgacgatgaccccggcggagctgctggagctgtgcggcctgctggacccggag<br>cggcagcccgggcggctcaccctcatcgcccgccagggggccggcgcggtgcg<br>gcgcaaccttcccgcgctcgtggagtgcgtacgccgggccgggcatcccgtggtct<br>ggatgtgcgacccgatgcacggcaacaccgtcaccaccccggccggccgcaaga<br>cccgcctcgtcgaggaggtcgtccgcgaggtccggggcttccggcaggccgtgga<br>gggcgccggcgggaccgccgggggctgcacctggaggccacgccggacgag<br>gtgctggagtgcgtggccgacgccgacggcatcgccgggctcgacgggcggccc<br>agaccgctctgcgacccccggctgaacctggagcaggccgccacggccgtgtccg<br>cctggcgcatctgagcgccgccaaccacctctcaccagctgggatacaattaaagg<br>ctccttttggagcctttttttttggaatgtcgaaagctacatataa | *Pme*I site |
| kasOp*-T7 terminator-aac(3)IV-ΔSK-gbk | **ggtcgacggacggtccgcgcagagcgcggaccgtctggtcggcatgttggtcc**<br>**at**aactcccccagtcctgcacgctgtcgtattctcctggccacgactttacaacaccg<br>cacagcatgttgtcaaagcagagaccgttcgaatgtgaacaggatcccgcggatata<br>gttcctcctttcagcaaaaaacccctcaagacccgtttagaggccccaaggggttatg<br>ctagttattgctcagcggtggcagcaCATATGaggaacttatgagctcagccaa<br>tcgactggcgagcggcatcgcattcttcgcatcccgcctctggcggatgcaggaag<br>atcaacggatctcggcccagttgacccagggctgtcgccacaatgtcgcgggagcg | *Nde*I site |

| | | |
|---|---|---|
| | gatcaaccgagcaaaggcatgaccgactggaccttccttctgaaggctcttctccttg<br>agccacctgtccgccaaggcaaagcgctcacagcagtggtcattctcgagataatcg<br>acgcgtaccaacttgccatcctgaagaatggtgcagtgtctcggcaccccatággga<br>acctttgccatcaactcggcaagatgcagcgtcgtgttggcatcgtgtcccacgccga<br>ggagaagtacctgcccatcgagttcatggacacgggcgaccgggcttgcaggcga<br>gtgaggtggcagggggcaatggatcagagatgatctgctctgcctgtggccccgctg<br>ccgcaaaggcaaatggatgggcgctgcgctttacatttggcaggcgccagaatgtgt<br>cagagacaactccaaggtccggtgtaacgggcgacgtggcaggatcgaacggctc<br>gtcgtccagacctgaccacgagggcatgacgagcgtccctcccggacccagcgca<br>gcacgcagggcctcgatcagtccaagtggcccatcttcgagggggccggacgctac<br>ggaaggagctgtggaccagcagcacaccgccgggggtaaccccaaggttgagaa<br>gctgaccgatgagctcggcttttcgccattcgtattgcacgacattgcactccaccgct<br>aatgacatcagtcgatcatagcacgatcaacggcactgttgcaaatagtcggtggtga<br>taaacttatcatccccttttgcttatggagctgcacatgaacccattcaaaggccggcat<br>tttcagcgtgacatcattctgtgggccgtacgctggtactgcaaatacggcatcagtta<br>ccgtgagctgcaCATATG**tcgacggtaccaagtcgccatgatcacgctgaa**<br>**taccagccggggaagaaaccctgtgagacgcgtctcgtgcggac** | |
| aac(3)IV-T7 terminator-<br>kasOp*-kap-gbk | **ctctctcgctttcctcattcggcgctccccttccgcgtctcattggtcttccgctccg**<br>**tttttgctttgccgatgttacttggggagaggtgcgataatcctttcgcaaaaact**<br>**cggtttgacgcctcccatggtataaatagtggc**CTCGAGtgcagctcacggt<br>aactgatgccgtatttgcagtaccagcgtacggcccacagaatgatgtcacgctgaa<br>aatgccggcctttgaatgggttcatgtgcagctccataagcaaaaggggatgataagt<br>ttatcaccaccgactatttgcaacagtgccgttgatcgtgctatgatcgactgatgtcatt<br>agcggtggagtgcaatgtcgtgcaatacgaatggcgaaaagccgagctcatcggtc<br>agcttctcaaccttggggttaccccccggcggtgtgctgctggtccacagctccttccgt<br>agcgtccggcccctcgaagatgggccacttggactgatcgaggccctgcgtgctgc<br>gctgggtccgggagggacgctcgtcatgccctcgtggtcaggtctggacgacgag<br>ccgttcgatcctgccacgtcgcccgttacaccggaccttggagttgtctctgacacatt<br>ctggcgcctgccaaatgtaaagcgcagcgcccatccatttgcctttgcggcagcggg<br>gccacaggcagagcagatcatctctgatccattgcccctgccacctcactcgcctgc<br>aagcccggtcgcccgtgtccatgaactcgatgggcaggtacttctcctcggcgtggg<br>acacgatgccaacacgacgctgcatcttgccgagttgatggcaaaggttccctatgg<br>ggtgccgagacactgcaccattcttcaggatggcaagttggtacgcgtcgattatctc<br>gagaatgaccactgctgtgagcgctttgccttggcggacaggtggctcaaggagaa<br>gagccttcagaaggaaggtccagtcggtcatgcctttgctcggttgatccgctcccgc<br>gacattgtggcgacagccctgggtcaactgggccgagatccgttgatcttcctgcatc<br>cgccagaggcgggatgcgaagaatgcgatgccgctcgccagtcgattggctgagc<br>tcataagttcctCTCGAGtgctgccaccgctgagcaataactagcataacccctt<br>ggggcctctaaacgggtcttgaggggttttttgctgaaaggaggaactatatccgcgg<br>gatcctgttcacattcgaacggtctctgctttgacaacatgctgtgcggtgttgtaaagt<br>cgtggccaggagaatacgacagcgtgcaggactgggggagtt**atgccccacaag**<br>**gtgattctggcgcttctcgcaatcggctcctttgccgtcggcacggacggctttgt**<br>**catcagcgggattcttccgcgcatcgccggtgaactggacgtgtc** | *Xho*I site |
| A37 promoter-<br>**riboswitch/aptamer-**<br>**RBS**-*start codon*-dual<br>direction typeIIS sites-<br>***Linker + 3xFLAG tag*** | gaacgaccgagcgcaacgctagagattgcgaaggttcaatccaccgacgtacactg<br>tactcggatccaattgcaatgatccatacgactcactataggtacc**ggtgataccagc**<br>**atcgtcttgatgcccttggcagcacc**ctgct**aaggagg**caacaag*atg*agaaga<br>gctcttct***ggaggtggttcggactacaaggatgatgatgacaaggactacaaaga***<br>***tgatgatgataaagattataaagacgatgacgataaatga***tatcctgcaatcaagc<br>ttatcgatgtcg | riboBlock<br>cassette |

**Table S9. Plasmids used in this study.**

| Plasmids | Description | Source |
|---|---|---|
| pUC18 | *bla rep(pMB1) lacZ*, general cloning vector | Addgene |
| pCGW | *aac(3)IV ura3 cen/ARS trp1 sopABC repE ori2 oriV cat neo traJ-oriT attP-int*$_{\Phi C31}$, TAR cloning vector | Lab stock |
| pR9406 | *bla*, pUB307 derived *E. coli*-*Streptomyces* tri-parental conjugation helper plasmid | [1] |
| pKD46 | *gam bet exo*, λ-red recombination plasmid | [2] |
| pADSK-cap | pCGW inserted with ADSK-gbk between *Nde*I/*Xho*I sites, ADSK BGC capture plasmid | This study |
| pSvn-cap | pCGW inserted with ermEp*-svnN-svnA-fd terminator-gbk between *Nde*I/*Xho*I sites, svn BGC capture plasmid | This study |
| pADSK | pCGW carrying ADSK BGC | This study |
| pSvn-ermEp* | pCGW carrying svn BGC driven by ermEp* promoter | This study |
| pUC18-k*TaSK | kasOp*-T7 terminator-aac(3)IV-ΔSK-gbk inserted into pUC18 vector between *Hind*III/*EcoR*I sites | This study |
| pUC18-aTk*k | aac(3)IV-T7 terminator-kasOp*-kap-gbk inserted into pUC18 vector between *Hind*III/*EcoR*I sites | This study |
| pADSKΔsvn::aac(3)IV | pADSK Δ*svnA-svnN*:: *aac(3)IV* | This study |
| pADSKΔsvn | pADSK Δ*svnA-svnN* | This study |
| pADSKΔsvn2::aac(3)IV | pADSK Δ*aciD-svnN*::*aac(3)IV* | This study |
| pADSKΔsvn2 | pADSK Δ*aciD-svnN* | This study |
| pADSKΔsvn3::aac(3)IV | pADSK Δ*svnA-kapF*::*aac(3)IV* | This study |
| pADSKΔsvn3 | pADSK Δ*svnA-kapF* | This study |
| pADSKΔDS::aac(3)IV | pADSK Δ*dapT-svnA*::*aac(3)IV* | This study |
| pADSKΔDS | pADSK Δ*dapT-svnA* | This study |
| pADSKΔSK::aac(3)IV-kasOp* | pADSK Δ *svnA-kapF*::kasOp*-T7 terminator-*aac(3)IV* | This study |
| pAD-kasOp* | pADSK Δ*svnA-kapF*::kasOp*-T7 terminator | This study |
| pAD-kasOp*ΔA::aac(3)IV | pADSK Δ*svnA-kapF*::kasOp*-T7 terminator Δ*aciT-aciE*::*aac(3)IV* | This study |
| pDap-kasOp* | pADSK Δ*svnA-kapF*::kasOp*-T7 terminator Δ*aciT-aciE* | This study |
| pADSKΔDSK::aac(3)IV | pADSK Δ*dapT-kapF*::*aac(3)IV* | This study |
| pAci | pADSK Δ*dapT-kapF* | This study |
| pAci::aac(3)IV-kasOp* | pADSK Δ*dapT-kapF* Δ12121-12222::*aac(3)IV*-T7 terminator-kasOp*[a] | This study |

| | | |
|---|---|---|
| pAci-kasOp* | pADSK Δ*dapT-kapF* Δ12121-12222::T7 terminator-kasOp* | This study |
| pADSKΔADS::aac(3)IV-kasOp* | pADSK Δ*orf-9-svnA*::*aac(3)IV*-T7 terminator-kasOp*[b] | This study |
| pKap-kasOp* | pADSK Δ*orf-9-svnA*::T7 terminator-kasOp* | This study |
| pSUC01 | *aac(3)IV traJ-oriT kasOp*-SaindC* $ori_{colE1}$, suicide plasmid for homologous recombination-mediated gene deletion in *Streptomyces* | [3] |
| pDSK | pADSK Δ*aciT-aciE* | This study |
| pSUC01-bioAB | *bioAB* deletion plasmid | This study |
| pSUC01-aciB | *aciB* deletion plasmid | This study |

a, Intergenic region between *avd* and *aciT* spanning from 12121 bp to 12222 bp on pAci plasmid was deleted and replaced with *aac(3)IV*-T7 terminator-kasOp* cassette.

b, 9 genes upstream of *avd* (*orf-9*) to *svnA* region on pADSK was replaced with *aac(3)IV*-T7 terminator-kasOp* cassette.

**Table S10. Strains used in this study.**

| Strains | Description | Source |
|---|---|---|
| *E. coli* Top10 | *F⁻ mcrA Δ(mrr-hsdRMS-mcrBC) φ80lacZΔM15 ΔlacX74 nupG recA1 araD139 Δ(ara-leu)7697 galE15 galK16 rpsL(*$Str^R$*) endA1 λ⁻*, general cloning strain | Invitrogen |
| *E. coli* TransforMax EPI300 | *F⁻ mcrA Δ(mrr-hsdRMS-mcrBC) φ80dlacZΔM15 ΔlacX74 recA1 endA1 araD139 Δ(ara, leu)7697 galU galK λ⁻ rpsL (*$Str^R$*) nupG trfA tonA*, host strain for inducible propagation of pCGW-derived plasmids | Lucigen |
| *E. coli* ET12567 | *F⁻ dam-13::Tn9 dcm-6 hsdM hsdR zjj-202::Tn10 recF143 galK2 galT22 ara-14 lacY1 xyl-5 leuB6 thi-1 tonA31 rpsL136 hisG4 tsx-78 mtl-1 glnV44*, *E. coli* host strain for *E. coli-Streptomyces* mating | [4] |
| *E. coli* BW25113 | Wild-type strain<br>$lacI^+ rrnB_{T14}$ $\Delta lacZ_{WJ16}$ *hsdR514* $\Delta araBAD_{AH33}$ $\Delta rhaBAD_{LD78}$ *rph-1* Δ(*araB–D*)*567* Δ(*rhaD–B*)*568* Δ*lacZ4787*(::*rrnB-3*) *hsdR514 rph-1*, λ-red PCR targeting host strain | CGSC<br>Lab stock |
| *E. coli* BW25113 | Δ*dppA*, Δ*oppA*, Δ*dtpA* | [5] |
| *E. coli* WCC C0244 | | IIDR clinical isolate collection |
| *S. cerevisiae* VL6-48N | *MAT α, his3-D200, trp1-Δ1, ura3-Δ1, lys2, ade2−101, met14, psi+*$cir^O$, host strain for TAR cloning | [6] |
| *S.* sp. WAC05950 | Parental strain producing stravidins | Wright Actinomycete Collection |
| *S. coelicolor* M1154 | Δ*act* Δ*red* Δ*cpk* Δ*cda rpoB[C1298T] rpsL[A262G]*, host strain for heterologous expression | [6, 7] |
| *S. coelicolor* M1154/pADSK | pADSK integrated into the chromosome of *S. coelicolor* M1154 at $attB_{\varphi C31}$ site | This study |
| *S. coelicolor* M1154/pSvn-ermEp* | pSvn-ermEp* integrated into the chromosome of *S. coelicolor* M1154 at $attB_{\varphi C31}$ site | This study |
| *S. coelicolor* M1154/pADSKΔsvn | pADSKΔsvn integrated into the chromosome of *S. coelicolor* M1154 at $attB_{\varphi C31}$ site | This study |

| | | |
|---|---|---|
| *S. coelicolor* M1154/pADSKΔsvn2 | pADSKΔsvn2 integrated into the chromosome of *S. coelicolor* M1154 at attB$_{φC31}$ site | This study |
| *S. coelicolor* M1154/pADSKΔsvn3 | pADSKΔsvn3 integrated into the chromosome of *S. coelicolor* M1154 at attB$_{φC31}$ site | This study |
| *S. coelicolor* M1154/pADSKΔDS | pADSKΔDS integrated into the chromosome of *S. coelicolor* M1154 at attB$_{φC31}$ site | This study |
| *S. coelicolor* M1154/pAD-kasOp* | pAD-kasOp* integrated into the chromosome of *S. coelicolor* M1154 at attB$_{φC31}$ site | This study |
| *S. coelicolor* M1154/pDap-kasOp* | pDap-kasOp* integrated into the chromosome of *S. coelicolor* M1154 at attB$_{φC31}$ site | This study |
| *S. coelicolor* M1154/pAci | pAci integrated into the chromosome of *S. coelicolor* M1154 at attB$_{φC31}$ site | This study |
| *S. coelicolor* M1154/pAci-kasOp* | pAci-kasOp* integrated into the chromosome of *S. coelicolor* M1154 at attB$_{φC31}$ site | This study |
| *S. coelicolor* M1154/pKap-kasOp* | pKap-kasOp* integrated into the chromosome of *S. coelicolor* M1154 at attB$_{φC31}$ site | This study |
| *S. coelicolor* M1154/pDSK | pDSK integrated into the chromosome of *S. coelicolor* M1154 at attB$_{φC31}$ site | This study |
| WAC05950Δ*bioAB* | *bioAB* deletion mutant of WAC05950 | This study |
| WAC05950Δ*aciB* | *bioAB* deletion mutant of WAC05950 | This study |
| *K. pneumoniae* MKP103 | Wild-type strain | [8] |
| *K. pneumoniae* ATCC 43816 | Wild-type strain | ATCC |
| *A. baumannii* ATCC 17978 | Wild-type strain | ATCC |
| *P. aeruginosa* PA01 | Wild-type strain | Lab stock |
| *M. smegmatis* mc$^2$155 | Wild-type strain | Lab stock |
| *M. fortuitum* ATCC6891 | Wild-type strain | ATCC |
| *M. abscessus* ATCC19977 | Wild-type strain | ATCC |
| *M. tuberculosis* H37Ra | Wild-type strain | Lab stock |
| *M. bovis* BCG Pasteur ATCC35734 | Wild-type strain | ATCC |

**Table S11. Plasmid and strain information for the recombinant expression of *E. coli* BioA, BioD, BirA and AccB.**

| Protein | Plasmid Construct | Tag | Tag Position | Antibiotic Resistance | Expression Strain |
|---|---|---|---|---|---|
| BioA | pCA24N-bioA | His-tag | N-terminal | Chloramphenicol | *E. coli* K12 AG1-pCA24N-bioA (JW0757) |
| BioD | pET-21a(+)-bioD | His-tag | C-terminal | Ampicillin | *E. coli* BL21 Star (DE3) pLysS (Invitrogen) |
| BirA | pET-28a(+)-sumo-birA | His-SUMO-tag | N-terminal | Kanamycin | *E. coli* K12 SHuffle T7 (NEB) |
| AccB | pET151/D-TOPO-accB | His-tag | C-terminal | Ampicillin | *E. coli* BL21 Star (DE3; Invitrogen) |

**Table S12. Protein purification buffer components.**

| Protein | Buffer A | Buffer W | Buffer E | Storage Buffer |
|---|---|---|---|---|
| BioD | 50 mM HEPES, 300 mM NaCl, 10 mM imidazole, pH 8 | Buffer A, 20 mM imidazole | Buffer A, 250 mM imidazole | 50 mM HEPES pH 8, 10% Glycerol |
| BirA | 25 mM Tris-HCl, 200 mM NaCl, 10 mM imidazole, pH 8 | Buffer A, 20 mM imidazole | Buffer A, 300 mM imidazole | 25 mM Tris-HCl, pH 8, 10% Glycerol |
| AccB | 50 mM $NaH_2PO_4$, 500 mM NaCl, 10 mM imidazole, pH 8 | Buffer A, 25 mM imidazole | Buffer A, 250 mM imidazole | 50 mM $NaH_2PO_4$, pH 8, 20% Glycerol |
